# Different molecular recognition by three domains of the full-length GRB2 to SOS1 proline-rich motifs and EGFR phosphorylated sites

**DOI:** 10.1101/2024.04.20.590390

**Authors:** Keita Tateno, Takami Ando, Maako Tabata, Haruka Sugasawa, Toshifumi Hayashi, PM Sayeesh, Kohsuke Inomata, Tsutomu Mikawa, Yutaka Ito, Teppei Ikeya

## Abstract

The adaptor protein human GRB2 plays crucial roles in mediating signal transduction from cell membrane receptors to RAS and its downstream proteins by recruiting SOS1. Recent studies have revealed that GRB2 also serves as a scaffold for liquid-liquid phase separation (LLPS) with SOS1 and transmembrane receptors, which is thought to regulate the magnitude of cell signalling pathways. In this study, we employed solution NMR spectroscopy to investigate the interactions of the full-length GRB2 with proline-rich motifs (PRMs) derived from ten potential GRB2-binding sites in SOS1, as well as a peptide from a phosphorylation site of EGFR. Our findings indicate that the binding affinity of the two SH3 domains of GRB2 for PRMs differs by a factor of ten to twenty, with the N-terminal SH3 domain (NSH3) exhibiting a markedly higher affinity. The interactions of PRMs with the SH3 domains affected not only the regions surrounding the PRM binding sites on the SH3 domains but also the linker area connecting the three domains and parts of the SH2 domain. Analysis of the interaction between the phosphorylated EGFR binding site and the SH2 domain revealed chemical shift perturbations in regions distal from the known binding site of SH2. Moreover, we observed that the inter-domain interactions of the two SH3 domains with the SH2 domain of GRB2 are asymmetric. These findings suggest that the local binding of PRMs and phosphorylated EGFR to GRB2 impacts the overall structure of the GRB2 molecule, including domain orientation and dimerisation, which may contribute to LLPS formation.

## Introduction

The RAS-mediated signal pathways play crucial functions in intracellular events, such as cell proliferation, differentiation, and survival^1,2^. This cascade is initiated by the binding of extracellular ligands with receptor Tyr kinases (RTKs) on the cell membrane, and then transmits the signal to its downstream proteins. The ligand binding stimulates trans-autophosphorylation of tyrosine in the intracellular regions of RTKs, recruiting growth factor receptor-bound protein 2 (GRB2) along with Son of sevenless homologue 1 (SOS1) near the inner-membrane surface. This event allows SOS1 to encounter RAS anchored on the transmembrane and activate RAS by exchanging the bound GDP for GTP. The activated RAS then propagates the signal to the cytosol and nucleus through further downstream proteins.

GRB2 is a key mediator in this upstream process, comprising a Src homology 2 (SH2) domain flanked by N-terminal and C-terminal Src homology 3 (SH3) domains (Fig. S1a). While the SH2 domain associates with the phosphorylated tyrosine of RTKs, the N-terminal and C-terminal SH3 domains (henceforth referred to as NSH3 and CSH3, respectively) interact with proline-rich motifs (PRMs) of SOS1. SOS1, serving as a guanine-nucleotide-exchange factor (GEF) of RAS and RAC, is also a multi-domain protein composed of six major domains or motifs, a Histone Fold (HF), Dbl homology (DH), Pleckstrin homology (PH), RAS exchange motif (REM), cell division cycle 25 (CDC25), and proline-rich (PR) (Fig. S1b). The PR domain, which associates with the SH3 domains of GRB2 through PRMs, is an intrinsically disordered region (IDR) as an extended random coil-like conformation in solution.

In addition to the role of GRB2 as a carrier of SOS1 to the membrane, recent studies have shown that GRB2 also serves as a major component for liquid-liquid phase separation (LLPS) in the RAS-mediated transduction pathway by acting as a scaffold for other proteins^3, 4^. Several experiments indicate that the association and dissociation dynamics of LLPS can regulate the magnitude of the signal transduction on the RAS-mediated signal pathways^5^. LLPS involving GRB2 was initially discovered in the system with GRB2, SOS1, and linker for activation of T cells (LAT), and has recently been found in a system with GRB2 and EGFR^6^. Although the precise molecular mechanisms of phase separation in these systems are not fully understood, it is thought that the three domains of GRB2 work together to bundle the other proteins within these complexes. In the ternary complex composed of GRB2, SOS1 and LAT, it is proposed that GRB2 serves as a bridge between LAT and SOS1, owing to the association of its SH2 domain with phosphorylated tyrosine residues on LAT and two SH3 domains with PRMs on SOS1, respectively (Fig. S1c). In the binary complex formation of GRB2 and EGFR, the dimerisation of GRB2 is thought to be a key for the connection with EGFR^7^. However, sufficient experimental evidence to substantiate these models remains lacking.

Hence, it is crucial to elucidate the interactions of GRB2 with SOS1 at atomic resolution for a detailed understanding of the regulatory mechanisms in the signal transduction pathway. While the 3-dimensional (3D) structure of GRB2 is determined by X-ray crystallography^8^, other structural and computational studies, such as small angle X-ray scattering (SAXS)^9^, molecular dynamics simulation ^10^, and protein structure predictions via AlphaFold 2 (UniProt Accession ID: P62993), suggest that the relative orientation of the three domains connected with two flexible loops significantly differs from the crystal structure (Fig. S1d) in aqueous solutions. The domain orientation of GRB2 appears to be dynamic even in the condensed condition, allowing for flexible interactions with SOS1, LAT or EGFR. Considering that the PR domain of SOS1 is also extraordinarily flexible, or intrinsically disordered region (IDR), the molecular interaction of GRB2 and SOS1 could be highly dynamic, leading to an enormous number of different conformations in the condensed droplet. Optical microscope experiments have shown that these associations and dissociations are also highly dynamic on a time scale of seconds to minutes^3, 5, 6^. Yet, it remains unclear how the microscopic molecular motions of GRB2 and SOS1 on a time scale of μsec to msec result in macroscopic droplet formation at a much slower time rate. It is therefore requisite to investigate the 3D structures and dynamics of these proteins in solution at atomic resolution. It still remains a challenge to observe it by generally used structural analysis methods, such as X-ray crystallography and cryo-electron microscopy (cryo-EM). Thus, we employed solution Nuclear Magnetic Resonance (NMR) spectroscopy which is an optimal tool for studying highly dynamical proteins, including multi-domain proteins with flexible linkers and IDRs in aqueous or condensed environments.

Liao et al. recently reported interactions of isolated NSH3 and CSH3 domains of GRB2 with several PR peptides derived from potential PRMs of SOS1 by solution NMR^10^. This work proposed that the SOS1 PR domain possesses ten potential binding sites for the SH3 domains of GRB2 (henceforth referred to as S1-S10 from the N-terminus of the SOS1-PR domain; Fig. S1b). This report also suggested that nine of the ten sites preferentially bind to the NSH3 domain, while one site has a higher affinity for CSH3. However, these data were obtained using the isolated NSH3 and CSH3 domains, not the full-length GRB2. It is assumed that if the two SH3 domains of GRB2 bind to a single binding site in SOS1, competition between the two SH3 domains for the single SOS1 binding site would occur, resulting in different binding affinities and modes compared to independent bindings. Furthermore, the two linkers connecting the SH3 domains are highly flexible, and their relative orientations can significantly change, which may affect their respective interactions with SOS1. In fact, significant differences in binding affinity have been reported between a single domain and a full-length protein^11^, highlighting the need for careful investigation of the binding sites of the full-length GRB2 and SOS1 PRMs.

Hence, in this work, we elucidate the molecular recognition mechanisms and binding affinities of the full-length GRB2 with several SOS1 PRMs and an EGFR phosphorylated peptide. Through NMR titration as well as isothermal titration calorimetry (ITC), we carefully identify the binding sites and modes, and their affinities for each SOS1 PRM by considering several theoretical binding models. In addition, docking simulations based on the experimental data provide insights into the interaction mechanisms based on tertiary structures.

## Results

### Backbone and side-chain resonance assignments

Yuzawa et al. show that binding of GRB2 with two peptides, EpYINSQV derived from EGFR (henceforth referred to as EGFR peptide) and VPPPVPPRRR from SOS1 (1149-1158; henceforth referred to as SOS1 S4 PRM), stabilise GRB2 in a monomeric state at 25 °C in 20 mM potassium phosphate buffer (pH 7.2) ^9^. This report also mentions that a double mutant of GRB2 [C32S, C198A] exhibits higher stability while having the same structure as the wild-type. Thus, we principally employed the double mutant GRB2 with the EGFR and SOS1-derived peptides for most of NMR analyses, except for titration experiments with other SOS1 PRMs. Although the backbone ^1^H^N^, ^15^N, ^13^C^α^, and ^13^C′, and ^13^C^β^ resonances of GRB2 with and without the EGFR peptide and SOS1 S4 have been assigned in the previous studies^9, 12^, the resonance assignments for side-chains and portions of the backbones have not been fully analysed. Thus, we initially assigned their resonances using three-dimensional (3D) triple-resonance spectra with ^13^C,^15^N-uniformly and ^2^H,^13^C,^15^N-uniformly isotope-labelled protein samples as well as the samples with amino acid-selective isotope labelling techniques to resolve highly overlapped signals. The amino acid-selective isotope labelling (lysine and leucine selectively labelled GRB2) and TROSY-type NMR experiments enabled us to correct some erroneous resonance assignments by the previous reports (Fig. S2). Overall, we achieved 89.2 % and 62.3 % of accurate backbone (^1^H^N^, ^15^N, ^13^C^α^, and ^13^C′) and all atom resonance assignments, respectively, for the full-length GRB2 in the presence of the EGFR peptide and SOS1 S4 (Fig. S3 and S4). Despite measuring and carefully analysing TROSY-type spectra with predeuterated samples and spectra with three amino acid-selective labelled samples,, the resonance assignments in some regions, particularly the two linkers connecting each domain, remain incomplete. This suggests that these linker regions may possess significant dynamics or exist in multiple conformers (Fig. S3).

### Comparison of NMR spectra between the full-length GRB2 and each isolated domain

To confirm the spatial relationships among the three domains, or inter-domain interactions, we independently measured 2D ^1^H-^15^N HSQC spectra of isolated NSH3, SH2 and CSH3 domains in addition to the full-length GRB2 (Fig. S5). A comparison of these spectra revealed that peaks corresponding to NSH3 and SH2 domains did not overlap well with those in the full-length GRB2. In contrast, the CSH3 spectrum showed strong overlap with its corresponding peaks in the full-length GRB2. This divergence in the spectra of NSH3 and SH2 from that of the full-length GRB2 could be due to extensive contacts between NSH3 and SH2 within the full-length protein, causing them to behave as a single unit and affecting the chemical shifts of individual atoms. On the other hand, the strong overlap between the isolated CSH3 and the full-length GRB2 suggests that CSH3 independently exists from the other two domains most of the time. These observations are consistent with previous NMR relaxation and small angle X-ray scattering (SAXS) studies^9^. The NMR relaxation data show that the correlation times *τ*_e_ for residues in CSH3 were smaller than those in the other regions, which was supported by the SAXS profile that each domain is more widely distributed. Notably, these data contrast with the X-ray crystal structure (Fig. S1d), where both SH3 domains are adjacent to each other but relatively distant from the SH2 domain.

### NMR titration experiments with different PRMs derived from SOS1

It is known that the GRB2-NSH3 domain associates with a proline-rich motif (PxxPxR) derived from the SOS1 PR domain, while CSH3 interacts more with another proline-rich motif (PxxxRxxKP) from GRB2-associated-binding protein 2 (GAB2)^13, 14^. Liao et al proposed that the SOS1 PR domain contains ten PxxPxR motifs but not the PxxxRxxKP motif^10^. To elucidate the binding modes of the full-length GRB2 towards the SOS1 PRMs, we performed NMR titration experiments of ^15^N-labelled full-length GRB2 samples. Based on the previous report by Liao et al., we selected and analysed four PRMs (S4, S5, S9 and S10) among ten motifs through NMR titration experiments. S4 (1149— 1158; VPPPVPPRRR), S5 (1177 — 1188; DSPPAIPPRQPT), and S9 (1287 — 1298. IAGPPVPPRQST), which contain the PxxPxR motif, exhibited relatively high affinities against isolated NSH3 domain. Conversely, S10 (1303—1314; PKLPPKTYKREH) is the only PRM that was reported to bind more strongly to the CSH3 domain than the NSH3 domain. The remaining PRMs as well as the above four were analysed by ITC (Fig. S15).

2D ^1^H-^15^N HSQC spectra of GRB2 in the presence of S4, S5, S9, and S10 PRMs with various protein-PRM concentration ratios are shown in Figures 1 and S6-8. Chemical shift perturbations (CSPs) of the backbone ^1^H and ^15^N resonances during the titration with the SOS1 PRMs are plotted against the amino sequence (Fig. 2). This figure presents CSPs at each titration point with different colours from blue to red, demonstrating that, for SOS1 S4, S5, and S9 PRMs, the NSH3 bars predominantly show blue colours, whereas the CSH3 shades are mostly orange or yellow. This indicates that the CSPs in NSH3 occurred at early stages in the titration experiments, whereas those in CSH3 were seen even at its later points. This observation supports the previous report, utilising isolated NSH3 and CSH3 domains, that GRB2 possesses two distinct binding affinities, with NSH3 exhibiting higher affinities for S4, S5, and S9 PRMs compared to CSH3^10^. Figure S9 further illustrates this trend, showing NSH3 undergoes early CSP changes, while CSH3 shifts occur later in the titration. Additionally, our data using the full-length GRB2 reveals that the CSPs are also observed in some residues of the SH2 domain, occurring at later stages in the titration compared to NSH3, thus aligning or presumably synchronising with the CSP patterns of CSH3 (Figure 2 and S9). Moreover, the previous study with the isolated SH3 domains proposed that the binding affinity of SOS1 S10 PRM was higher for CSH3 rather than for NSH3. However, our titration experiments with SOS1 S10 PRM do not show a clear difference in CSPs between NSH3 and CSH3, indicating that the CSP patterns for S10 PRM are clearly different from the others PRMs probably due to its unique properties.

**Figure 1.**
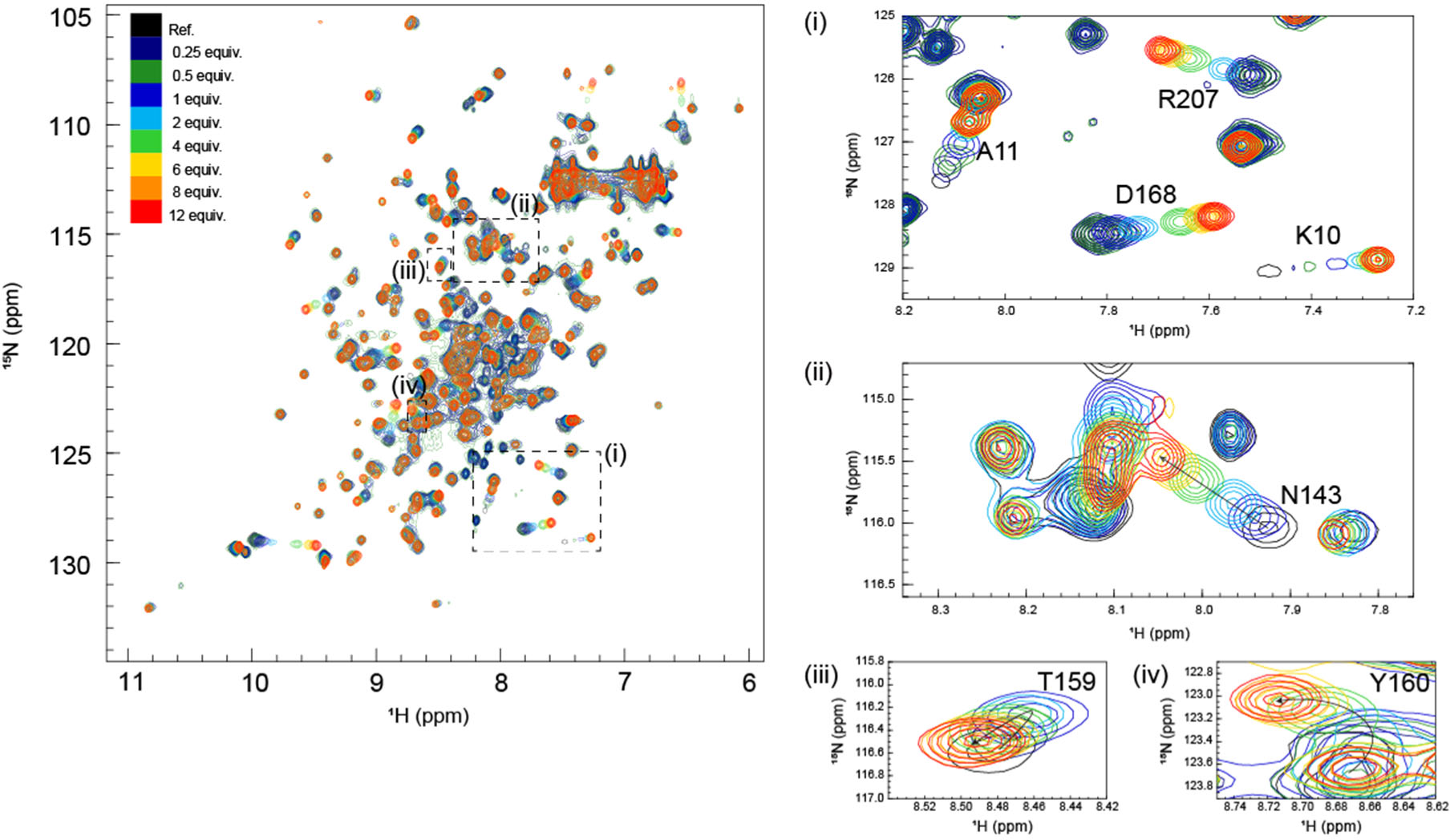
Overlays of 2D ^1^H-^15^N HSQC spectra from multipoint titrations of ^15^N-labelled GRB2 with SOS1-S4 PRM (VPPPVPPRRR). The PRM concentration was increased stepwise (the protein: PRM molar ratio of 1:0.25, 1:0.5, 1:1, 1:2, 1:4, 1:6, 1:8, and 1:12). In this figure, the colour codes of ^1^H-^15^N correlation cross-peaks at each titration point, showing the molar ratio of GRB2: SOS1-S4, are as follows: black (1:0); dark blue (1:0.25); dark green (1:0.5); blue (1:1); cyan (1:2); green (1:4); yellow (1:6); orange (1:8); red (1:12). Cross peaks that showed large chemical shift changes were annotated.

**Figure 2.**
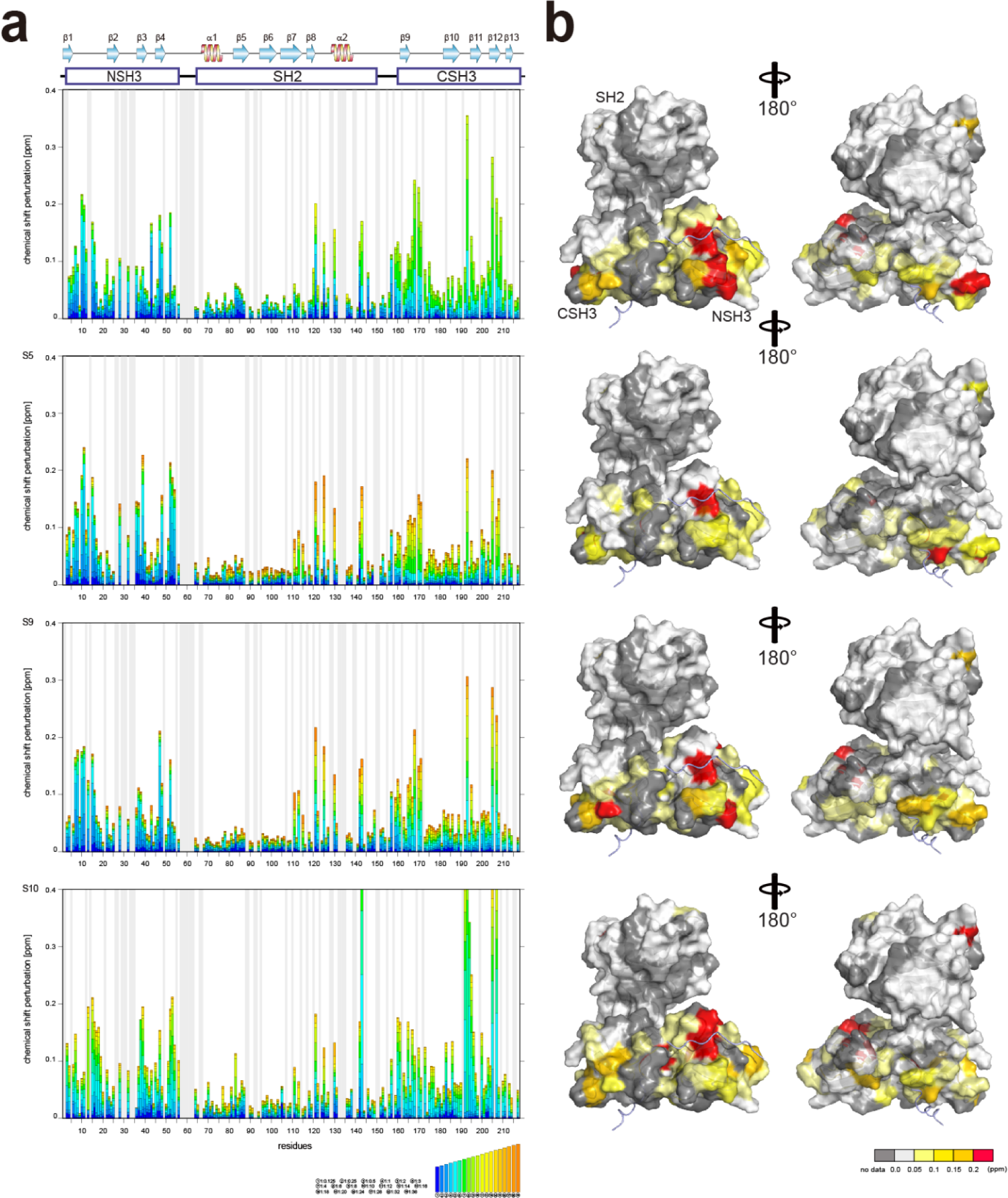
Plots of chemical shift perturbation of backbone ^1^H^N^ and ^15^N nuclei of GRB2 upon the titration with S4, S5, S9, and S10 PRMs. (a) Bar plots of chemical shift perturbation of backbone ^1^H^N^ and ^15^N nuclei of GRB2 upon the titration with S4, S5, S9, and S10 PRMs. The mean shift difference Δδ_ave_ was calculated as [(Δδ^1^H^N^)^2^ + (Δδ^15^N/5)^2^]^1/2^ where Δδ^1^H^N^ and Δδ^15^N are the chemical shift differences (ppm) between GRB2 on its own and in the presence of the PRMs. The bar graphs are colour-coded according to the protein-peptide concentration ratio and are overlaid. The proline residues as well as the residues for which ^1^H^N^-^15^N correlation cross peaks were not analysed due to signal overlap or other reasons are shown in grey. The secondary structures of GRB2 are also shown. (b) Chemical shift perturbation upon the titration with S4, S5, S9, and S10 PRMs represented on the crystal structure of GRB2 (PDB ID: 1GRI).

Significant CSPs observed upon PRM binding were mapped onto the crystal structure of GRB2 (Fig. 2b). As expected, for all PRMs, notable CSPs were predominantly located within the RT loops of both SH3 domains and the adjacent β-strands (β2, 3, 11, and 12). Meanwhile, as described above, significant CSPs were also identified at residues 120-145 in the SH2 domain, which is not considered to have PRM binding sites. The CSPs of N143 in the SH2 domain were especially pronounced, with the S10 PRM inducing the most significant change (Fig. 1 and S6-S8). The spatial separation of the SH3 and SH2 domains in the crystal structure (Fig. S1d) implies interactions beyond the direct binding sites of SH3. Furthermore, substantial CSPs were detected in the linker region connecting SH2 and CSH3 but not in the linker between NSH3 and SH2. Near CSH3 in the linker to SH2, non-linear trajectories, such as curve or U-turn patterns, were observed at residues Q157, T159, and Y160 (Fig. 1 and S6-S8). Three possible explanations can be considered for this observation: (1) direct but weak interactions of the PRMs with SH2, (2) PRM-binding to the SH3 domains altering their relative domain locations by affecting the linker and the adjacent areas of the SH2 domain, and (3) GRB2 dimerisation influencing the SH2 domain and linker. Considering the unique U-turn shaped perturbations in the linker between SH2 and CSH3, the CSPs synchronised between CSH3 and SH2, as well as the independently dynamic behaviour of CSH3, these potentially indicate a shift in the overall domain orientation of GRB2 upon PRM bindings to the SH3 domains. Subsequently, this domain orientation may also cause GRB2 dimerisation. More interestingly, exclusively S10 shows a linear chemical shift change but not U-turn patterns in this region (residues 157-160), similar to those in the other residues, also suggesting that S10 has a characteristic binding mode from the others. These CSPs in the SH2 domain and the linker regions, other than the SH3 domains, were not previously considered and were revealed for the first time by analysis using full-length GRB2.

Comparing the magnitude of the CSPs in the titration experiments, the overall shift changes were more significant for CSH3 than for NSH3.

### Dissociation constants of the SOS1-derived PR peptides calculated from NMR chemical shift perturbation

Arai et al. proposed a model with two-site independent binding with and without interconversion between the binding sites^15^, while Yagi et al. assumed that affinities of substrates to two different binding sites are not independent but cooperative with each other to explain a catalytic reaction^16^. For the two SH3 domains of GRB2 interacting with the SOS1 PRMs, despite each domain having a binding site, their proximity in the crystal structure (Fig. S1d) implies that binding at one site could influence the other. Consequently, we initially evaluated both a two-site independent model and a two-site cooperative binding model, conforming their reliabilities against CSP data via bootstrap resampling (Fig. 3a and S10). Models with two binding sites basically require a total of four dissociation constants (*K*_D1_–*K*_D4_). The two-site independent binding model equates *K*_D3_ and *K*_D4_ with *K*_D2_ and *K*_D1_, respectively. In contrast, the two-site cooperative binding model (two-site dependent model) differentiates the four dissociation constants, with the assumption that *K*_D3_ is modulated by another ligand binding to *K*_D2_, as a function of *K*_D2_ and a modulation factor *α*. *K*_D4_ is then calculated from *K*_D1_ and *K*_D2_, as the equation provided in the figure (The detailed models and theory are described in materials & methods). To determine these dissociation constants concerning the SOS1 S4, S5, S9, and S10 PRMs, the spectra of GRB2 from a single titration series were fitted to the two two-site binding models by simulating the CSPs along with the PRM additions using non-linear least-squares fitting. The bootstrap analysis with 1000 iterations demonstrated that the residual errors of the two-site independent binding models were sufficiently small (Fig. S10a). This led us to conclude that the two-site independent binding model provides a sufficient explanation for the global dissociation constants of the entire molecule, suggesting minimal interference between the two binding sites at least in terms of chemical shift perturbation. Thus, we computed exclusively *K*_D1_ and *K*_D2_ utilising the two-site independent binding model (Fig. S11-S14). Based on the assumption of non-interference between the two binding sites from the NMR titration experiments, we also employed a two-site independent model in the ITC analysis, referred to as the ‘Two Set of Sites’ model in MicroCal ITC-ORIGIN Analysis Software (Malvern Panalytical). The two dissociation constants (K_D1_ and K_D2_ for NSH3 and CSH3, respectively) derived from the NMR and ITC data using the two-site independent binding model were summarised in Tables 1 and S1. Consistent with the trends of the above CSPs, the dissociation constants for NSH3 of S4, S5 and S9 were notably lower than those for CSH3. The dissociation constants of S10 showed no significant difference between NSH3 and CSH3.

**Figure 3.**
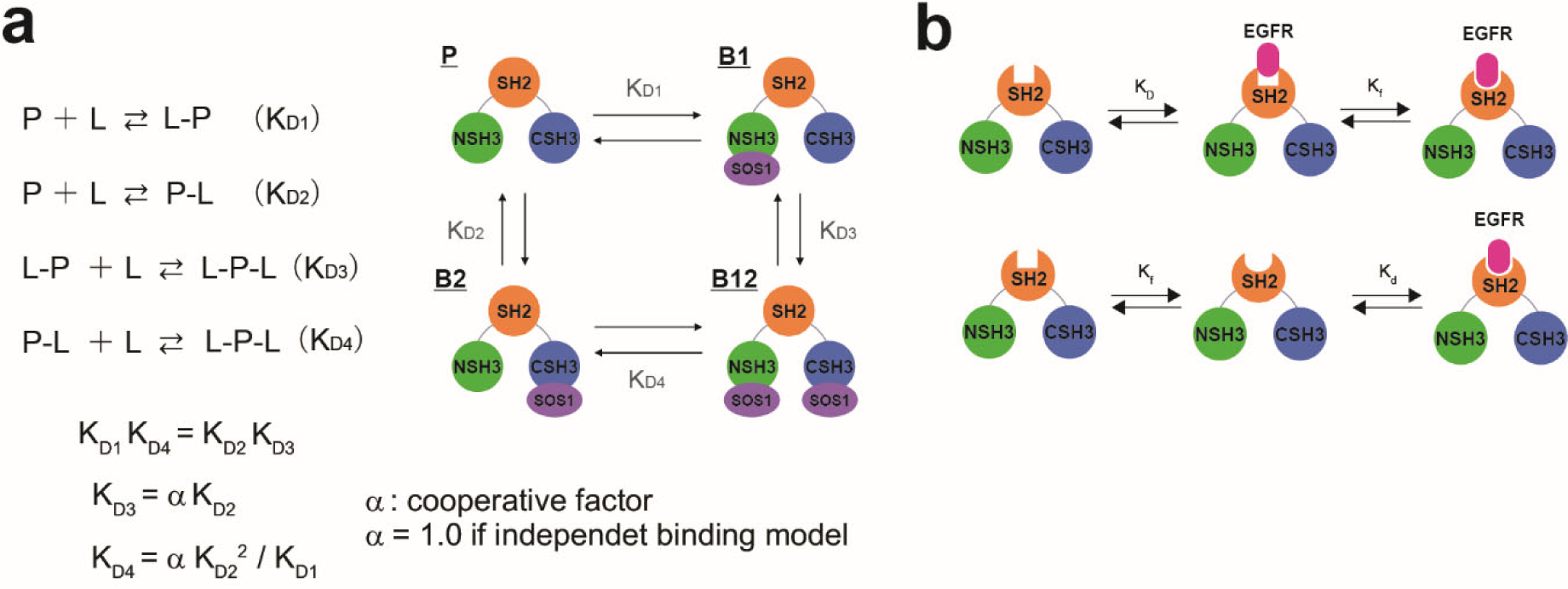
Ligand-binding equilibrium models assuming multiple coupling modes. (**a**) Schematic illustration for the ligand-binding equilibrium models assuming to include two-binding sites. The ligand-binding equilibrium models with two-binding sites are composed of four stages: ligand-free (P), ligand-bound on one site (L-P and P-L), and ligand-bound on two sites (L-P-L) (left). (**b**) Schematic illustration for the ligand-binding equilibrium models (left), assuming induced fit (upper) and conformational selection (bottom).

**Table 1.**
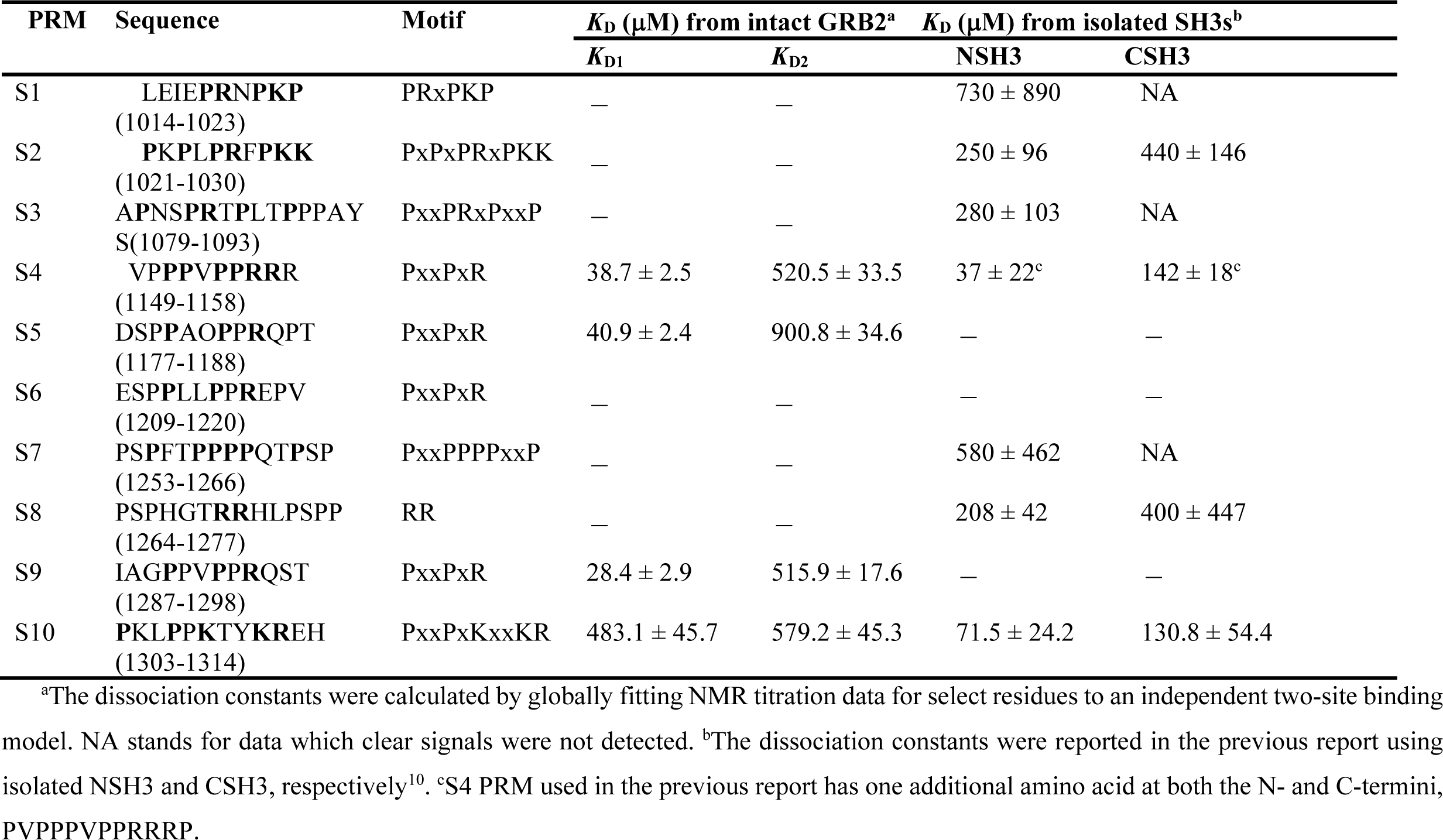
Dissociation constants for NSH3 and CSH3 against SOS1 PRMs in the full-length GRB2 and isolated SH3 domains.

The two dissociation constants for PRMs S4, S5, and S9 were quite similar among the three PRMs, with S9 exhibiting the lowest values. In contrast, those for S10 were approximately ten times higher compared to the others. The K_D1_ values associating the NSH3 domain for S4, S5, and S9 were significantly lower, by a factor of ten to twenty, than the corresponding K_D2_ values for CSH3. Notably, these variances are more pronounced than the 1.5 to 4.0-fold differences observed in the previous research using the isolated SH3 domains^10^. The profound differences in the K_D_ values between the two domains are also reflected in the CSP data (Fig. 2), where early titration points showed substantial CSPs for NSH3, whereas the perturbations for CSH3 were only observed at later stages of the titration. This considerable difference in binding affinity between the two domains could markedly impact the nature of the multivalent interactions with SOS1. Meanwhile, the two dissociation constants for S10 were nearly equivalent, or the K_D1_ for NSH3 was slightly lower than K_D2_ for CSH3. This also contrasts with the previous analysis employing the isolated SH3 domains in which its K_D1_ was four times larger than its K_D2_. This ten to twenty-fold difference between the two dissociation constants for S4, S5, and S9 was further corroborated by our ITC analysis of the full-length GRB2 (Table S1). The ITC analysis was also conducted for the remaining six PRMs. For S1, S3 and S7, the binding affinity to the two SH3 domains was likely so weak that little calorimetric change was observed, preventing the determination of their dissociation constants. For S2 and S8, their dissociation constants for one-side with higher binding affinity could be determined, but other constants with weaker affinity could not be obtained. Nevertheless, a comparison of the dissociation constants obtained from the ITC data suggests that these six PRMs have considerably lower affinities than S4, S5 and S9.

### NMR titration experiments with a phosphorylated peptide derived from EGFR with and without the SOS1 PRMs

It has been reported that GRB2, typically existing in a dimeric form in solution, dissociates into a monomeric form upon binding to EGFR^17^. Another study proposes an enhancement of the SH2 domain’s binding affinity to EGFR when SOS1 PRMs are bound to the SH3 domains^7^. To investigate the relationships between the binding affinities of the EGFR peptide and SOS1 PRMs, and the dimerisation of GRB2, we conducted titration experiments using a phosphorylated EGFR peptide (1062 — 1076;FLPVPEpYINQSVPKR) both with and without the binding of SOS1 S4 PRM (Fig. 4, S16, and S17). Major chemical shift perturbations, regardless of the presence of SOS1 S4 PRM, were predominantly localised in the region encompassing two β-sheets and α-helix (β6, β7, and α1; Ile65, Ala68, Arg86, Ser96, Phe108 and Lys109) within the SH2 domain, which is consistent with the previously identified interaction pocket with the EGFR peptide^18, 19^. Specifically, Phe108 and Lys109 exhibited substantial CSPs, likely due to proximity to the aromatic ring of the phospho-tyrosine at the centre of the EGFR peptide, causing a ring current effect. Ala68 presented a decrease in peak intensity and the emergence of a new peak during the titration experiment with SOS1 S4 PRM, suggesting a slow exchange between the free and the EGFR peptide-bound forms of GRB2 on the NMR timescale. This reduction in peak intensity for Ala68 was similarly observed in the spectra without SOS1 S4 PRM, but the corresponding new peak remained unassigned. Notably, significant CSPs were also identified in the regions far from the binding surface of the EGFR peptide (residues 72–74, 101– 104, 141,144) (Fig. 4). Considering that several EGFR peptides commonly form U-shaped structures upon binding to SH2 domains^19^, it is unlikely for the peptide to reach these distant areas. This observation may suggest an intrinsic conformational change within the SH2 domain itself. Meanwhile, a previous study indicated that the binding of another phosphorylated EGFR peptide (1062–1076; FLPVPEpYINQSVPKR) may promote the transition of GRB2 from its dimeric to monomeric form^17^. However, the CSPs in this region are also distant from the predicted dimeric interaction surface seen in the crystal structure, suggesting that the effect of the monomeric transition by binding with the phosphorylated EGFR peptide may not be obvious.

**Figure 4.**
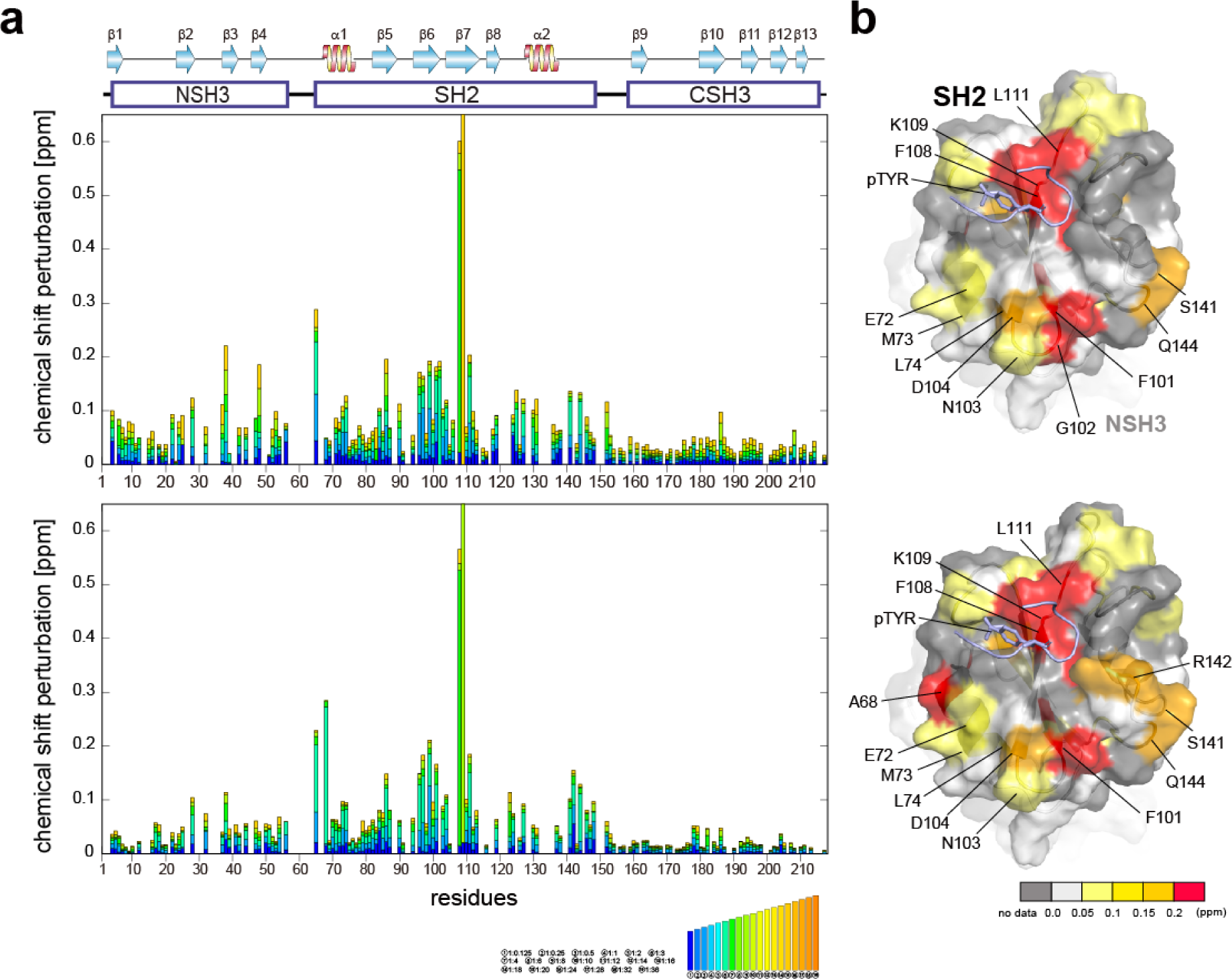
Plots of chemical shift perturbation of backbone ^1^H^N^ and ^15^N nuclei of GRB2 upon the titration with the EGFR phosphorylated peptide. (a) Bar plots of chemical shift perturbation of backbone ^1^H^N^ and ^15^N nuclei of GRB2 upon the titration of the EGFR phosphorylated peptide without (upper) and with (lower) SOS1 S4 PRM. The mean shift difference Δδ_ave_ was calculated as [(Δδ^1^H^N^)^2^ + (Δδ^15^N/5)^2^]^1/2^ where Δδ^1^H^N^ and Δδ^15^N are the chemical shift differences (ppm) between GRB2 on its own and in the presence of the PRMs. The bar graphs are colour-coded according to the protein-peptide concentration ratio and are overlaid. The proline residues as well as the residues for which ^1^H^N^-^15^N correlation cross peaks were not analysed due to signal overlap or other reasons are shown in grey. The secondary structures of GRB2 are also shown. (b) Chemical shift perturbation upon the titration with the EGFR phosphorylated peptide without (upper) and with (lower) SOS1 S4 PRM represented on the crystal structure of the GRB2 SH2 domain (PDB ID: 1BMB).

Interestingly, slight CSPs were observed not only in the SH2 domain but also in the two SH3 domains. A comparison of CSPs between NSH3 and CSH3 revealed a more pronounced overall chemical shift change in NSH3. The NSH3 domain is far from SH2 on the crystal structure (Fig. S1d), and therefore, the influence of NSH3 upon the EGFR peptide binding was unexpected. This observation further supports the model described earlier, suggesting that NSH3 and SH2 are in closer contact, while CSH3 exists independently.

### Dissociation constants of the EGFR peptides calculated from NMR chemical shift perturbation

Several studies have shown the flexibility of the GRB2 SH2 domain, which can be further subdivided into smaller sub-domains^20, 21^. They suggest that the binding of EGFR peptides might influence both the dynamics and conformation of these sub-domains. Our observations as described above, revealing pronounced chemical shift changes outside the recognised binding site, indicate that conformational changes may accompany with the SH2 domain binding to the phosphorylated EGFR peptide. To validate if our titration data imply the conformational change of the SH2 domain, we performed the calculation of the dissociation constants using either the conformational selection model or the induced fit model, as well as the simplest single-binding model (Fig. 3b and S10b). Detailed theory and models of these calculations are described in the materials and methods section. Model selection was based on the difference between experimental and predicted values and the 98% confidence intervals determined by bootstrap.

The results of the model validation via bootstrap for the three models demonstrated that there were no significant differences in residuals and standard deviations among the models, regardless of whether PRM S4 was bound or not (Fig. S10b). Consequently, the NMR titration data did not provide correlations between conformational changes and dissociation constants, and it was not determined whether the SH2 domain underwent conformational changes. Therefore, we adopted the dissociation constant from the most straightforward single-binding model. The derived dissociation constants were 4.2 ± 1.6 μM without S4 PRM and 2.3 ± 8.9 μM when S4 PRM was bound, approximately representing a two-fold increase in binding affinity for the S4 PRM-bound state (Table 2). This result is consistent with the Surface Plasmon Resonance (SPR) analysis, indicating a two-fold enhancement in the binding affinity of the phosphorylated EGFR peptide to GRB2 upon S4 PRM binding^7^. Our results validate the model that the SOS1 PRM binding promotes the association of phosphorylated EGFR, and it is very interesting that the interaction in the SH3 domains, which is quite distant in the crystal structure (Fig. S1d), affects the interaction with the EGFR peptide in the SH2 domain. Meanwhile, although the tendency of CSPs from the EGFR peptide titrations with and without SOS1 S4 PRM was clearly different (Fig. 4), the standard deviation values obtained by the bootstrap method were relatively large. Thus, further experiments will be required to confirm this.

**Table 2.** Dissociation constants in the full-length GRB2 against phosphorylated EGFR peptides.

| **Full-length GRB2** |  |  |
| --- | --- | --- |
| **protein** | **Sequence** | **$K_D$ (μM)** |
| GRB2 | EpYINVSQV | $4.21 \pm 1.61$ |
| GRB2+S4 | EpYINVSQV | $1.76 \pm 8.90$ |
| GRB2<sup>a</sup> | VPEpYINQSVPK | $0.713 \pm 0.145$ |
| GRB2+mSOS1<sup>a</sup> | VPEpYINQSVPK | $0.296 \pm 0.005$ |

<sup>a</sup> The dissociation constants obtained from ITC experiments in the previous works<sup>7</sup>.

The dissociation constants obtained in our study showed binding affinities that were 6–10 fold weaker than those obtained in the SPR analysis. This difference may be due to the phosphorylated EGFR peptide used in the present analysis being about four amino acids shorter than that used in the SPR analysis. Our previous analysis has also shown that differences in a few amino acids before and after the binding motif affect the magnitude of the overall dissociation constant^22^.

### Individual dissociation constants for each residue and docking simulations of PRMs on NSH3 and CSH3

NMR titration experiments of PRMs to the full-length GRB2 revealed that the NSH3 domain possesses binding affinities for PRMs that are ten to twenty times higher than those of CSH3. To clarify the underlying reasons for this difference, we calculated residue-specific dissociation constants fitted to chemical shift perturbations of individual nuclei instead of a global dissociation constant for a molecule and performed docking simulations to estimate the intermolecular interactions. Based on the distributions of the residues exhibiting significant chemical shift changes upon the SOS1 PRM interactions, docking simulations of the PRMs on NSH3 and CSH3 were carried out with AutoDock CrankPep (ADCP) software^23^. The S4, S5 and S9 PRMs, containing a PxxPxR motif, yielded similar dissociation constants from titration experiments, while the S10 PRM with a PxxPxKxxKR motif exhibited different dissociation constants. Consequently, docking simulations were selectively performed for S4 and S10. The top five models for S4 and S10 are depicted in Figure 5.

**Figure 5.**
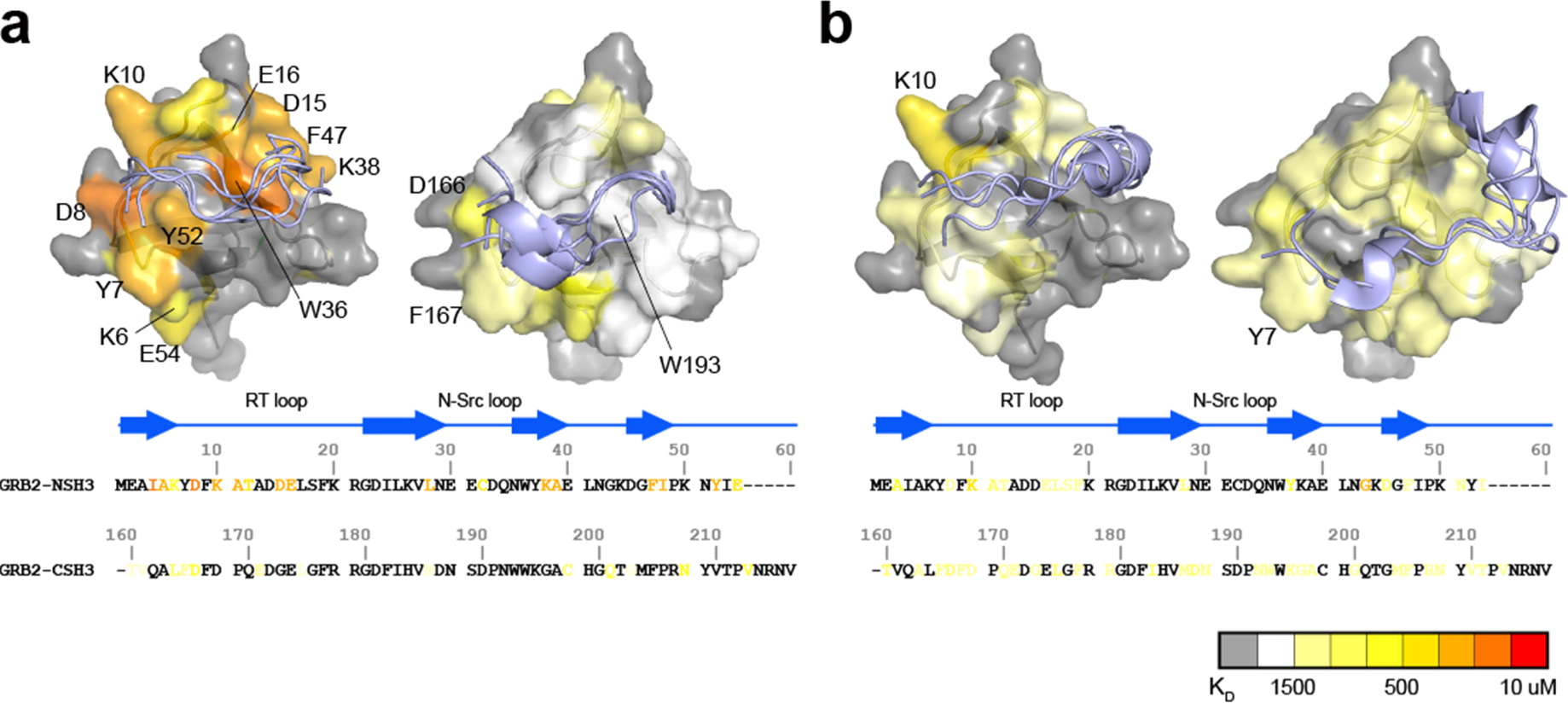
Comparisons of residue-specific K_D_s and docking simulations of S4 and S10 PRMs between NSH3 and CSH3. The top five structures of SOS1 PRM S4 (**a**) and S10 (**b**) with the lowest energy from the docking simulations are presented for NSH3 (left) and CSH3 (right), with mapping of residue-specific dissociation constants (K_D_s) on the crystal structure of the NSH3 (PDBID: 1AZE) and CSH3 domains (PDB ID: 1IO6). The magnitudes of residue-specific K_D_s are colour-coded both on the structures and aligned amino acid sequences of the SH3 domains.

As expected, the residue-specific dissociation constants with lower values and the PRM binding sites from the docking simulations were primarily localised to the RT loop and the adjacent β-strand. However, there was a clear difference in the pattern of residue-specific dissociation constants between NSH3 and CSH3. Regions around Y7 and D8 in NSH3, and F167 and D166 in CSH3, demonstrated high affinity for one edge of S4 PRM. Conversely, NSH3 exhibited high affinities at the opposite edge of S4 PRM, particularly at the region comprising D15, E16, W36, K38 and F47, whereas CSH3 showed markedly lower affinities at this site. This variance may account for the 10–20 fold difference in binding affinity for S4, S5, and S9 between NSH3 and CSH3. Regarding S10, the pattern of residue-specific dissociation constants was also considerably different between NSH3 and CSH3. In CSH3, the relatively high dissociation constants are broadly distributed across the surface of its domain without any significantly low values at specific sites. In addition, the top five conformations of S10 PRM predicted by docking simulations on CSH3 clustered into two groups: one associated with the RT loop and its neighbouring β-strand, and the other at the centre of the β-sheet. The dispersion for weak residue-specific dissociation constants over the domain’s surface and the non-arbitrary docking conformations suggest a unique binding mode for S10 PRM to CSH3.

## Discussion

GRB2 and SOS1 play a crucial role in the MAPK signalling pathway in transmitting signals from RTKs on the plasma membrane to RAS and subsequent downstream proteins. They are also thought to contribute to cell signalling regulation by acting as scaffold proteins and yielding condensed environments, such as droplets. The dynamic formation and dissolution of these droplets in living cells underscores the significance of analysing transient multivalent interactions between GRB2 and SOS1 at atomic resolution. In this study, we demonstrated the interactions and binding affinities between intact GRB2 and the SOS1 PRMs in detail mainly using solution NMR.

Comparison of the NMR spectra of the full-length GRB2 and its individual isolated domains revealed that the spectrum of the isolated CSH3 domain agrees well with that of the full-length GRB2, whereas the spectra of the isolated NSH3 and SH2 showed little similarity to the full-length protein. This inconsistency suggests that the NSH3 and SH2 domains of intact GRB2 engage in interdomain contacts in solution, while the CSH3 domain exists independently. In the crystal structure (Fig. S1d), the two SH3 domains are proximal to each other but distant from the SH2 domain, indicating that significant differences in domain orientations in the crystal compared to in solution, as predicted by previous studies^9^. Although the linker region between NSH3 and SH2 (linker 1) appears as a long random coil in the crystal structure, the amino acid sequence properties in the region, which includes several proline and aromatic residues (MKPHPWFF), suggest quite bulky and low flexibility in this region (Fig. S1d). This amino acid sequence property also supports our NMR data indicating that NSH3 and SH2 are in contact and CSH3 is independent.

NMR titration experiments with GRB2 and SOS1 PRMs showed significant chemical shift perturbations at the RT-loop as well as at the β-strand located parallel to the loop in both NSH3 and CSH3. These results confirm that this region is the primary binding site for PRMs in the two SH3 domains, as previously shown^10^. Meanwhile, chemical shift perturbations were also observed in the linker region and the SH2 domain, indicating that the PRM bindings may induce alterations in the relative domain orientations of the GRB2’s three domains and subsequently yield GRB2 dimerisation. Notably, the trajectories of the chemical shift changes in the region (Q157, T159, and Y160), near CSH3 in linker 2, were curved or U-turn shaped unlike the perturbation patterns in other regions, suggesting the presence of multiple-states with different chemical shifts in this area. This region corresponds to the crossing point of linkers 1 and 2 in the crystal structure (Fig. S1d), where an inhibitory effect on GRB2 dimerisation was observed upon mutation of Tyr160 to glutamate (Y160E)^6^. This implies that the PRM bindings may impact this crossed structure or the GRB2 dimerisation. Furthermore, the comparison of the NMR spectra between intact GRB2 and the three isolated domains, as well as the previous NMR relaxation experiment^9^, suggest that only the CSH3 domain of GRB2 exhibits independent motion (Fig. 6a). The structure predicted by AlphaFold 2 shows that while NSH3 and SH2 are adjacent to each other, CSH3 is distant from the others with low values of AlphaFold’s Model Confidence in the linker 2 connecting SH2 and CSH3. This also supports the model that the CSH3 domain moves independently. These significant dynamics of CSH3 may facilitate dynamic equilibrium between dimeric and monomeric conformations (Fig. 6a) and yield dynamic droplet formation and dissociation by GRB2, SOS1, and LAT.

**Figure 6.**
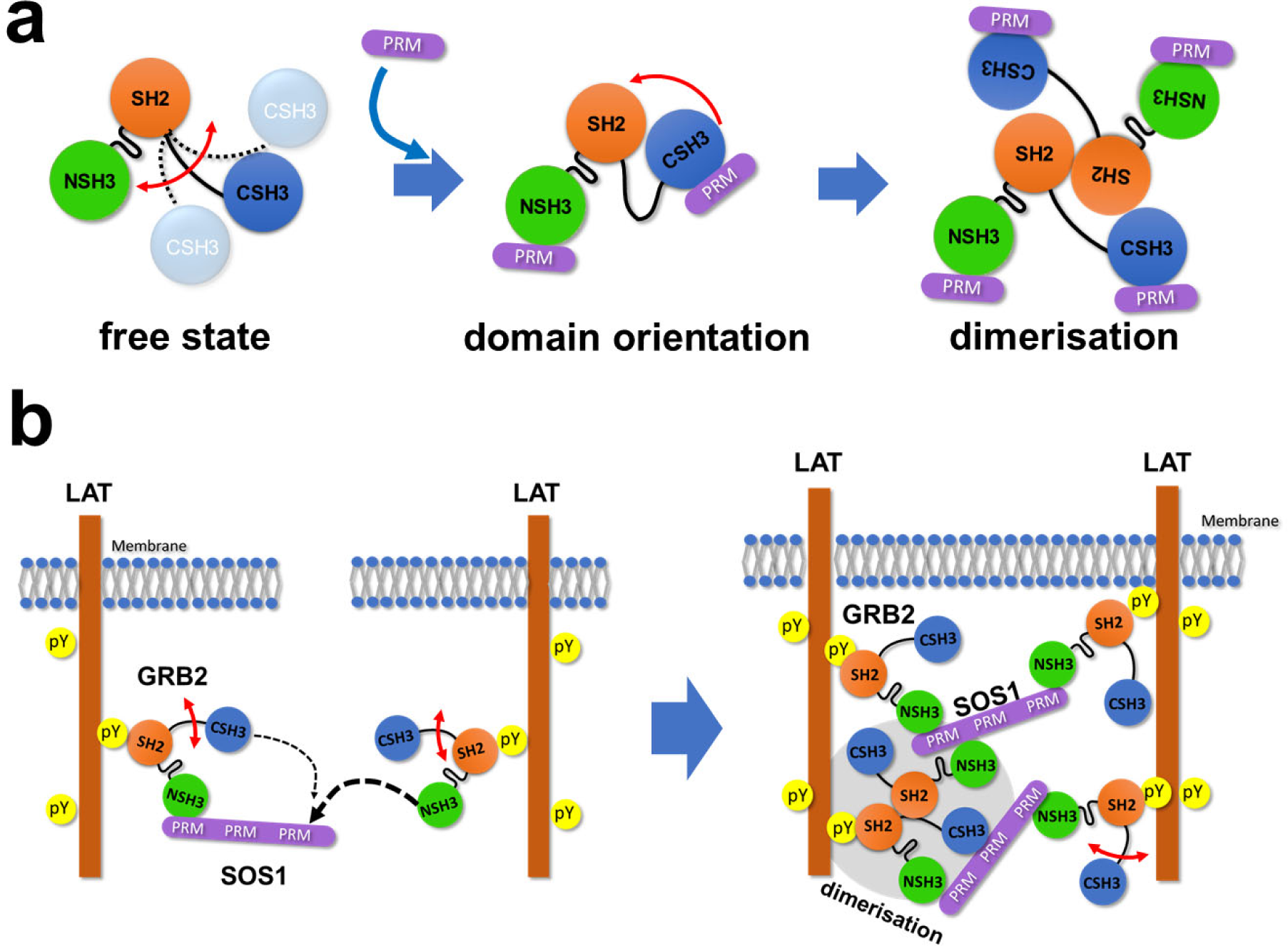
Schematic illustrations of the interactions of GRB2 with the SOS1 PR domain and LAT. (a) A schematic illustration of GRB2 from free to complex. (b) A schematic illustration of liquid-liquid phase separation by GRB2, SOS1 and LAT. The thickness of dotted arrows indicates the strength of binding affinity.

The calculated dissociation constants revealed that the binding affinities of NSH3 for the PRMs S4, S5 and S9 were 10–20 times higher than those of CSH3. The differences in affinity between NSH3 and CSH3 in the analysis of the full-length GRB2 were considerably larger than the 4-to 5-fold difference obtained by the previous research isolating the individual domains^10^. It is conceivable that two binding sites with different binding affinities, high and low ones, coexist and compete in the interaction with a counterpart binning site, making the difference more pronounced. This significant difference between the two experiments indicates that analysing using the full-length GRB2 is necessary for accurate measurement of the protein’s affinities.

GRB2 possesses two SH3 domains which are thought to have almost identical functions but very different binding affinities. This biological significance must be considered. According to the database OpenCell (https://opencell.czbiohub.org)^24^, the GRB2 concentration in HEK293 cells is 1.4μM, whereas the SOS1 concentration is 32 nM. This shows that SOS1 is considerably less abundant than GRB2 in intracellular environments, and the two SH3 domains of a GRB2 molecule are likely to competitively bind to PRM sites in a single SOS1 molecule. This interaction process may be mediated by the following mechanism: First, NSH3 of GRB2 binds to one of the PRMs of SOS1 due to its higher affinity. Second, whereas CSH3 of the same GRB2 molecule has more chances to bind to the remaining sites of the PRMs due to their spatial proximity and the single-strand connection, NSH3 of a second GRB2 molecule, containing higher affinity, preferentially interacts with the PRMs than CSH3 of the first GRB2 (Fig. 6b, left). This leads to the bindings of several GRB2 molecules to a single SOS1 and the formation of GRB2-SOS1 cross-links. This process might be a key for the LLPS formation or network structures by GRB2 and SOS1 (Fig. 6b, right). Furthermore, CSH3 is expected to have weak contacts with other domains, to move independently, and to be highly mobile, which could help capture remaining free PRMs. These asymmetric physical properties of the two SH3 domains of GRB2, including dynamics and binding affinity, suggest that GRB2 has a mechanism beyond the simple function of binding to SOS1 and recruiting it near the transmembrane, as previously thought. These properties may enable GRB2 to form complex multivalent interactions and bridges among molecules, potentially facilitating LLPS formation. In terms of the formation of complex networks, it is also interesting to note that exclusively for the SOS1 S10 PRM, NSH3 and CSH3 exhibit comparable binding affinities and no U-turn shape trajectories of CSPs of residues 157, 159, and 160, indicating that GRB2 can engage in multivalent interactions with different binding modes and different affinities to SOS1. As described in the previous research, dimerisation of GRB2 would also contribute to the complex network in LLPS (Figure 6b, left).

In this study, we analysed the interaction of full-length GRB2 with proline-rich motifs (PRMs) in detail, primarily using solution NMR. Solution NMR provides information on binding affinity and dynamics at atomic resolution. We analysed the data in statistical detail, considering several possible models, to extract the maximum amount of information from the NMR data. We believe that the extensive modelling presented here will become increasingly effective in biophysical fields. The models and calculation methods are packaged as a single software, which can be made available upon request. By using a large amount of data and these data analysis methods, we have succeeded in observing local interaction patterns and changes in the relative orientations of the three domains of GRB2 that cannot be seen by ITC and SPR, which have been commonly used for interaction analysis. NMR is particularly useful in the structural analysis of highly flexible proteins such as multidomain proteins, IDPs, and RNAs. Thus, we believe that it will further contribute to the understanding of biological phenomena such as liquid-liquid phase separation, in which these molecules are often involved, at atomic resolution.

## Material and Methods

### Protein expression

The gene encoding human GRB2 in addition to a N-terminal His_6_ tag with a tobacco etch virus (TEV) protease cleavage site was constructed into the vector pET14b (Merck) for over-expression in the *Escherichia coli* BL21 (DE3) pLysS strain. Uniformly ^13^C/^15^N-labelled GRB2 protein was obtained by growing the bacteria at 37 °C in M9 minimal media, containing [^13^C_6_]-glucose (ISOTEC) and ^15^NH_4_Cl (ISOTEC) as the sole carbon and nitrogen sources, respectively, supplemented with 20 mM MgSO_4_, 0.1 mM CaCl_2_, 0.4 mg/ml thiamin, 20 μM FeCl_3_, salt mix (4 μM ZnSO_4_, 0.7 μM CuSO_4_, 1 μM MnSO_4_, 4.7 μM H_3_BO_3_), and 50 mg/l ampicillin. Protein expression was induced by adding 119 mg/l isopropyl β-D-thiogalactoside (IPTG) at an OD 600 nm of 0.5. After 12 h of further growth at 20 °C, cells were harvested. Uniformly ^15^N-labelled GRB2 was produced identically except using [^12^C_6_]-glucose. Uniformly 90 % ^2^H/^13^C/^15^N-labelled GRB2 was obtained by growing cells in M9 medium containing 90 % ^2^H_2_O/10 % ^1^H_2_O. ^15^N-Leu and Lys selectively labelled GRB2 samples were obtained by growing cells in M9 medium supplemented with ^15^N-Lys and ^15^N-Leu, respectively, when protein expression was induced.

GRB2 SH2 domain (54–159) in addition to a N-terminal GST tag, a TEV protease cleavage site, and GG linker was constructed into the expression vector pLEICS02 and over-expressed in the *E. coli* Rosetta^TM^ 2 (DE3) strain (Merk). Uniformly ^13^C/^15^N and ^15^N-labelled GRB2 SH2 samples were prepared by growing the bacteria at 37 °C in LB media in the same M9 minimal media described above, containing [^13^C_6_]-glucose and ^15^NH_4_Cl as the sole carbon and nitrogen sources, respectively. The expression was induced by adding IPTG at an OD 600 nm of 0.5. The cells were further incubated for 18 h at 20 °C, and harvested.

S1(LEIEPRNPKP), S2(PKPLPRFPKK), S3(APNSPRTPLTPPPAYS), S4(VPPPVPPRRR), S5(DSPPAIPPRQPT), S6(ESPPLLPPREPV), S7(PSPFTPPPPQTPSP), S8(PSPHGTRRHLPSPP), S9(IAGPPVPPRQST), and S10(PKLPPKTYKREH) were commercially obtained from Eurofins Genomics. EGFR (EpYINSQV) peptides were obtained from Toray Research Center Incorporated.

### Protein purification

All the procedures described below were carried out at 4 °C unless stated otherwise. All isotope-labelled GRB2 samples were purified in the same way. The cells dispersed in buffer A [50 mM Tris-HCl (pH 7.5), 20 mM Imidazole] were disrupted by sonication for 30 min on ice in the presence of hen egg lysozyme (0.1 mg/ml). The cell debris was clarified by centrifugation at 14,000 *g* for 1 h. The supernatant was loaded onto a Ni-NTA agarose bead slurry (Qiagen) histidine-tagged affinity column pre-equilibrated with buffer A. After washing the column with buffer A until sufficiently low UV absorption at 280 nm, the GRB2 protein was eluted by linearly increasing the concentration of Imidazole from 20 to 500 mM with a flow rate of 1 ml/min in buffer A using Econo Gradient Pump (BIO-RAD). The fractions containing the target protein were concentrated to 5 ml with Amicon Ultra-15 10 kDa (Merck). The concentrated sample was incubated with uTEV3 protease^25^(Addgene) to remove the His_6_ tag for approximately 20 h at 25 °C. The sample was loaded onto a HiLoad 16/600 Superdex 75pg column (Cytiva) gel filtration column with a flow rate of 1 ml/min in buffer B [50 mM Tris-HCl (pH 7.5), 1 mM DTT, 1 mM EDTA, 150 mM NaCl] using a FPLC system (AKTA pure 25, Cytiva). The 5 ml sample concentrated, from the fractions involving the target proteins, with Amicon Ultra-15 10 kDa was loaded on a Resource Q (Cytiva) anion-exchange column pre-equilibrated with buffer C [50 mM Tris-HCl (pH 7.5), 1 mM DTT, 1 mM EDTA] using the FPLC system. After washing the column with 30 ml of buffer C, the GRB2 protein was eluted by linearly increasing the concentration of NaCl from 5 mM to 300 mM with a flow rate of 0.5 ml/min in buffer C. The purity of the GRB2 samples was confirmed in each step by SDS-PAGE. Protein concentrations were determined by NanoDrop 2000 (ThermoFisher) measuring UV absorption at 280 nm. Protein samples for NMR measurements were concentrated and dissolved in NMR buffer [90% ^1^H_2_O/ 10% ^2^H_2_O containing 20 mM Na_2_HPO_4_-NaH_2_PO_4_, pH 7.2, 0.05% NaN_3_].

For purification of GRB2 SH2 domain, the harvested cells were re-suspended in a phosphate buffer [pH 7.2, 1 % Triton X100, 1 % 2-Mercaptoethanol, 0.1 mM p-Amidinophenyl Methansulfonyl Fluoride (p-APMSF; FUJIFILM), 0.1% Halt^TM^ Protenase Inhibitor single-use cocktail EDTA-free (Thermo Scientific)] and lysed by sonication in the presence of hen egg lysozyme (0.1 mg/ml). The cleared lysate was collected after centrifugation. The supernatant was loaded onto Glutathione Sepharose 4B (Cytiva) pre-equilibrated with buffer D [50 mM sodium phosphate (pH 7.2), 150 mM NaCl]. The column was washed with 6 column volumes of the same buffer, followed by elution with a linear gradient of 0-50 mM glutathione. The fractions containing GRB2 SH2 domain were concentrated by Amicon Ultra-15 3 kDa (Merck) and then incubated with uTEV3 protease to remove the GST tag in buffer D for approximately 16 h at 25 °C. The cleaved sample was loaded onto a HiLoad 16/60 Superdex 75 column pre-equilibrated with D buffer. The final GRB2 SH2 domain fractions were dissolved in NMR buffer [90% ^1^H_2_O/ 10% ^2^H_2_O containing 20 mM Na_2_HPO_4_-NaH_2_PO_4_, pH 7.2, 0.05% NaN_3_].

### NMR spectroscopy

NMR experiments for GRB2 were performed at 25°C probe temperature on Bruker AVANCE-III HD 600 MHz and AVANCE-III 700 MHz spectrometers equipped with pulsed field gradient triple-resonance cryoprobes. NMR experiments for GRB2 SH2 domain were measured at 25°C probe temperature on Bruker AVANCE-III 600 MHz equipped with pulsed field gradient triple-resonance room-temperature probes. All spectra were processed with the Azara software package. For all 3D NMR data, a non-uniform sampling (NUS) scheme was used for the indirectly observed dimensions to reduce experimental time. In brief, approximately 1/8 of the points were selected in the Poisson-gap sampling from the conventional regularly spaced grid of *t_1_*, *t_2_* points^26^. The two-dimensional maximum entropy method (2D MEM) or Quantitative Maximum Entropy (QME) ^27^ was applied for the indirectly acquired two dimensions after processing the directly acquired dimension (*t_3_*) by Fourier transformation using the Azara 2.8 software. NMR spectra were visualised and analysed using the CcpNmr Analysis 2.5.0 software^28^. Backbone resonance assignments for GRB2 were achieved by analysing 3D triple-resonance NMR spectra, TROSY-HNCO, TROSY-HN(CA)CO, TROSY-HNCA, TROSY-HN(CO)CA, TROSY-HN(CB)CA, TROSY-HN(COCB)CA, measured on 90%-^2^H/^13^C/^15^N-labelled samples. Sidechain resonance assignments for GRB2 were performed by analysing 3D HBHA(CBCACO)NH, (H)CC(CO)NH, H(CCCO)NH, HCCH-COSY, and HCCH-TOCSY spectra, measured on ^13^C/^15^N-labelled samples. The ^1^H/^15^N resonance assignments of isoleucine, leucine and lysine were confirmed unambiguously by 2D ^1^H-^15^N HSQC spectra measured on the Ile-, Leu- and Lys-selectively ^15^N-labelled samples. Backbone and sidechain resonance assignments for GRB2 SH2 domain were achieved by analysing 3D triple-resonance NMR spectra, HNCO, HN(CA)CO, HNCA, HN(CO)CA, CBCANH, CBCA(CO)NH, HBHA(CBCACO)NH, H(CCCO)NH, CC(CO)NH, HCCH-COSY, and HCCH-TOCSY, measured on ^13^C/^15^N-labelled samples. The sidechain aromatic ^1^H/^13^C resonances of phenylalanine, tyrosine and tryptophan residues were confirmed by analysing 2D (HB)CB(CGCD)HD and (HB)CB(CGCDCE)HE spectra. For the collection of NOE-derived distance restraints for GRB2 and SH2 domain, 3D ^15^N-separated and 3D ^13^C-separated NOESY-HSQC spectra were measured on uniformly ^15^N-labelled or ^13^C/^15^N-labelled samples, respectively. A 100 ms NOE mixing period was employed for the 3D NOESY experiments.

The titration experiments with the SOS1- and EGFR-derived peptides were performed in an NMR buffer [20 mM sodium phosphate, 0.05% (w/v) NaN_3_] at 25 °C. ^15^N-labelled GRB2 was successively titrated up to a molar ratio of 1:12 with the S4, 1:36 with S5 and S9, 1:24 with S10, and 1:2.0 with EGFR peptide, respectively. 2D ^1^H-^15^N HSQC spectra were measured at each titration point. Chemical shift differences, *Δ*, were analysed by *Δ* = ((*Δ*_H_)^2^ + (*Δ*_N_/5)^2^)^1/2^, where *Δ*_H_ and *Δ*_N_ are the differences in ppm for the backbone amide ^1^H and ^15^N chemical shifts between the two conditions. 1 ppm corresponds to 600.13 Hz for ^1^H and 60.81 Hz for ^15^N.

### Isothermal titration calorimetry (ITC)

GRB2 and SOS1 S1-S10 PRMs were separately dissolved in a buffer [20 mM sodium phosphate (pH 7.2)]. The ITC experiments were carried out at 25°C on a MicroCal Auto-iTC 200 instrument (Malvern Panalytical) with the following experimental parameters: 20 injections per experiment; 2 μL injection volume, except that the first injection of 0.4 μL to be discarded at the analysis; 150 s waiting period between the injections; stirring at 750 rpm; 5 μcal/sec reference power. The binding experiments were performed by injecting 1.0 mM PRM solution into a mixture of 50 μM GRB2. Each binding experiment was accompanied with a corresponding reference experiment exclusively with the sodium phosphate buffer and ultrapure water. Each parameter was determined by analyzing row ITC data with the model, Two Set of Sites, using MicroCal ITC-ORIGIN Analysis Software (Malvern Panalytical).

### Lineshape analysis of two-dimensional signals and determination of *K*_D_s and *K*_f_

The dissociation constants for each residue were calculated by non-linear regression analysis with the equation for the single state model^29^:

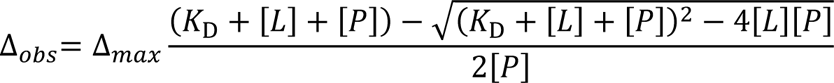

where Δ_obs_ is the observed chemical shift perturbation, Δ_max_is the maximum chemical shift perturbation, *K*_D_ is the dissociation constant, and [L] and [P] are the ligand and protein concentrations respectively.

For the conformational selection model, the *K*_D_ and *K*_f_ between two states were calculated with the equation.

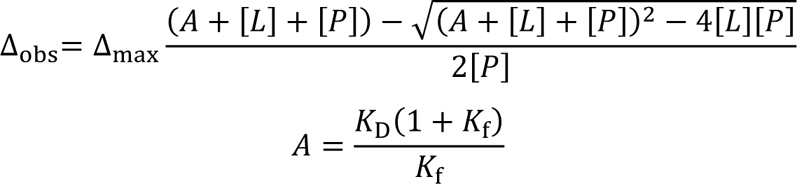

For the induced-fit model, the *K*_D_ and *K*_f_ between free and complex states were computed with the equation.

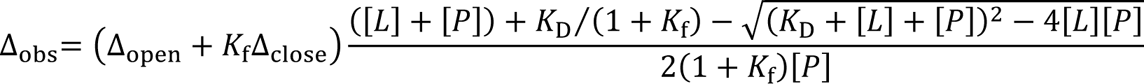

where Δ_open_ and Δ_close_ are the chemical shift perturbations in the active and inactive states for the complex formation.

For the two-site independent binding site model^15^, the *K*_D1_ and *K*_D2_ between P-L and L-P complex states in Figure 3a were computed with the equation.

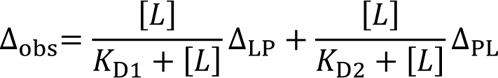

where Δ_LP_ and Δ_PL_ are the chemical shift perturbations in the P-L and L-P complex states.

For the two-site cooperative binding site model,

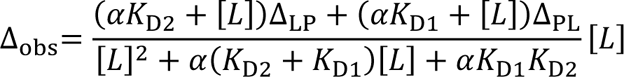

where α is the modulation factor.

In all models, the non-linear least-squares fitting was performed by the Levenberg–Marquardt algorithm. The precision of the model parameters was assessed by conducting the bootstrap resampling on the residuals of the regression model. This procedure was repeated 1000 times to generate the bootstrap distribution of the model parameters. The bootstrap confidence intervals of the model parameters were computed with the percentile method, i.e., the 25th and the 975th smallest values of the bootstrap distribution were regarded as the lower and the upper limits of the 95% bootstrap confidence intervals, respectively.

### Molecular docking

Ab initio modelling of NSH3 and CSH3 of GRB2 with SOS1-derived peptides was performed to explore the binding mode of the peptides onto the protein surface of GRB2-NSH3 and CSH3 using the program AutoDock CrankPep (ADCP) 0.1^23^. PDB IDs 6SDF and 1IO6 were used for template structures of the NSH3 and CSH3 domains, respectively. A grid box was defined using the program PMV 1.5.7^30^ so as to cover the region where the chemical shift perturbation was observed. For NSH3 in the cases of S4, S5 and S9 bindings, its grid dimensions were set to 23.250 x 26.250 x 17.750 Å^3^ with a spacing of 0.375 Å and a centre coordinate of 15.958, −19.046 and 21.781 in x, y, and z, respectively, while 25.500 x 24.750 x 20.250 Å^3^ with a same spacing and a centre coordinate of 20.176, −18.234 and 20.379 in x, y, and z for S10. Similarly, for CSH3 in the cases of S4 and S9 bindings, its grid dimensions were set to 24.000 x 25.500 x 18.000 Å^3^ with a spacing of 0.375 Å and a centre coordinate of 13.747, 2.841 and −4.020 in x, y, and z, respectively, while 21.000 x 25.500 x 18.000 Å^3^ with a same spacing and a centre coordinate of 16.595, 2.841 and −4.020 in x, y, and z for S5, and 12.358 x 24.000 x 18.000 Å^3^ with a same spacing and a centre coordinate of 12.358, 2.282 and −1.809 in x, y, and z for S10. A total of 4000 confirmers were calculated with 10000000 steps. Default settings were used for all other parameters.

## Data availability

All NMR chemical shifts have been deposited in the BioMagResBank with accession numbers 52412. All other data are available from the corresponding author upon reasonable request.

## Acknowledgment

The authors thank Dr. Yuji Sguita (RIKEN) and Prof. Noritaka Nishida (Chiba University) for useful discussion on general features of GRB2 and SOS1 interactions. We gratefully acknowledge financial support from the Funding Program for Core Research for Evolutional Science and Technology (CREST; JPMJCR13M3 to Y.I., JPMJCR21E5 to T.I.) from the Japan Science and Technology Agency (JST), Grants-in-Aid for Scientific Research (JP15K06979 to T.I., JP19H05645 to Y.I.) and Scientific Research on Innovative Areas (JP15H01645, JP16H00847, JP17H05887 and JP19H05773 to Y.I. JP26102538, JP25120003, JP16H00779 and JP21K06114 to T.I.) from the Japan Society for the Promotion of Science (JSPS), Shimadzu foundation, and the Precise Measurement Technology Promotion Foundation. The NMR experiments were performed at NMR Platform supported by the Ministry of Education, Culture, Sports, Science and Technology (MEXT) Program Grant Number JPMXS0450100021. The NMR experiments were performed at NMR Platform supported by the Ministry of Education, Culture, Sports, Science and Technology (MEXT) Program Grant Number JPMXS0450100021.

## Author Contributions

T.I, and Y.I designed the research and wrote the manuscript. K.T., T.A., M.T., H.S., T.H., K.I., T.M., and T.I. conducted sample preparation. T.I. implemented the model fitting software from NMR data. K.T., T.A., S.PM., H.S., Y.I., and T.I. performed NMR measurement, data analysis and docking simulation.

## References

1. A. D. Cox and C. J. Der, Ras history: The saga continues, Small GTPases, 2010, 1, 2–27.

2. Y. Pylayeva-Gupta, E. Grabocka and D. Bar-Sagi, RAS oncogenes: weaving a tumorigenic web, Nat Rev Cancer, 2011, 11, 761–774.

3. X. Su, J. A. Ditlev, E. Hui, W. Xing, S. Banjade, J. Okrut, D. S. King, J. Taunton, M. K. Rosen and R. D. Vale, Phase separation of signaling molecules promotes T cell receptor signal transduction, Science, 2016, 352, 595–599.

4. D. Milovanovic, Y. Wu, X. Bian and P. De Camilli, A liquid phase of synapsin and lipid vesicles, Science, 2018, 361, 604–607.

5. J. K. Chung, W. Y. C. Huang, C. B. Carbone, L. M. Nocka, A. N. Parikh, R. D. Vale and J. T. Groves, Coupled membrane lipid miscibility and phosphotyrosine-driven protein condensation phase transitions, Biophys J, 2021, 120, 1257–1265.

6. C. W. Lin, L. M. Nocka, B. L. Stinger, J. B. DeGrandchamp, L. J. N. Lew, S. Alvarez, H. T. Phan, Y. Kondo, J. Kuriyan and J. T. Groves, A two-component protein condensate of the EGFR cytoplasmic tail and Grb2 regulates Ras activation by SOS at the membrane, Proc Natl Acad Sci U S A, 2022, 119, e2122531119.

7. Y. M. Chook, G. D. Gish, C. M. Kay, E. F. Pai and T. Pawson, The Grb2-mSos1 complex binds phosphopeptides with higher affinity than Grb2, J Biol Chem, 1996, 271, 30472–30478.

8. S. Maignan, J. P. Guilloteau, N. Fromage, B. Arnoux, J. Becquart and A. Ducruix, Crystal structure of the mammalian Grb2 adaptor, Science, 1995, 268, 291–293.

9. S. Yuzawa, M. Yokochi, H. Hatanaka, K. Ogura, M. Kataoka, K. Miura, V. Mandiyan, J. Schlessinger and F. Inagaki, Solution structure of Grb2 reveals extensive flexibility necessary for target recognition, J Mol Biol, 2001, 306, 527–537.

10. T. J. Liao, H. Jang, R. Nussinov and D. Fushman, High-Affinity Interactions of the nSH3/cSH3 Domains of Grb2 with the C-Terminal Proline-Rich Domain of SOS1, Journal of the American Chemical Society, 2020, 142, 3401–3411.

11. S. H. Stitzinger, S. Sohrabi-Jahromi and J. Soding, Cooperativity boosts affinity and specificity of proteins with multiple RNA-binding domains, NAR Genom Bioinform, 2023, 5, lqad057.

12. L. Pinet, Y. H. Wang, A. Vogel, F. Guerlesquin, N. Assrir and C. V. Heijenoort, [Formula: see text]H, [Formula: see text]C and [Formula: see text]N assignments of human Grb2 free of ligands, Biomol NMR Assign, 2020, 14, 323–327.

13. U. Schaeper, N. H. Gehring, K. P. Fuchs, M. Sachs, B. Kempkes and W. Birchmeier, Coupling of Gab1 to c-Met, Grb2, and Shp2 mediates biological responses, J Cell Biol, 2000, 149, 1419–1432.

14. M. Harkiolaki, T. Tsirka, M. Lewitzky, P. C. Simister, D. Joshi, L. E. Bird, E. Y. Jones, N. O’Reilly and S. M. Feller, Distinct binding modes of two epitopes in Gab2 that interact with the SH3C domain of Grb2, Structure, 2009, 17, 809–822.

15. M. Arai, J. C. Ferreon and P. E. Wright, Quantitative analysis of multisite protein-ligand interactions by NMR: binding of intrinsically disordered p53 transactivation subdomains with the TAZ2 domain of CBP, Journal of the American Chemical Society, 2012, 134, 3792–3803.

16. H. Yagi, T. Kasai, E. Rioual, T. Ikeya and T. Kigawa, Molecular mechanism of glycolytic flux control intrinsic to human phosphoglycerate kinase, Proc Natl Acad Sci U S A, 2021, 118.

17. Z. Ahmed, Z. Timsah, K. M. Suen, N. P. Cook, G. R. t. Lee, C. C. Lin, M. Gagea, A. A. Marti and J. E. Ladbury, Grb2 monomer-dimer equilibrium determines normal versus oncogenic function, Nat Commun, 2015, 6, 7354.

18. J. E. DeLorbe, J. H. Clements, B. B. Whiddon and S. F. Martin, Thermodynamic and Structural Effects of Macrocyclic Constraints in Protein−Ligand Interactions, ACS Medicinal Chemistry Letters, 2010, 1, 448–452.

19. P. Nioche, W. Q. Liu, I. Broutin, F. Charbonnier, M. T. Latreille, M. Vidal, B. Roques, C. Garbay and A. Ducruix, Crystal structures of the SH2 domain of Grb2: highlight on the binding of a new high-affinity inhibitor, J Mol Biol, 2002, 315, 1167–1177.

20. K. Sanches, I. P. Caruso, F. C. L. Almeida and F. A. Melo, The dynamics of free and phosphopeptide-bound Grb2-SH2 reveals two dynamically independent subdomains and an encounter complex with fuzzy interactions, Sci Rep, 2020, 10, 13040.

21. A. Sandouk, Z. Xu, S. Baruah, M. Tremblay, J. B. Hopkins, S. Chakravarthy, L. Gakhar, N. J. Schnicker and J. C. D. Houtman, GRB2 dimerization mediated by SH2 domain-swapping is critical for T cell signaling and cytokine production, Sci Rep, 2023, 13, 3505.

22. P. M. Sayeesh, M. Iguchi, Y. Suemoto, J. Inoue, K. Inomata, T. Ikeya and Y. Ito, Interactions of the N-and C-Terminal SH3 Domains of Drosophila Drk with the Proline-Rich Peptides from Sos and Dos, Int J Mol Sci, 2023, 24.

23. Y. Zhang and M. F. Sanner, AutoDock CrankPep: combining folding and docking to predict protein-peptide complexes, Bioinformatics, 2019, 35, 5121–5127.

24. N. H. Cho, K. C. Cheveralls, A. D. Brunner, K. Kim, A. C. Michaelis, P. Raghavan, H. Kobayashi, L. Savy, J. Y. Li, H. Canaj, J. Y. S. Kim, E. M. Stewart, C. Gnann, F. McCarthy, J. P. Cabrera, R. M. Brunetti, B. B. Chhun, G. Dingle, M. Y. Hein, B. Huang, S. B. Mehta, J. S. Weissman, R. Gomez-Sjoberg, D. N. Itzhak, L. A. Royer, M. Mann and M. D. Leonetti, OpenCell: Endogenous tagging for the cartography of human cellular organization, Science, 2022, 375, eabi6983.

25. M. I. Sanchez and A. Y. Ting, Directed evolution improves the catalytic efficiency of TEV protease, Nat Methods, 2020, 17, 167–174.

26. S. G. Hyberts, K. Takeuchi and G. Wagner, Poisson-gap sampling and forward maximum entropy reconstruction for enhancing the resolution and sensitivity of protein NMR data, Journal of the American Chemical Society, 2010, 132, 2145–2147.

27. J. Hamatsu, D. O’Donovan, T. Tanaka, T. Shirai, Y. Hourai, T. Mikawa, T. Ikeya, M. Mishima, W. Boucher, B. O. Smith, E. D. Laue, M. Shirakawa and Y. Ito, High-resolution heteronuclear multidimensional NMR of proteins in living insect cells using a baculovirus protein expression system, J. Am. Chem. Soc., 2013, 135, 1688–1691.

28. W. F. Vranken, W. Boucher, T. J. Stevens, R. H. Fogh, A. Pajon, M. Llinas, E. L. Ulrich, J. L. Markley, J. Ionides and E. D. Laue, The CCPN data model for NMR spectroscopy: development of a software pipeline, Proteins, 2005, 59, 687–696.

29. M. P. Williamson, Using chemical shift perturbation to characterise ligand binding, Prog Nucl Magn Reson Spectrosc, 2013, 73, 1–16.

30. M. F. Sanner, A component-based software environment for visualizing large macromolecular assemblies, Structure, 2005, 13, 447–462.

