## Supplementary Figure S1-17 and Table S1 for "Different molecular recognition by three domains of the full-length GRB2 to SOS1 proline-rich motifs and EGFR phosphorylated sites"

[a]Department of Chemistry, Graduate School of Science, Tokyo Metropolitan University, 1-1 Minamiosawa, Hachioji, Tokyo 192-0397, Japan; [b]RIKEN Center for Biosystems Dynamics Research, RIKEN, 1-7-22 Suehiro-cho, Tsurumi-ku, Yokohama 230-0045, Japan; [c]Theoretical Molecular Science Laboratory, RIKEN Cluster for Pioneering Research, Wako, Saitama 351-0198, Japan; [d]Laboratory for Biomolecular Function Simulation, RIKEN Center for Biosystems Dynamics Research, Kobe, Hyogo 650-0047, Japan; [e]Computational Biophysics Research Team, RIKEN Center for Computational Science, Kobe, Hyogo 650-0047, Japan

[†] These authors contributed equally to this work

### Contents

1. **Supplementary Figure S1.** Domain and three dimensional (3D) structures of GRB2 and SOS1, and schematic illustration of LLPS.
2. **Supplementary Figure S2.** Overlays of 2D $^{1}H$-$^{15}N$ HSQC spectra of $^{15}N$-uniformly and amino-acid-selectively labelled samples.
3. **Supplementary Figure S3.** Backbone resonance assignments of GRB2.
4. **Supplementary Figure S4.** Table summarising the backbone and side-chain resonance assignments of GRB2.
5. **Supplementary Figure S5.** Overlays of 2D $^{1}H$-$^{15}N$ HSQC spectra of the full-length GRB2 and each isolated domain.
6. **Supplementary Figure S6.** Overlays of 2D $^{1}H$-$^{15}N$ HSQC spectra from multipoint titrations of $^{15}N$-labelled GRB2 with SOS-S5 PRM.
7. **Supplementary Figure S7.** Overlays of 2D $^{1}H$-$^{15}N$ HSQC spectra from multipoint titrations of $^{15}N$-labelled GRB2 with SOS-S9 PRM.
8. **Supplementary Figure S8.** Overlays of 2D $^{1}H$-$^{15}N$ HSQC spectra from multipoint titrations of $^{15}N$-labelled GRB2 with SOS-S10 PRM.
9. **Supplementary Figure S9.** Plots of relative change in chemical shift perturbation (CSP) of backbone $^{1}H^{N}$ and $^{15}N$ nuclei of GRB2 upon the titration with S4, S5, S9, and S10 PRMs.
10. **Supplementary Figure S10.** Residual errors after the bootstrap calculations.
11. **Supplementary Figure S11.** Chemical shift perturbation profile mapped on each residue, against the concentration of ligand with SOS S4 PRM.
12. **Supplementary Figure S12.** Chemical shift perturbation profile mapped on each residue, against the concentration of ligand with SOS S5 PRM.
13. **Supplementary Figure S13.** Chemical shift perturbation profile mapped on each residue, against the concentration of ligand with SOS S9 PRM.
14. **Supplementary Figure S14.** Chemical shift perturbation profile mapped on each residue, against the concentration of ligand with SOS S10 PRM.
15. **Supplementary Figure S15.** ITC analysis for the binding of the full length GRB2 to S2,

S4, S5, S6, S8 S9, and S10 PRMs..

16. **Supplementary Figure S16.** O ;verlays of 2D $^{1}H$-$^{15}N$ HSQC spectra from multipoint titrations of $^{15}N$-labelled GRB2 exclusively with the EGFR phosphorylated peptide.
17. **Supplementary Figure S17.** Overlays of 2D $^{1}H$-$^{15}N$ HSQC spectra from multipoint titrations of $^{15}N$-labelled GRB2 with the EGFR phosphorylated peptide, in the presence of S4 PRM.
18. **Supplementary Table S1** Dissociation constants derived from ITC analysis for the full length GRB2 and isolated SH3 domains against SOS1 PRMs.

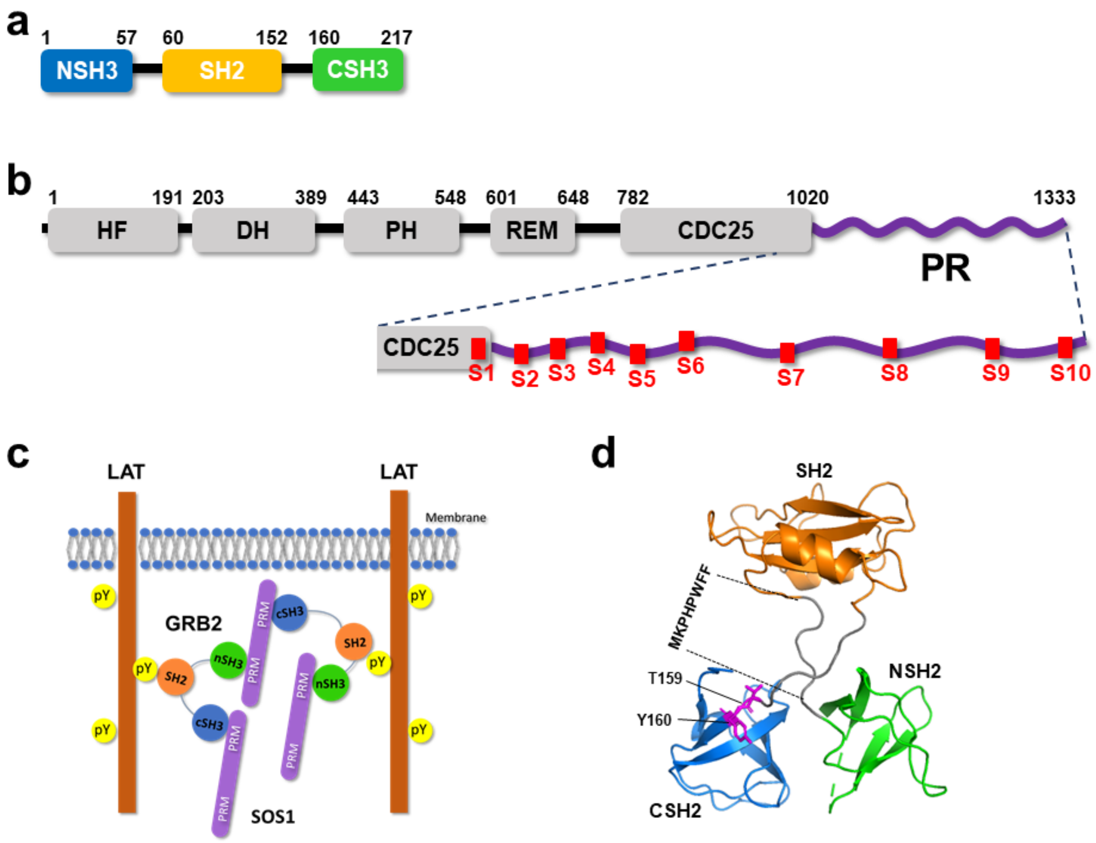


**Figure S1. Domain and three dimensional (3D) structures of GRB2 and SOS1, and schematic illustration of LLPS.** (a) the domain structure of GRB2. (b) the domain structure of SOS1 with the positions of proline-rich motifs, SOS1 S1-S10, indicated by red boxes. (c) Schematic illustration of the considered multivalent interactions among GRB2, SOS1 and LAT in liquid-liquid phase separation (LLPS), proposed in the previous reports. (d) The crystal structure of GRB2.

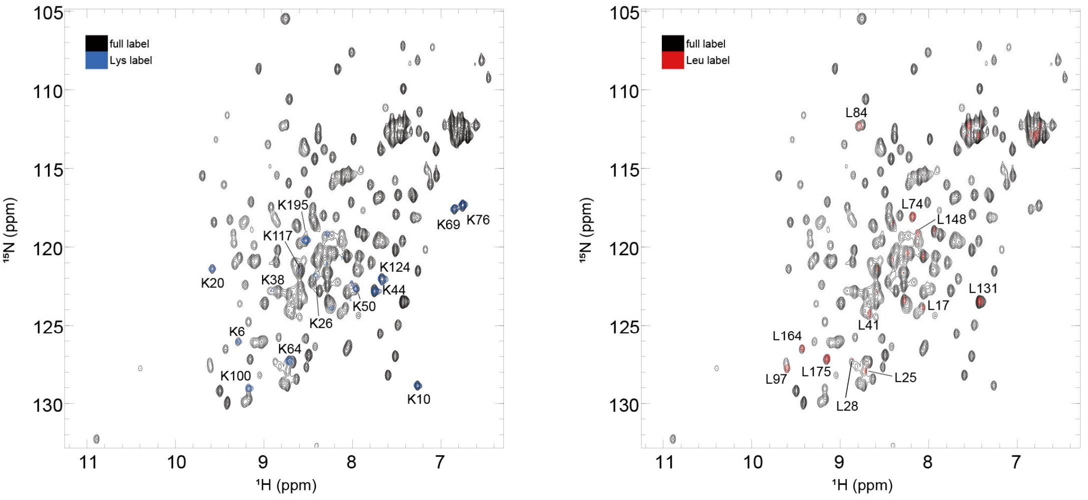


**Figure S2. Overlays of 2D $^{1}$H-$^{15}$N HSQC spectra of $^{15}$N-uniformly and amino-acid-selectively labelled samples.** The spectra of lysine- (left) and leucine- (right) selectively labelled GRB2 were overlaid onto those of the $^{15}$N-uniformly labelled samples.

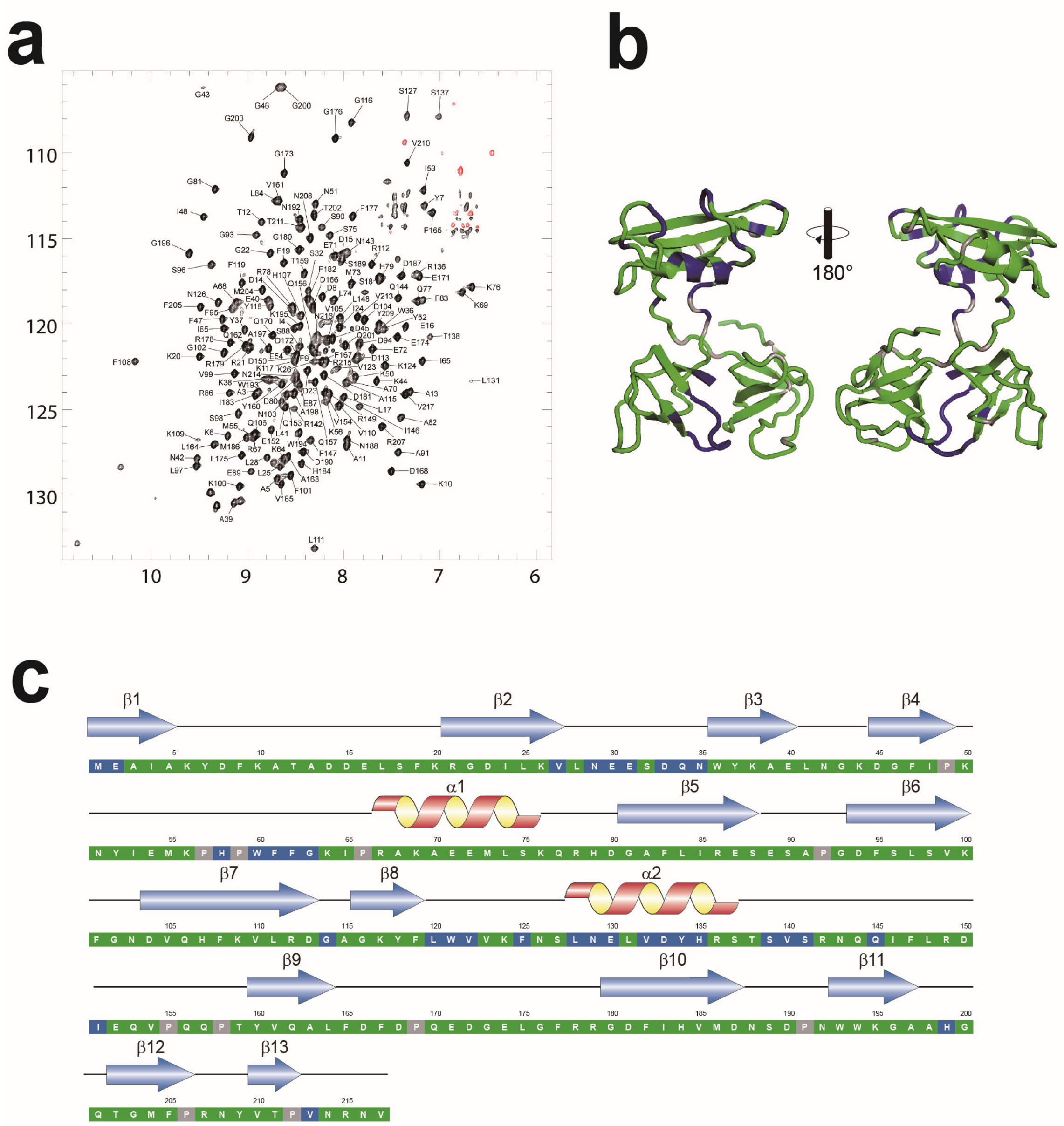


**Figure S3. Backbone resonance assignments of GRB2.** (a) 2D $^{1}$H-$^{15}$N HSQC spectrum and backbone resonance assignments of GRB2. Positive and negative signals (aliased on the $^{15}$N axis) are color-coded in black and red, respectively. (b) The backbone resonance assignments are mapped onto the crystal structure of GRB2. Assigned and unassigned residues are colour-coded in green and blue, respectively. (c) The backbone resonance assignments aligned with secondary structures. Assigned, unassigned, and proline residues are colour-coded in green, blue, and grey, respectively.

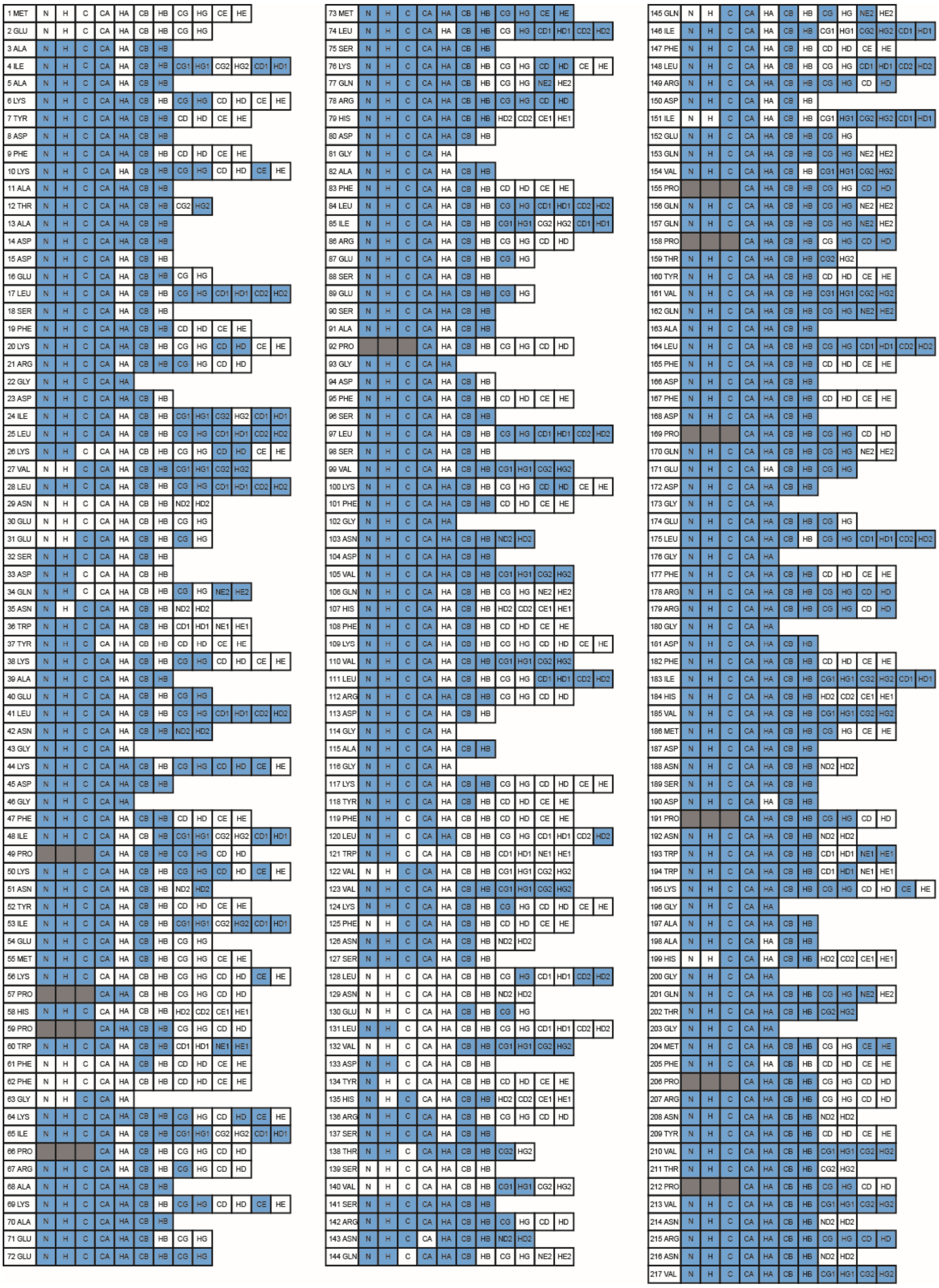

**Figure S4. Table summarising the backbone and side-chain resonance assignments of GRB2.** Blue and white columns indicate assigned and unassigned atoms, respectively.

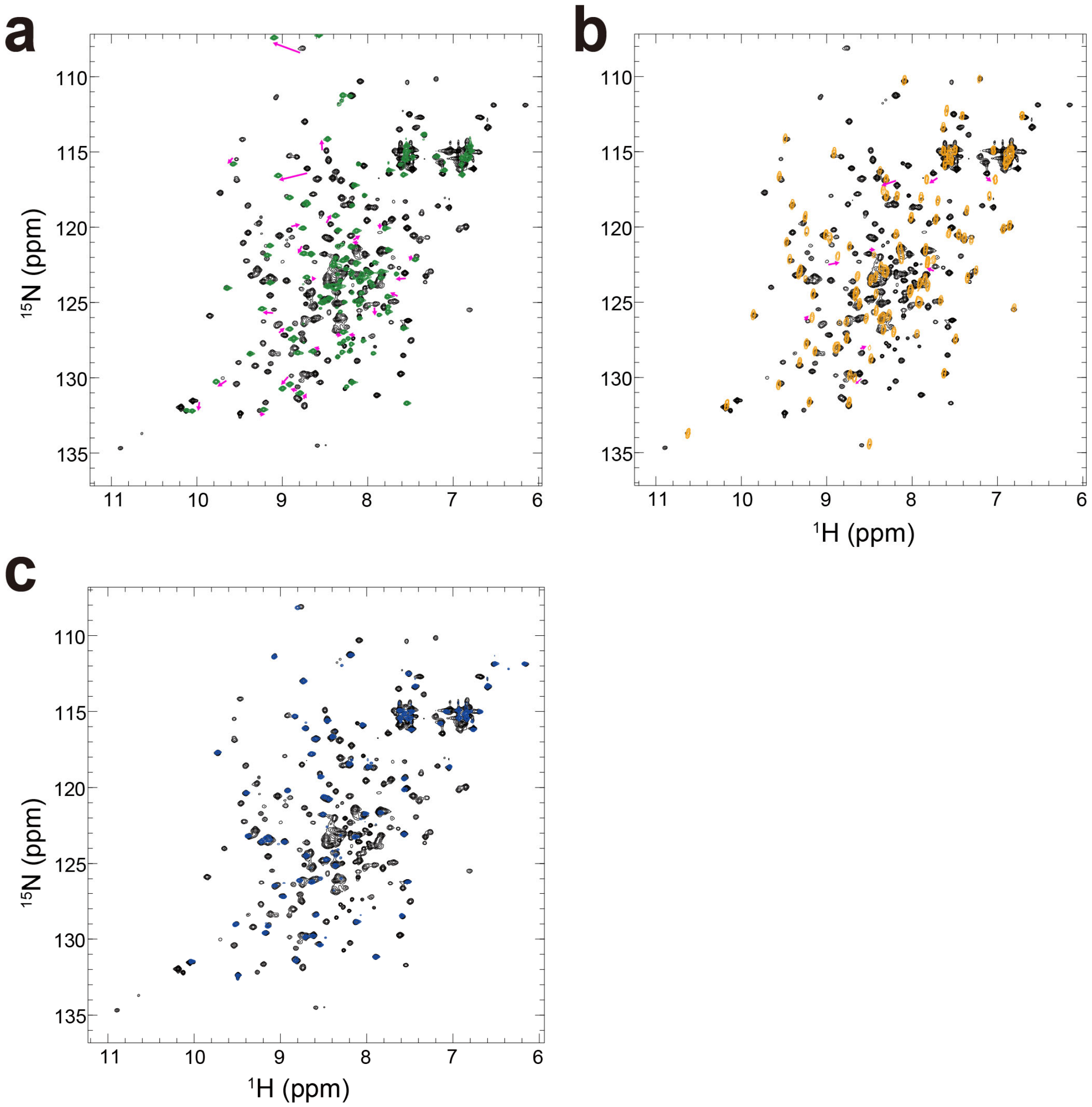


**Figure S5. Overlays of 2D $^{1}$H-$^{15}$N HSQC spectra of the full-length GRB2 and each isolated domain.** The black spectra indicate the full-length GRB2, and green, orange, and blue represent isolated NSH3, SH2, CSH3, respectively. The red arrows indicate representative peaks that significant chemical shift changes were observed.

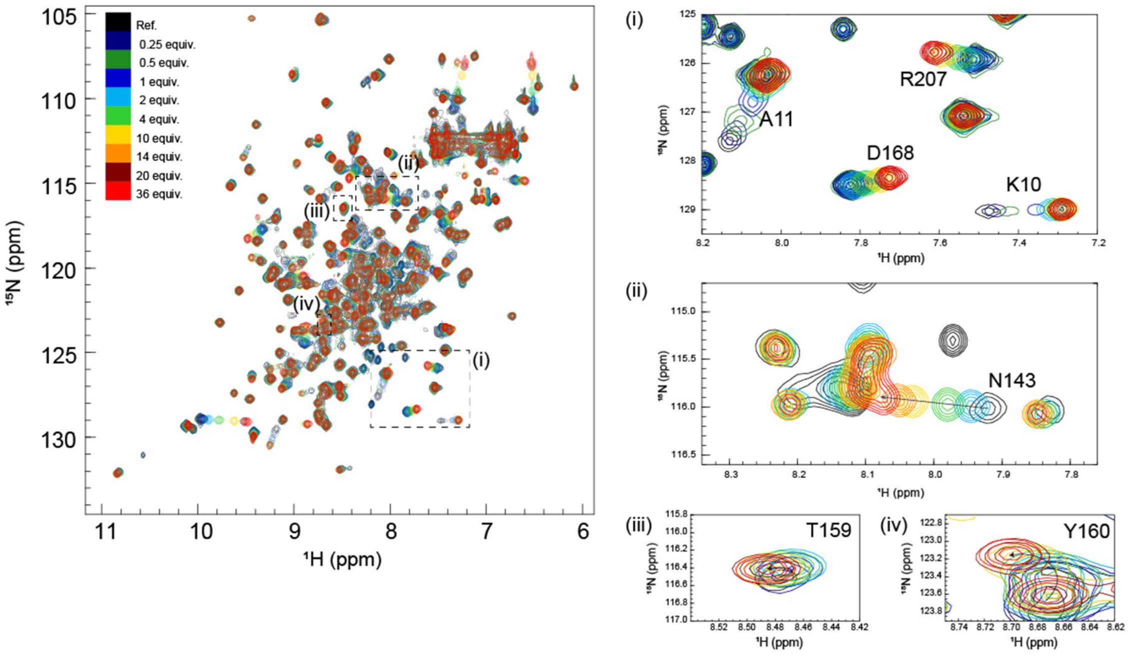


**Figure S6. Overlays of 2D $^{1}$H-$^{15}$N HSQC spectra from multipoint titrations of $^{15}$N-labelled GRB2 with SOS-S5 PRM (DSPPAOPPRQPT).** The peptide concentration was increased stepwise (the protein: peptide molar ratio of 1:0.25, 1:0.5, 1:1, 1:2, 1:4, 1:10, 1:14, 1:20, and 1:36). In this figure, the colour codes of $^{1}$H-$^{15}$N correlation cross-peaks at each titration point, showing the molar ratio of GRB2: SOS-S5, are as follows: black (1:0); dark blue (1:0.25); dark green (1:0.5); blue (1:1); cyan (1:2); green (1:4); yellow (1:10); orange (1:14); dark red (1:20), red (1:36). Cross peaks that showed large chemical shift changes were annotated.

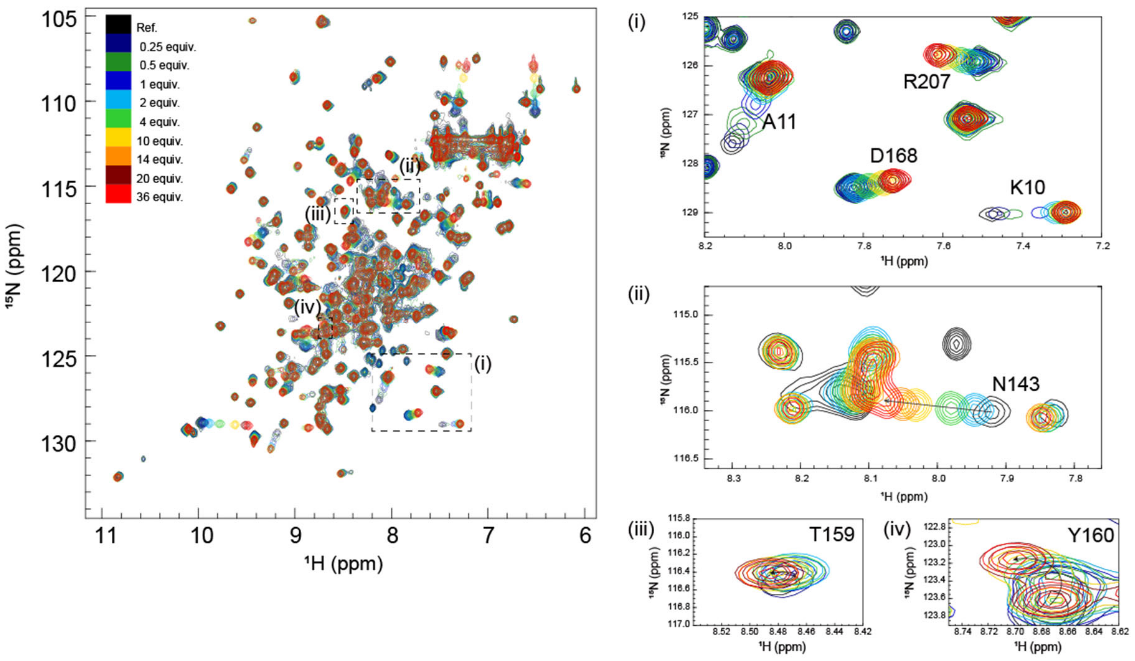


**Figure S7. Overlays of 2D $^{1}$H-$^{15}$N HSQC spectra from multipoint titrations of $^{15}$N-labelled GRB2 with SOS-S9 PRM (IAGPPVPPRQST).** The peptide concentration was increased stepwise (the protein: peptide molar ratio of 1:0.25, 1:0.5, 1:1, 1:2, 1:4, 1:10, 1:14, 1:20, and 1:36). In this figure, the colour codes of $^{1}$H-$^{15}$N correlation cross-peaks at each titration point, showing the molar ratio of GRB2: SOS-S9, are as follows: black (1:0); dark blue (1:0.25); dark green (1:0.5); blue (1:1); cyan (1:2); green (1:4); yellow (1:10); orange (1:14); dark red (1:20), red (1:36). Cross peaks that showed large chemical shift changes were annotated.

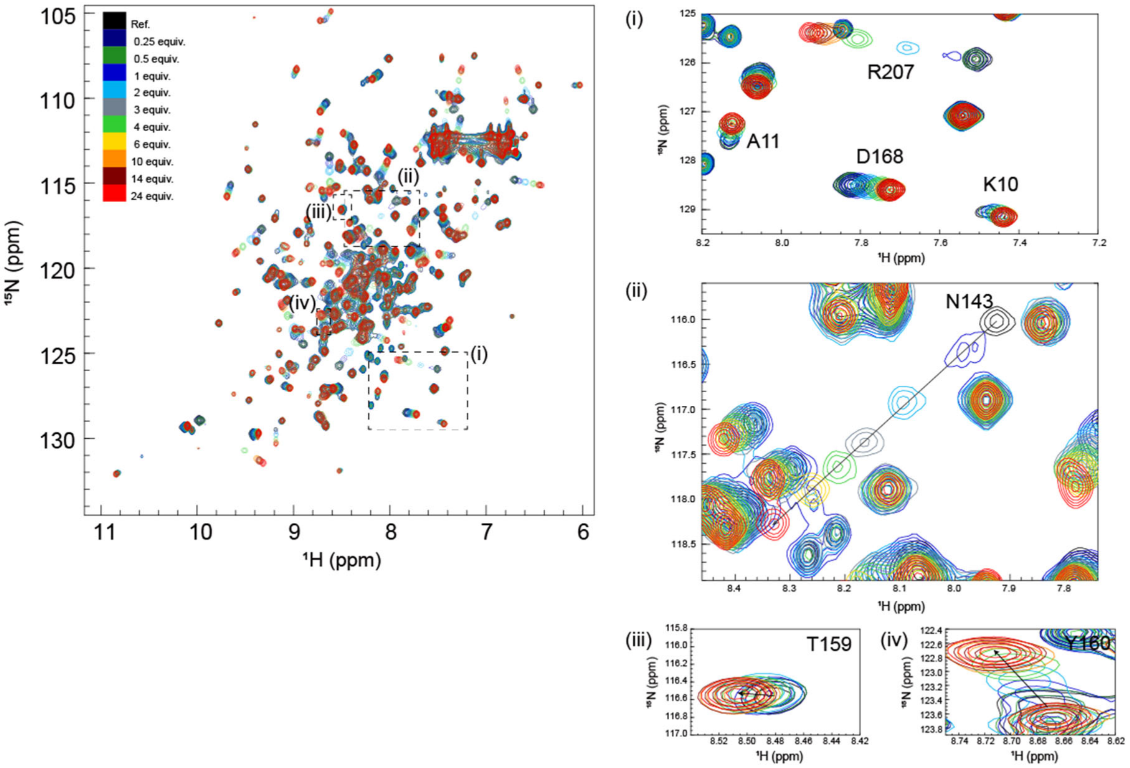


**Figure S8. Overlays of 2D $^{1}$H-$^{15}$N HSQC spectra from multipoint titrations of $^{15}$N-labelled GRB2 with SOS-S10 PRM (PKLPPKTYKREH).** The peptide concentration was increased stepwise (the protein: peptide molar ratio of 1:0.25, 1:0.5, 1:1, 1:2, 1:3, 1:4, 1:6, 1:10, 1:14, and 1:24). In this figure, the colour codes of $^{1}$H-$^{15}$N correlation cross-peaks at each titration point, showing the molar ratio of GRB2: SOS-S10, are as follows: black (1:0); dark blue (1:0.25); dark green (1:0.5); blue (1:1); cyan (1:2); grey (1:3), green (1:4); yellow (1:6); orange (1:10); dark red (1:14), red (1:24). Cross peaks that showed large chemical shift changes were annotated.

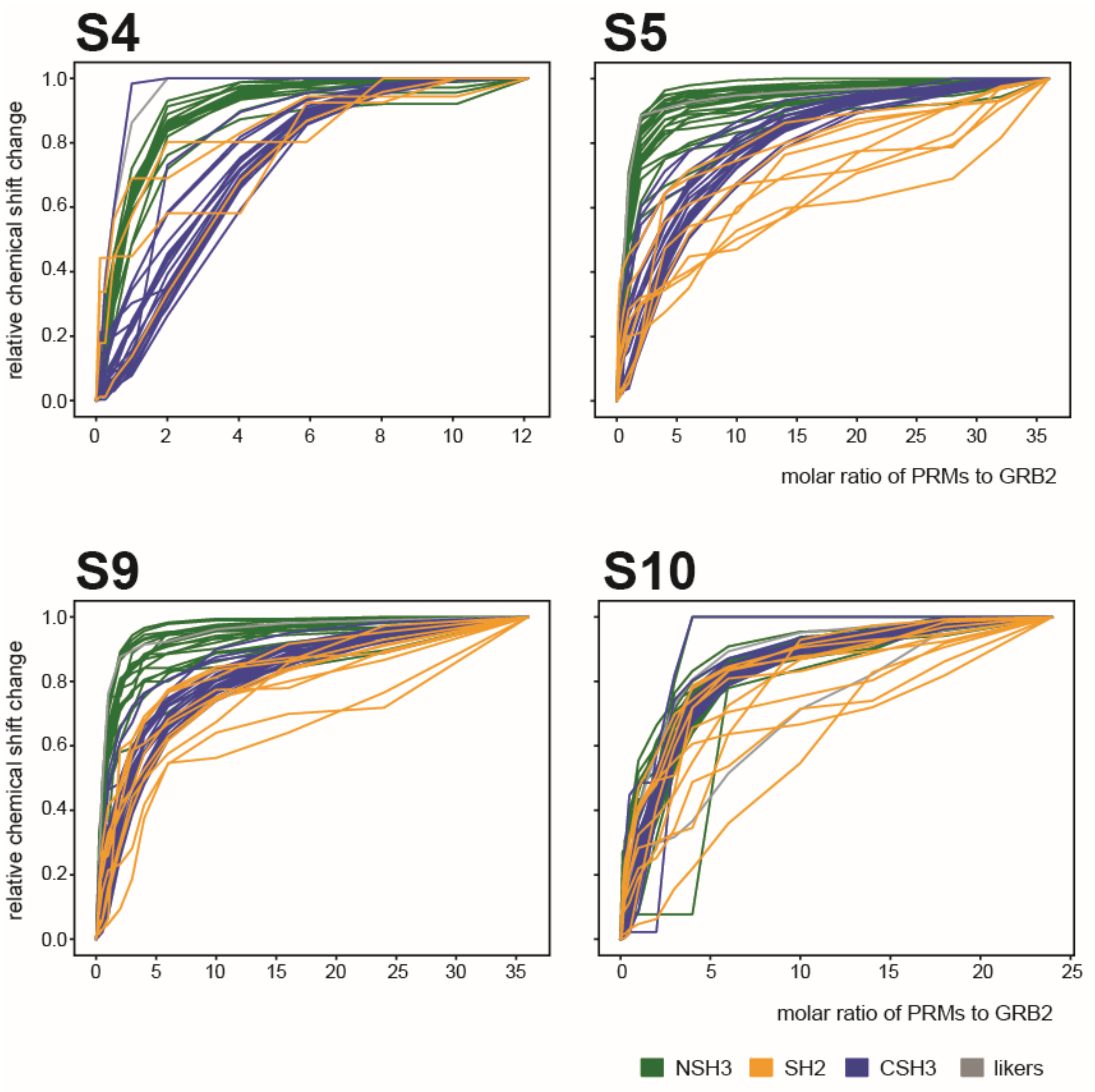


**Figure S9. Plots of relative change in chemical shift perturbation (CSP) of backbone $^{1}H^{N}$ and $^{15}N$ nuclei of GRB2 upon the titration with S4, S5, S9, and S10 PRMs.** A line represents the relative CSP beyond 0.06 ppm for a residue. Lines are colour-coded for each domain and linker.

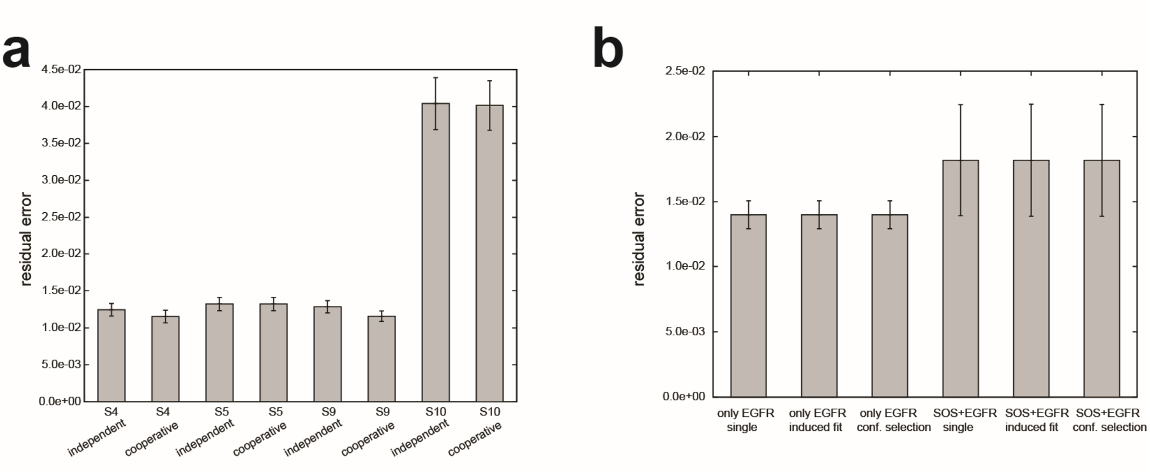


**Figure S10.** **Residual errors after the bootstrap calculations.** Residual errors after the bootstrap calculations with the CSPs of the S4, S5, S9, and S10 PRMs titration experiments for two-independent and two-cooperative binding models. residual errors after the bootstrap calculations for two-independent and two-cooperative binding models (right)

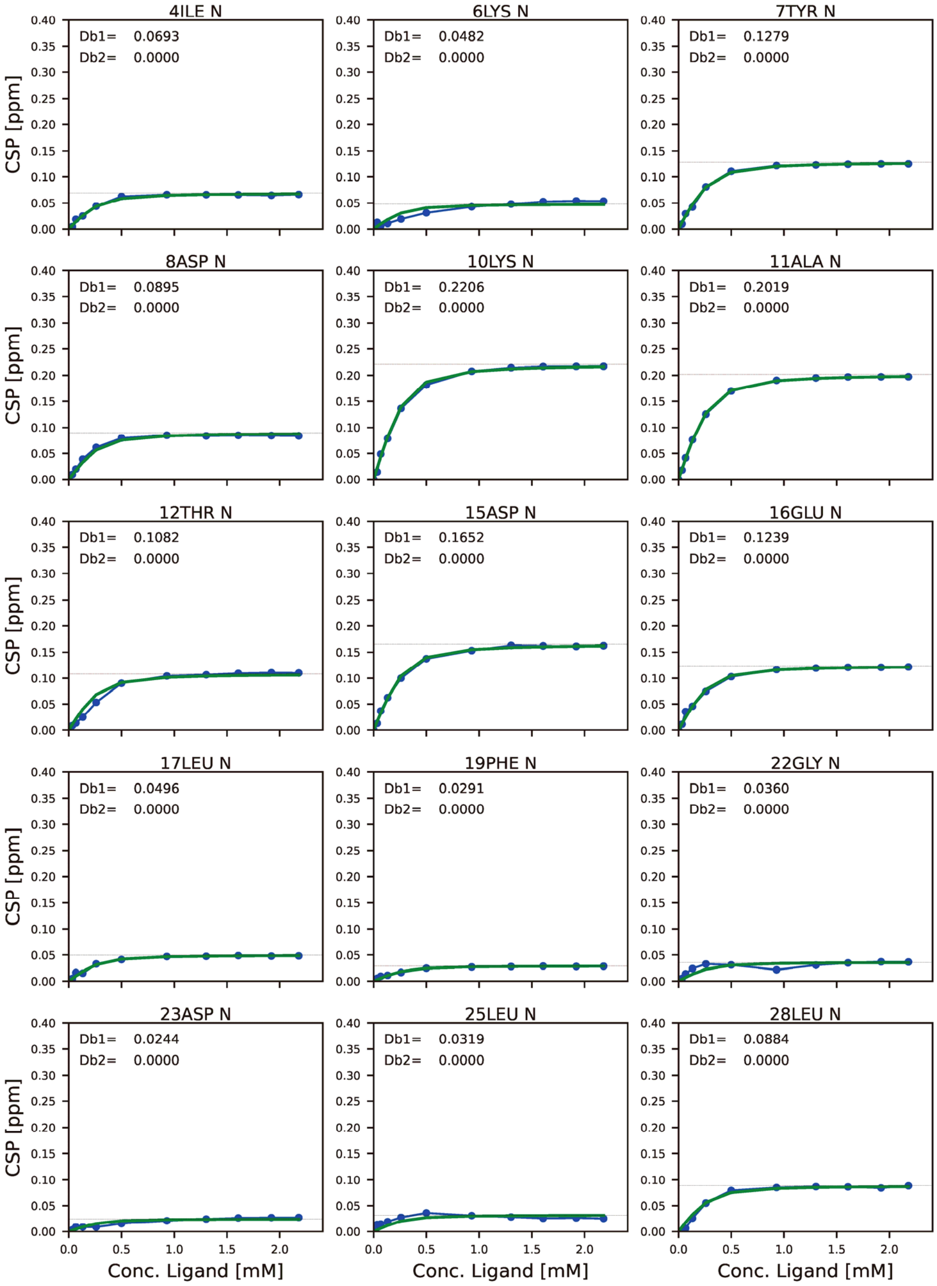
4ILE N
Db1= 0.0693
Db2= 0.0000
6LYS N
Db1= 0.0482
Db2= 0.0000
7TYR N
Db1= 0.1279
Db2= 0.0000
8ASP N
Db1= 0.0895
Db2= 0.0000
10LYS N
Db1= 0.2206
Db2= 0.0000
11ALA N
Db1= 0.2019
Db2= 0.0000
12THR N
Db1= 0.1082
Db2= 0.0000
15ASP N
Db1= 0.1652
Db2= 0.0000
16GLU N
Db1= 0.1239
Db2= 0.0000
17LEU N
Db1= 0.0496
Db2= 0.0000
19PHE N
Db1= 0.0291
Db2= 0.0000
22GLY N
Db1= 0.0360
Db2= 0.0000
23ASP N
Db1= 0.0244
Db2= 0.0000
25LEU N
Db1= 0.0319
Db2= 0.0000
28LEU N
Db1= 0.0884
Db2= 0.0000
CSP [ppm]
Conc. Ligand [mM]

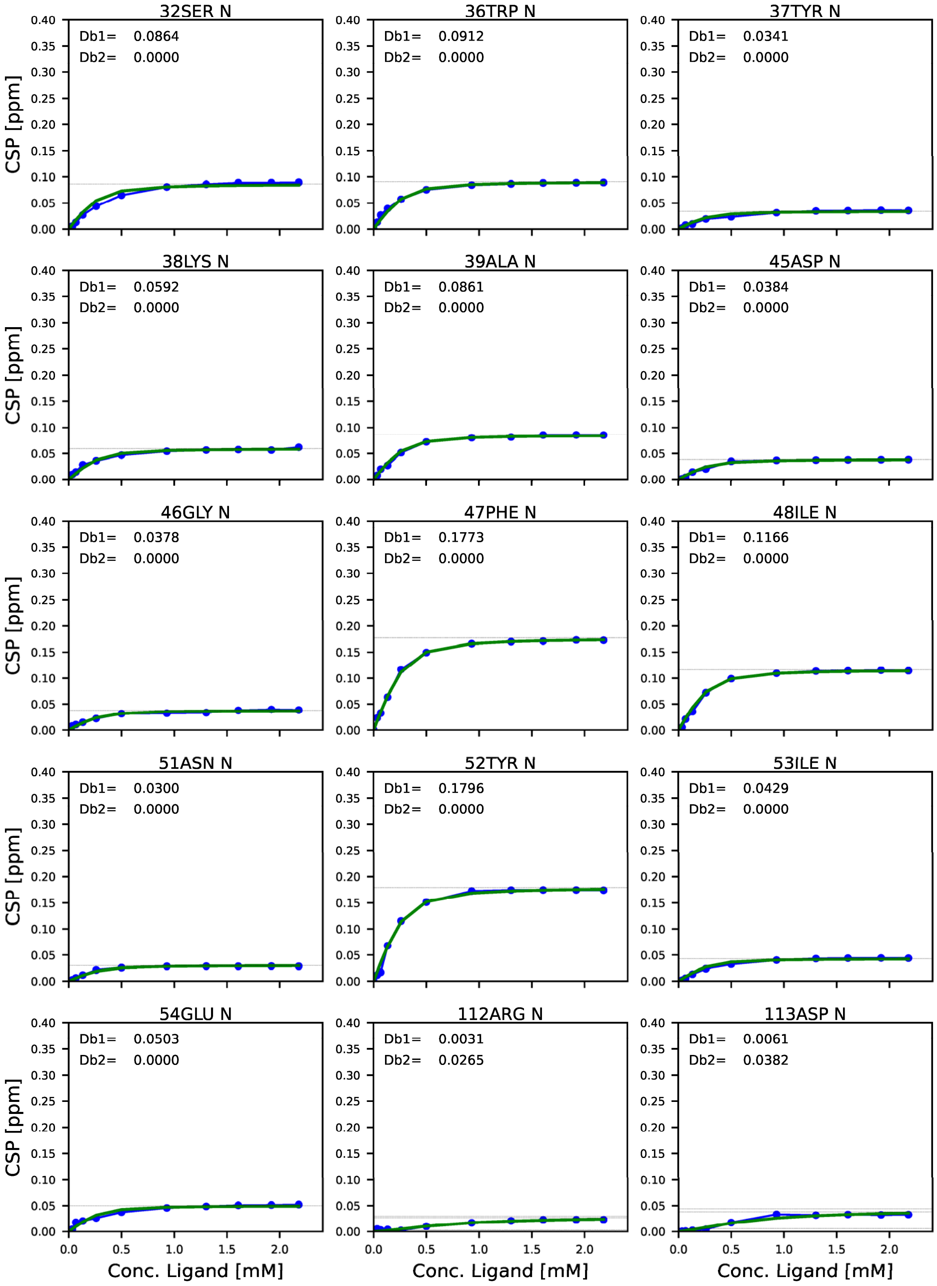

32SER N
Db1= 0.0864
Db2= 0.0000
36TRP N
Db1= 0.0912
Db2= 0.0000
37TYR N
Db1= 0.0341
Db2= 0.0000
38LYS N
Db1= 0.0592
Db2= 0.0000
39ALA N
Db1= 0.0861
Db2= 0.0000
45ASP N
Db1= 0.0384
Db2= 0.0000
46GLY N
Db1= 0.0378
Db2= 0.0000
47PHE N
Db1= 0.1773
Db2= 0.0000
48ILE N
Db1= 0.1166
Db2= 0.0000
51ASN N
Db1= 0.0300
Db2= 0.0000
52TYR N
Db1= 0.1796
Db2= 0.0000
53ILE N
Db1= 0.0429
Db2= 0.0000
54GLU N
Db1= 0.0503
Db2= 0.0000
112ARG N
Db1= 0.0031
Db2= 0.0265
113ASP N
Db1= 0.0061
Db2= 0.0382
CSP [ppm]
Conc. Ligand [mM]

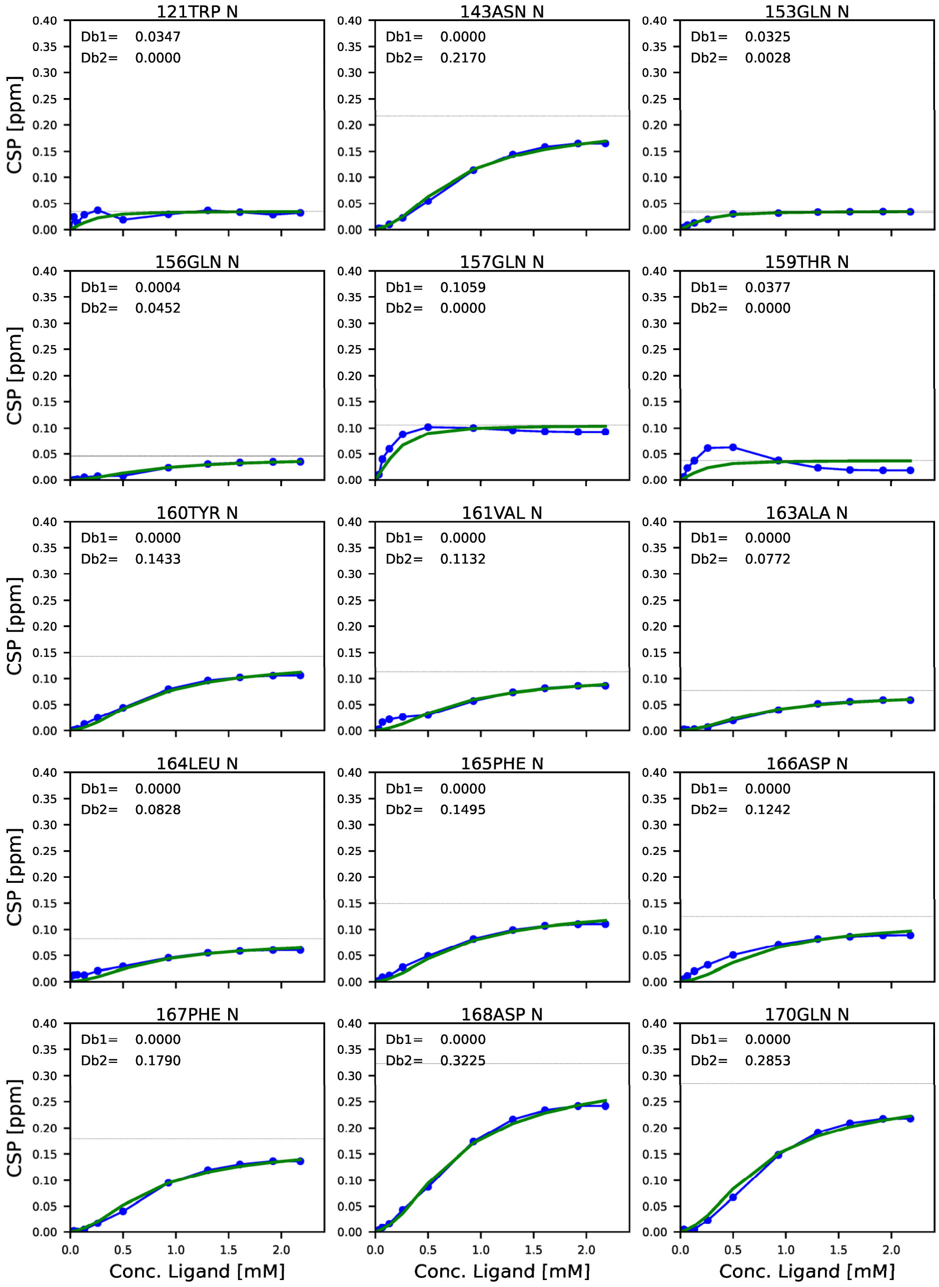
121TRP N
Db1= 0.0347
Db2= 0.0000
143ASN N
Db1= 0.0000
Db2= 0.2170
153GLN N
Db1= 0.0325
Db2= 0.0028
156GLN N
Db1= 0.0004
Db2= 0.0452
157GLN N
Db1= 0.1059
Db2= 0.0000
159THR N
Db1= 0.0377
Db2= 0.0000
160TYR N
Db1= 0.0000
Db2= 0.1433
161VAL N
Db1= 0.0000
Db2= 0.1132
163ALA N
Db1= 0.0000
Db2= 0.0772
164LEU N
Db1= 0.0000
Db2= 0.0828
165PHE N
Db1= 0.0000
Db2= 0.1495
166ASP N
Db1= 0.0000
Db2= 0.1242
167PHE N
Db1= 0.0000
Db2= 0.1790
168ASP N
Db1= 0.0000
Db2= 0.3225
170GLN N
Db1= 0.0000
Db2= 0.2853
CSP [ppm]
Conc. Ligand [mM]

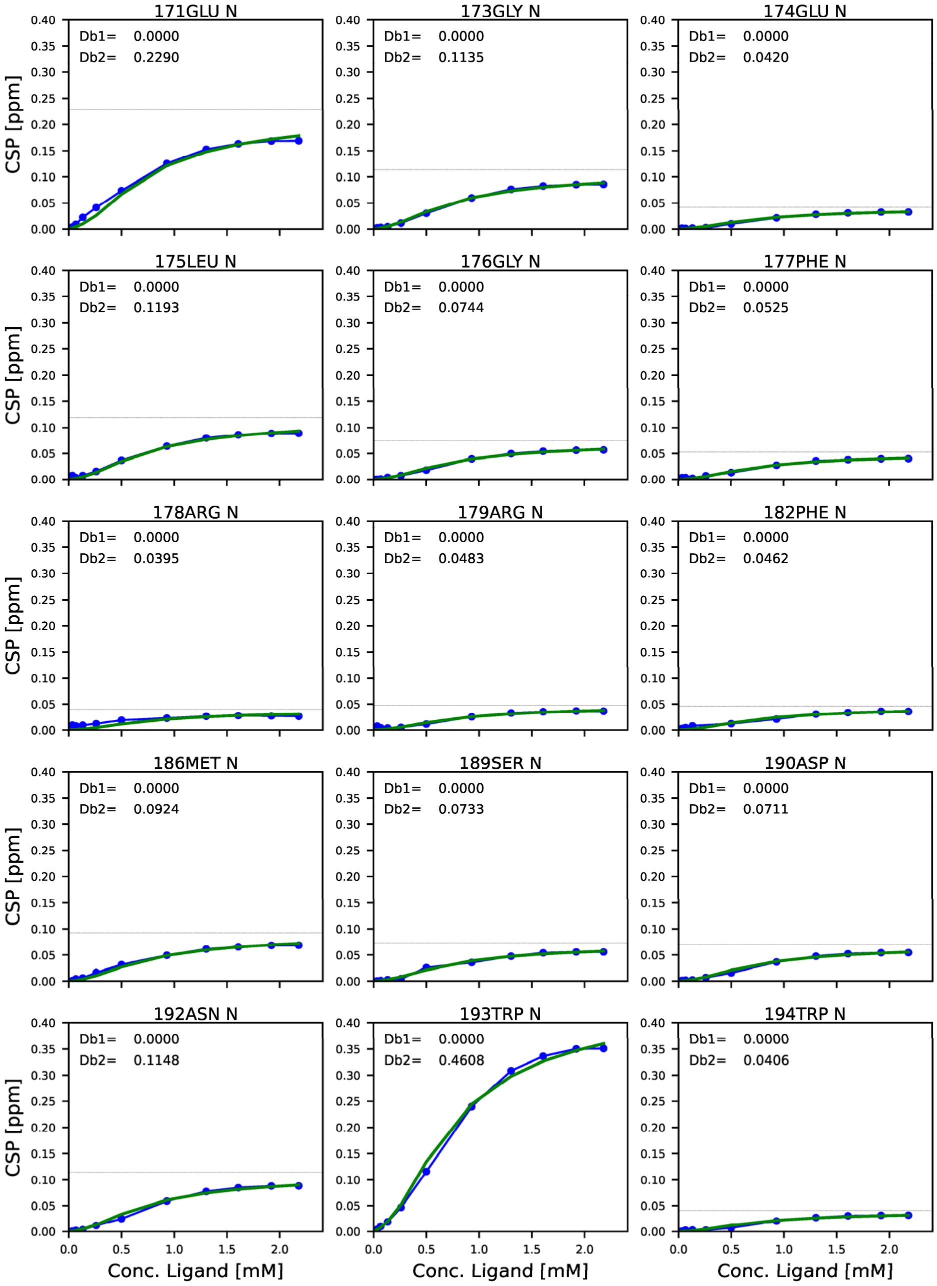
171GLU N
Db1= 0.0000
Db2= 0.2290
173GLY N
Db1= 0.0000
Db2= 0.1135
174GLU N
Db1= 0.0000
Db2= 0.0420
175LEU N
Db1= 0.0000
Db2= 0.1193
176GLY N
Db1= 0.0000
Db2= 0.0744
177PHE N
Db1= 0.0000
Db2= 0.0525
178ARG N
Db1= 0.0000
Db2= 0.0395
179ARG N
Db1= 0.0000
Db2= 0.0483
182PHE N
Db1= 0.0000
Db2= 0.0462
186MET N
Db1= 0.0000
Db2= 0.0924
189SER N
Db1= 0.0000
Db2= 0.0733
190ASP N
Db1= 0.0000
Db2= 0.0711
192ASN N
Db1= 0.0000
Db2= 0.1148
193TRP N
Db1= 0.0000
Db2= 0.4608
194TRP N
Db1= 0.0000
Db2= 0.0406
CSP [ppm]
Conc. Ligand [mM]

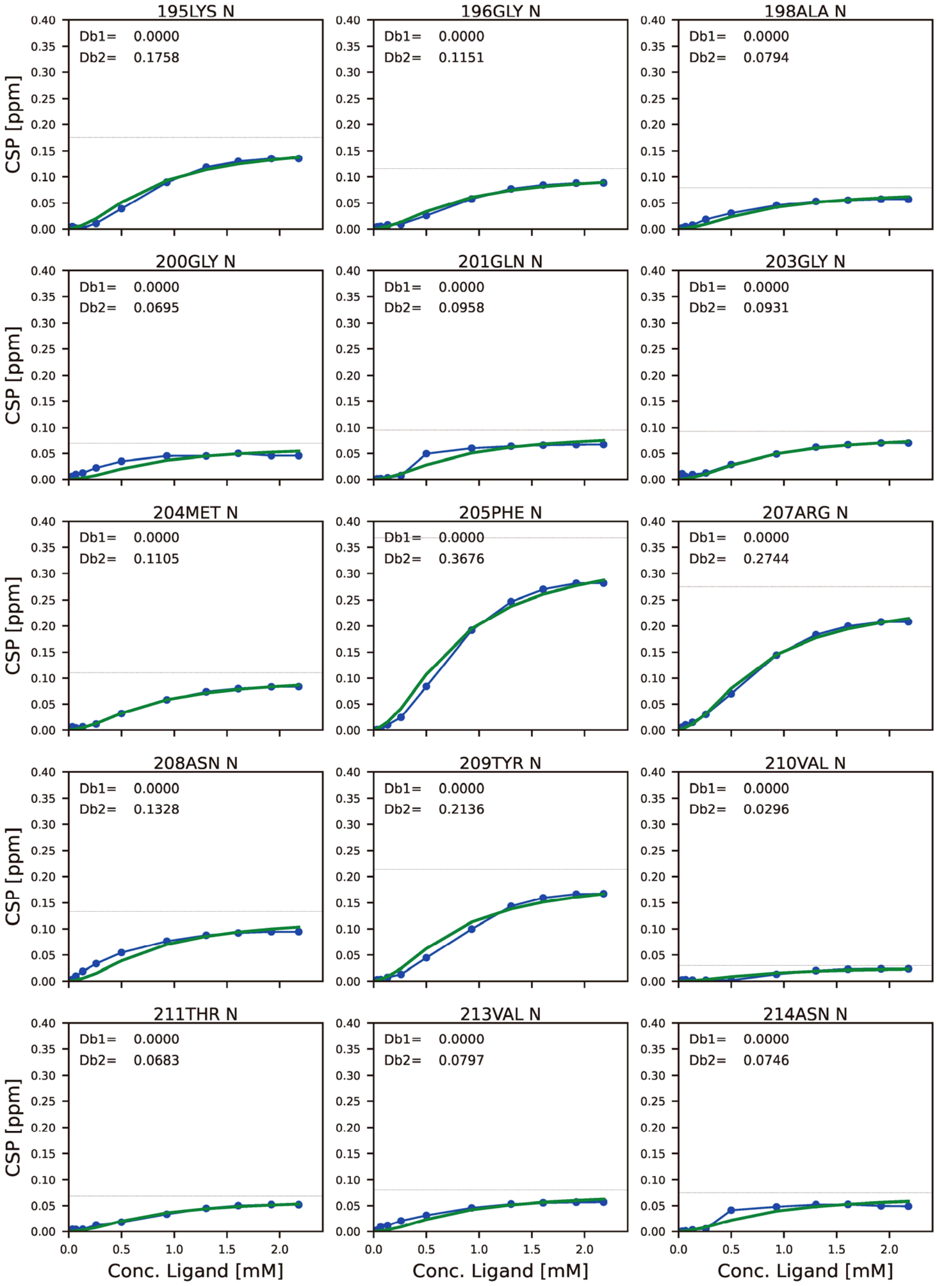
195LYS N
Db1= 0.0000
Db2= 0.1758
196GLY N
Db1= 0.0000
Db2= 0.1151
198ALA N
Db1= 0.0000
Db2= 0.0794
200GLY N
Db1= 0.0000
Db2= 0.0695
201GLN N
Db1= 0.0000
Db2= 0.0958
203GLY N
Db1= 0.0000
Db2= 0.0931
204MET N
Db1= 0.0000
Db2= 0.1105
205PHE N
Db1= 0.0000
Db2= 0.3676
207ARG N
Db1= 0.0000
Db2= 0.2744
208ASN N
Db1= 0.0000
Db2= 0.1328
209TYR N
Db1= 0.0000
Db2= 0.2136
210VAL N
Db1= 0.0000
Db2= 0.0296
211THR N
Db1= 0.0000
Db2= 0.0683
213VAL N
Db1= 0.0000
Db2= 0.0797
214ASN N
Db1= 0.0000
Db2= 0.0746
CSP [ppm]
Conc. Ligand [mM]

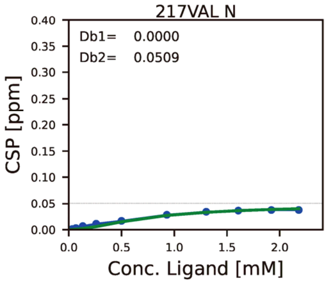


**Figure S11. Chemical shift perturbation profile mapped on each residue, against the concentration of ligand with SOS S4 PRM.**

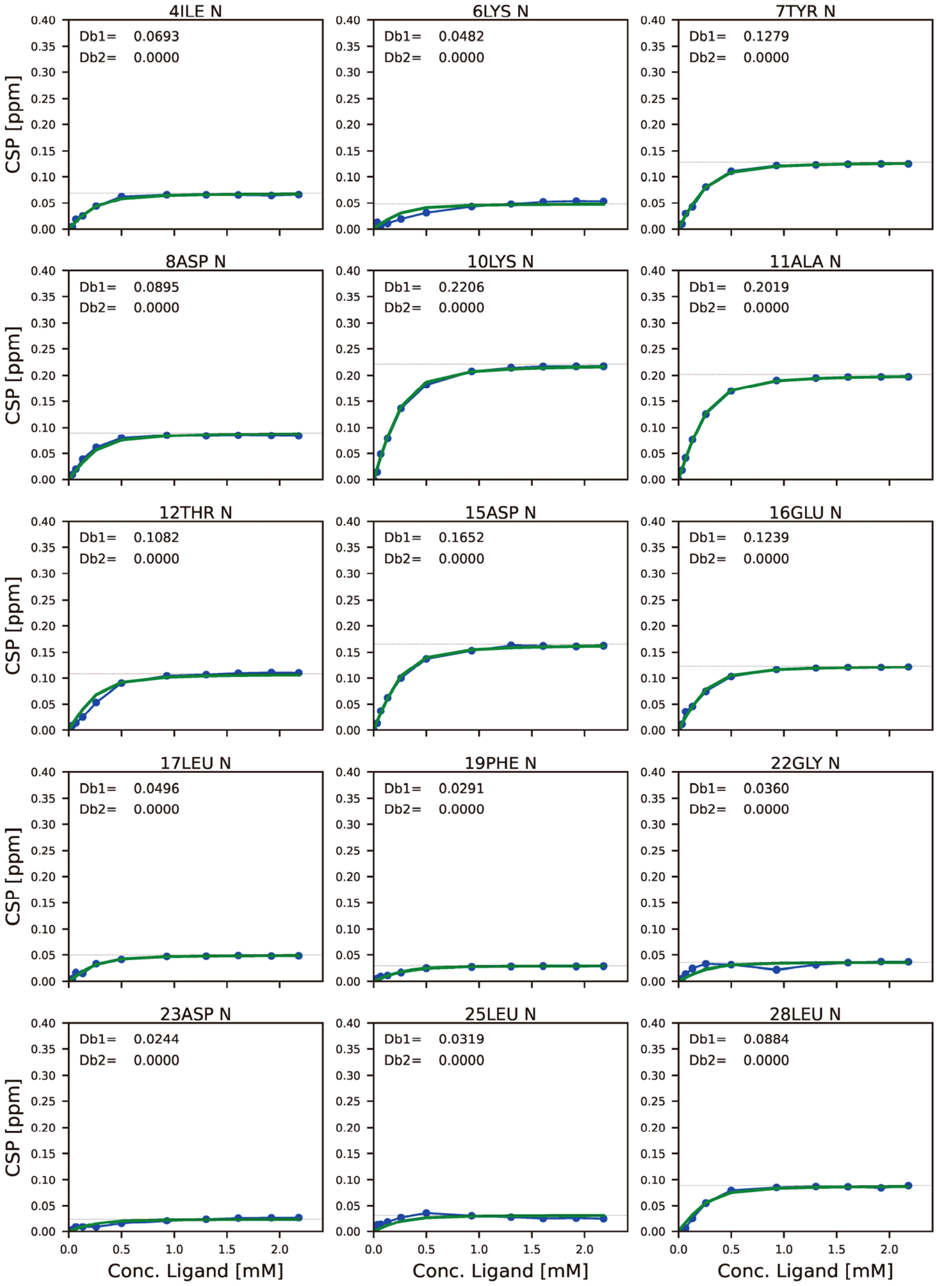
4ILE N
Db1= 0.0693
Db2= 0.0000
6LYS N
Db1= 0.0482
Db2= 0.0000
7TYR N
Db1= 0.1279
Db2= 0.0000
8ASP N
Db1= 0.0895
Db2= 0.0000
10LYS N
Db1= 0.2206
Db2= 0.0000
11ALA N
Db1= 0.2019
Db2= 0.0000
12THR N
Db1= 0.1082
Db2= 0.0000
15ASP N
Db1= 0.1652
Db2= 0.0000
16GLU N
Db1= 0.1239
Db2= 0.0000
17LEU N
Db1= 0.0496
Db2= 0.0000
19PHE N
Db1= 0.0291
Db2= 0.0000
22GLY N
Db1= 0.0360
Db2= 0.0000
23ASP N
Db1= 0.0244
Db2= 0.0000
25LEU N
Db1= 0.0319
Db2= 0.0000
28LEU N
Db1= 0.0884
Db2= 0.0000
CSP [ppm]
Conc. Ligand [mM]

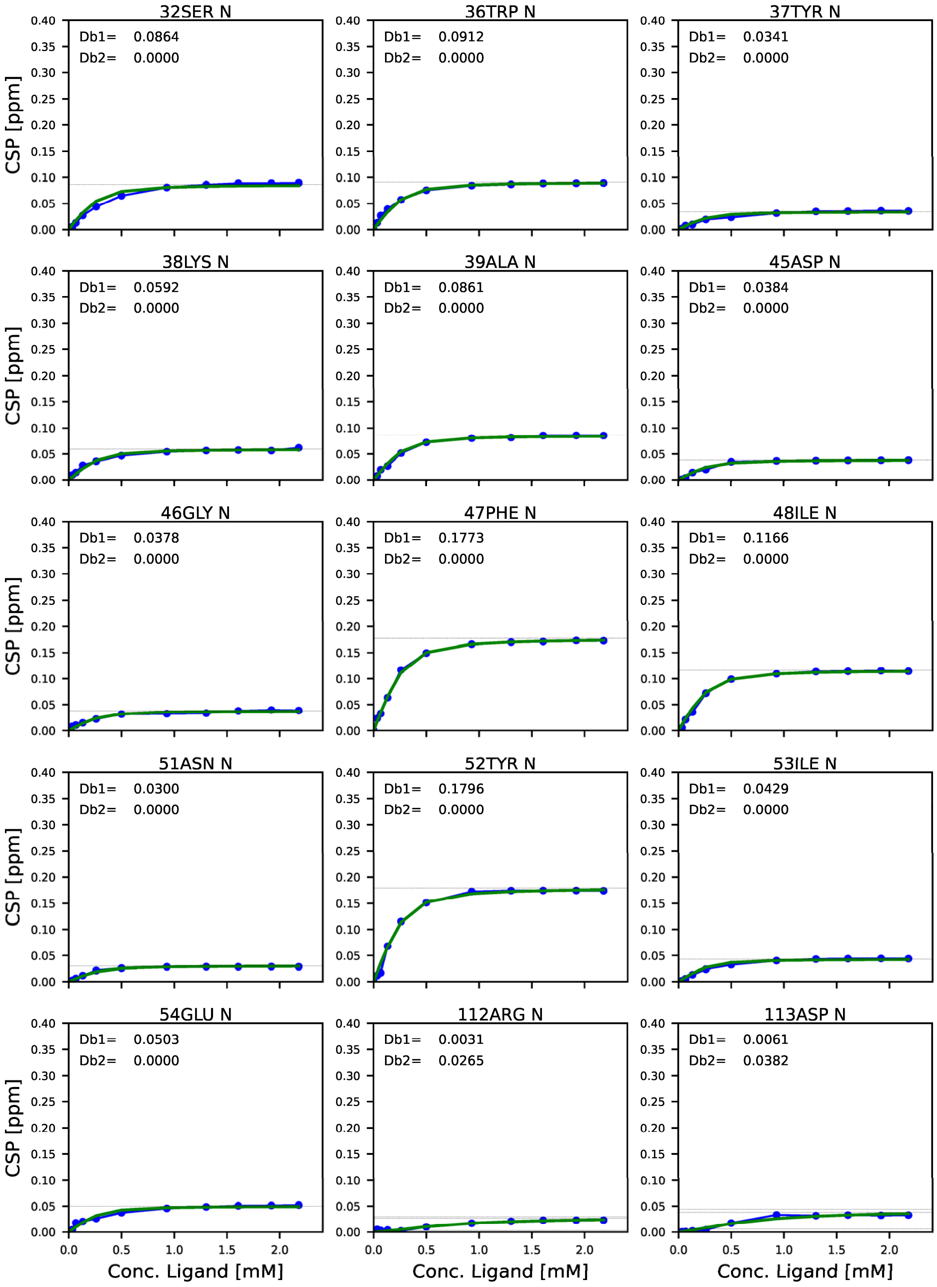
32SER N
Db1= 0.0864
Db2= 0.0000
36TRP N
Db1= 0.0912
Db2= 0.0000
37TYR N
Db1= 0.0341
Db2= 0.0000
38LYS N
Db1= 0.0592
Db2= 0.0000
39ALA N
Db1= 0.0861
Db2= 0.0000
45ASP N
Db1= 0.0384
Db2= 0.0000
46GLY N
Db1= 0.0378
Db2= 0.0000
47PHE N
Db1= 0.1773
Db2= 0.0000
48ILE N
Db1= 0.1166
Db2= 0.0000
51ASN N
Db1= 0.0300
Db2= 0.0000
52TYR N
Db1= 0.1796
Db2= 0.0000
53ILE N
Db1= 0.0429
Db2= 0.0000
54GLU N
Db1= 0.0503
Db2= 0.0000
112ARG N
Db1= 0.0031
Db2= 0.0265
113ASP N
Db1= 0.0061
Db2= 0.0382
CSP [ppm]
Conc. Ligand [mM]

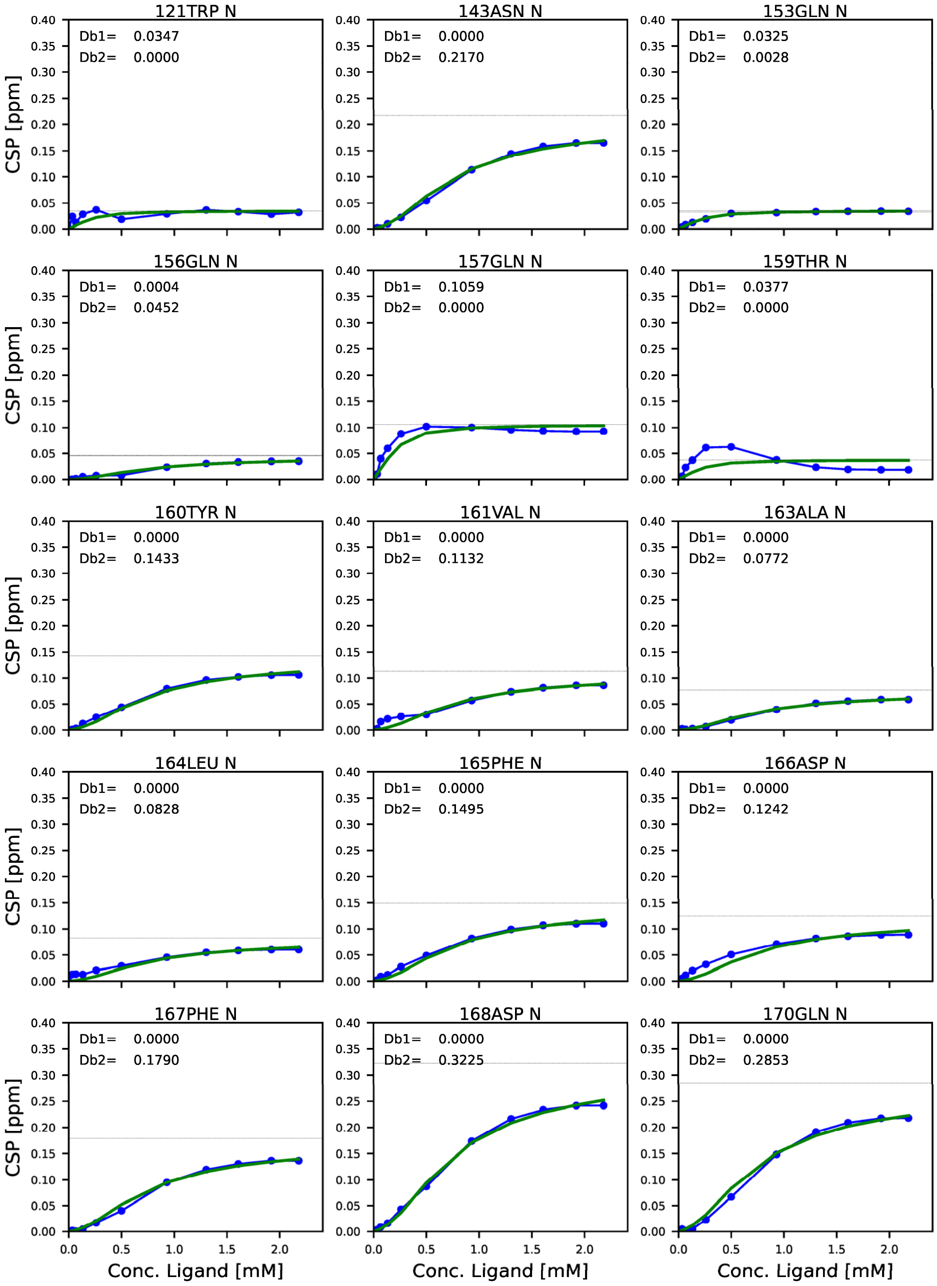
121TRP N
Db1= 0.0347
Db2= 0.0000
143ASN N
Db1= 0.0000
Db2= 0.2170
153GLN N
Db1= 0.0325
Db2= 0.0028
156GLN N
Db1= 0.0004
Db2= 0.0452
157GLN N
Db1= 0.1059
Db2= 0.0000
159THR N
Db1= 0.0377
Db2= 0.0000
160TYR N
Db1= 0.0000
Db2= 0.1433
161VAL N
Db1= 0.0000
Db2= 0.1132
163ALA N
Db1= 0.0000
Db2= 0.0772
164LEU N
Db1= 0.0000
Db2= 0.0828
165PHE N
Db1= 0.0000
Db2= 0.1495
166ASP N
Db1= 0.0000
Db2= 0.1242
167PHE N
Db1= 0.0000
Db2= 0.1790
168ASP N
Db1= 0.0000
Db2= 0.3225
170GLN N
Db1= 0.0000
Db2= 0.2853
CSP [ppm]
Conc. Ligand [mM]

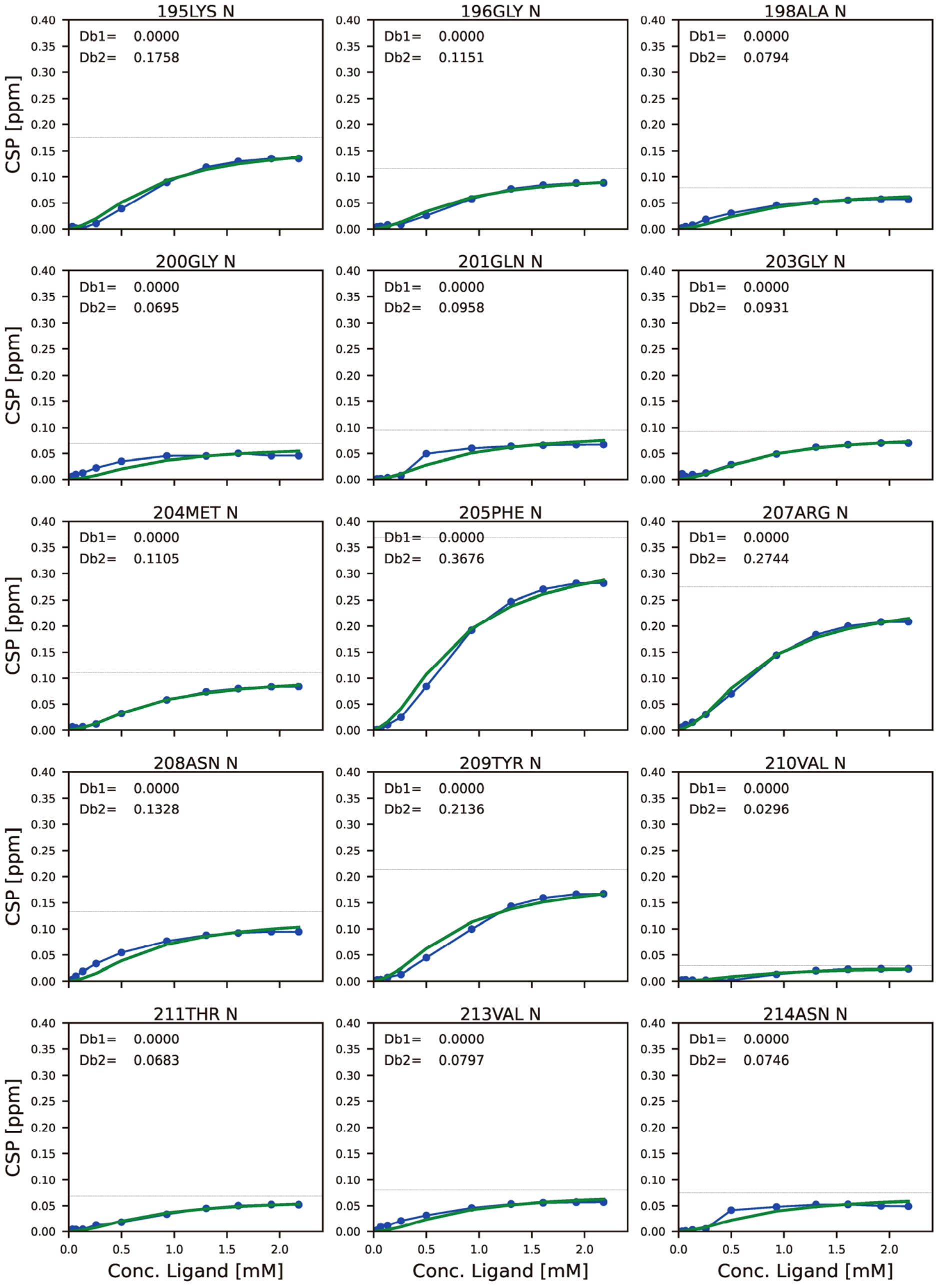
195LYS N
Db1= 0.0000
Db2= 0.1758
196GLY N
Db1= 0.0000
Db2= 0.1151
198ALA N
Db1= 0.0000
Db2= 0.0794
200GLY N
Db1= 0.0000
Db2= 0.0695
201GLN N
Db1= 0.0000
Db2= 0.0958
203GLY N
Db1= 0.0000
Db2= 0.0931
204MET N
Db1= 0.0000
Db2= 0.1105
205PHE N
Db1= 0.0000
Db2= 0.3676
207ARG N
Db1= 0.0000
Db2= 0.2744
208ASN N
Db1= 0.0000
Db2= 0.1328
209TYR N
Db1= 0.0000
Db2= 0.2136
210VAL N
Db1= 0.0000
Db2= 0.0296
211THR N
Db1= 0.0000
Db2= 0.0683
213VAL N
Db1= 0.0000
Db2= 0.0797
214ASN N
Db1= 0.0000
Db2= 0.0746
CSP [ppm]
Conc. Ligand [mM]

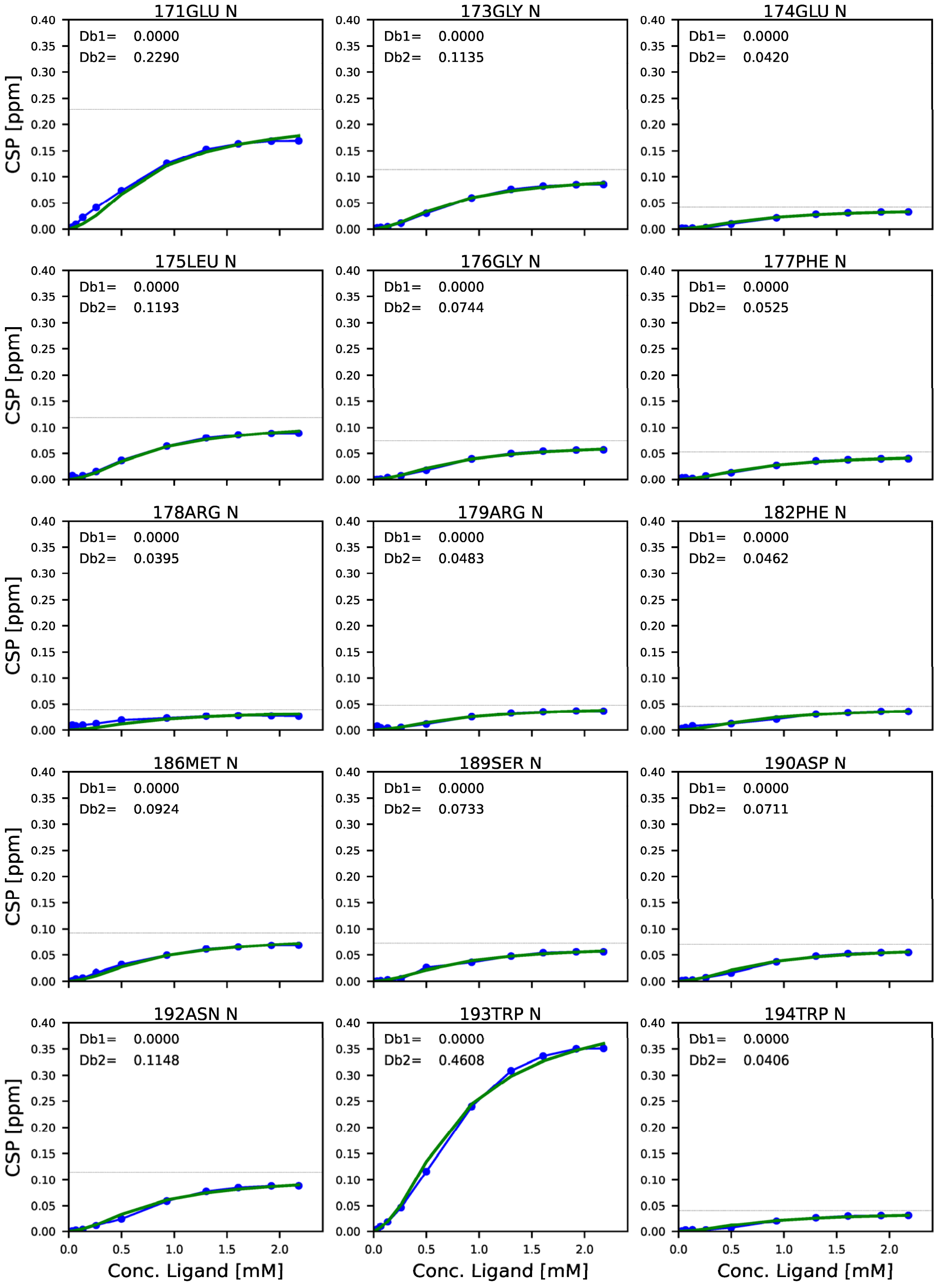
171GLU N
Db1= 0.0000
Db2= 0.2290
173GLY N
Db1= 0.0000
Db2= 0.1135
174GLU N
Db1= 0.0000
Db2= 0.0420
175LEU N
Db1= 0.0000
Db2= 0.1193
176GLY N
Db1= 0.0000
Db2= 0.0744
177PHE N
Db1= 0.0000
Db2= 0.0525
178ARG N
Db1= 0.0000
Db2= 0.0395
179ARG N
Db1= 0.0000
Db2= 0.0483
182PHE N
Db1= 0.0000
Db2= 0.0462
186MET N
Db1= 0.0000
Db2= 0.0924
189SER N
Db1= 0.0000
Db2= 0.0733
190ASP N
Db1= 0.0000
Db2= 0.0711
192ASN N
Db1= 0.0000
Db2= 0.1148
193TRP N
Db1= 0.0000
Db2= 0.4608
194TRP N
Db1= 0.0000
Db2= 0.0406
CSP [ppm]
Conc. Ligand [mM]

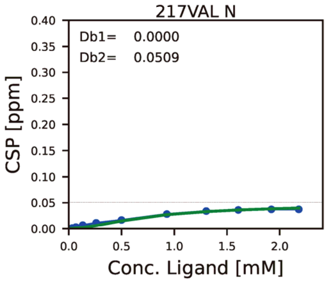


**Figure S12. Chemical shift perturbation profile mapped on each residue, against the concentration of ligand with SOS S5 PRM.**

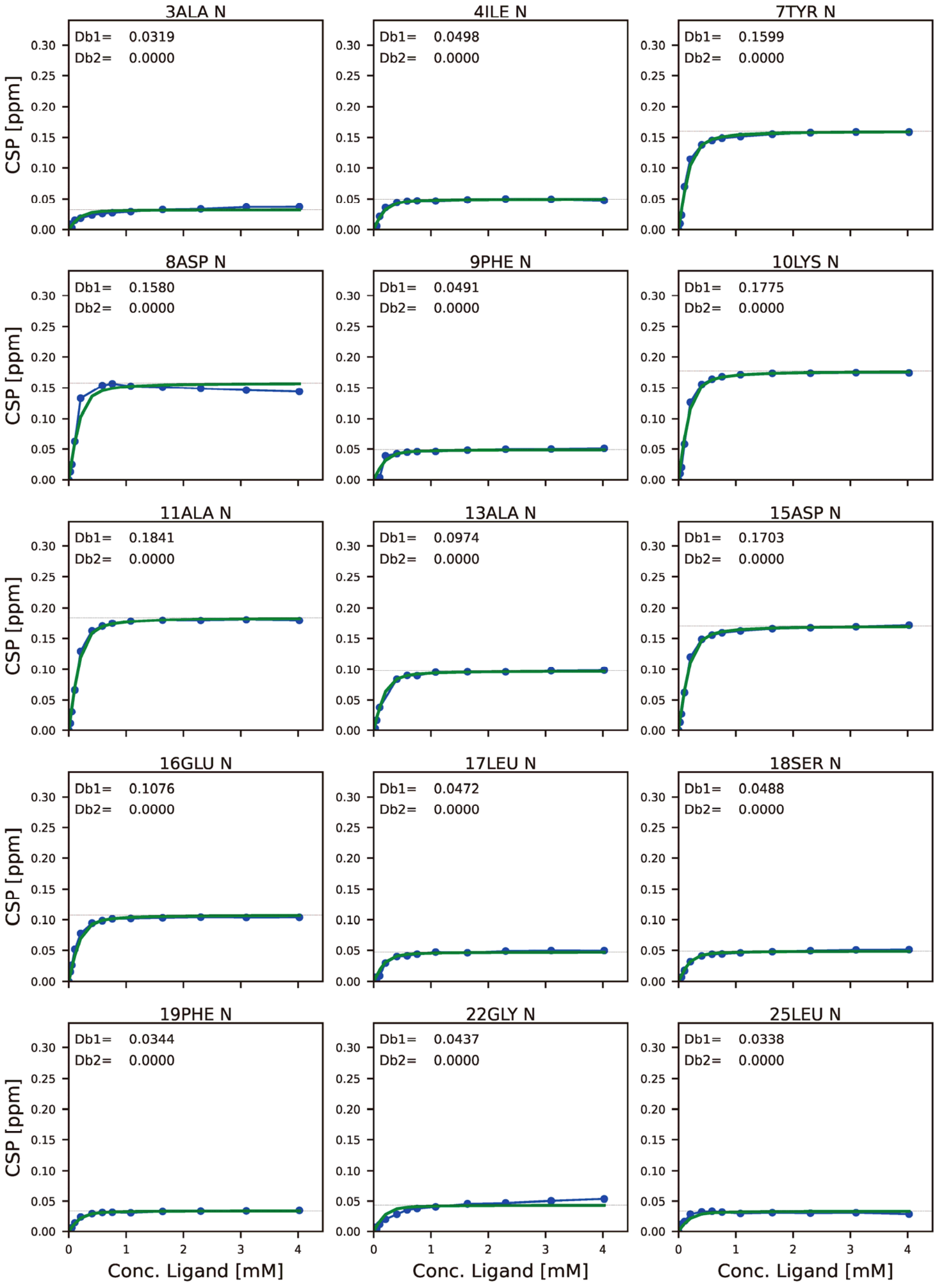
3ALA N
Db1= 0.0319
Db2= 0.0000
4ILE N
Db1= 0.0498
Db2= 0.0000
7TYR N
Db1= 0.1599
Db2= 0.0000
8ASP N
Db1= 0.1580
Db2= 0.0000
9PHE N
Db1= 0.0491
Db2= 0.0000
10LYS N
Db1= 0.1775
Db2= 0.0000
11ALA N
Db1= 0.1841
Db2= 0.0000
13ALA N
Db1= 0.0974
Db2= 0.0000
15ASP N
Db1= 0.1703
Db2= 0.0000
16GLU N
Db1= 0.1076
Db2= 0.0000
17LEU N
Db1= 0.0472
Db2= 0.0000
18SER N
Db1= 0.0488
Db2= 0.0000
19PHE N
Db1= 0.0344
Db2= 0.0000
22GLY N
Db1= 0.0437
Db2= 0.0000
25LEU N
Db1= 0.0338
Db2= 0.0000
CSP [ppm]
Conc. Ligand [mM]

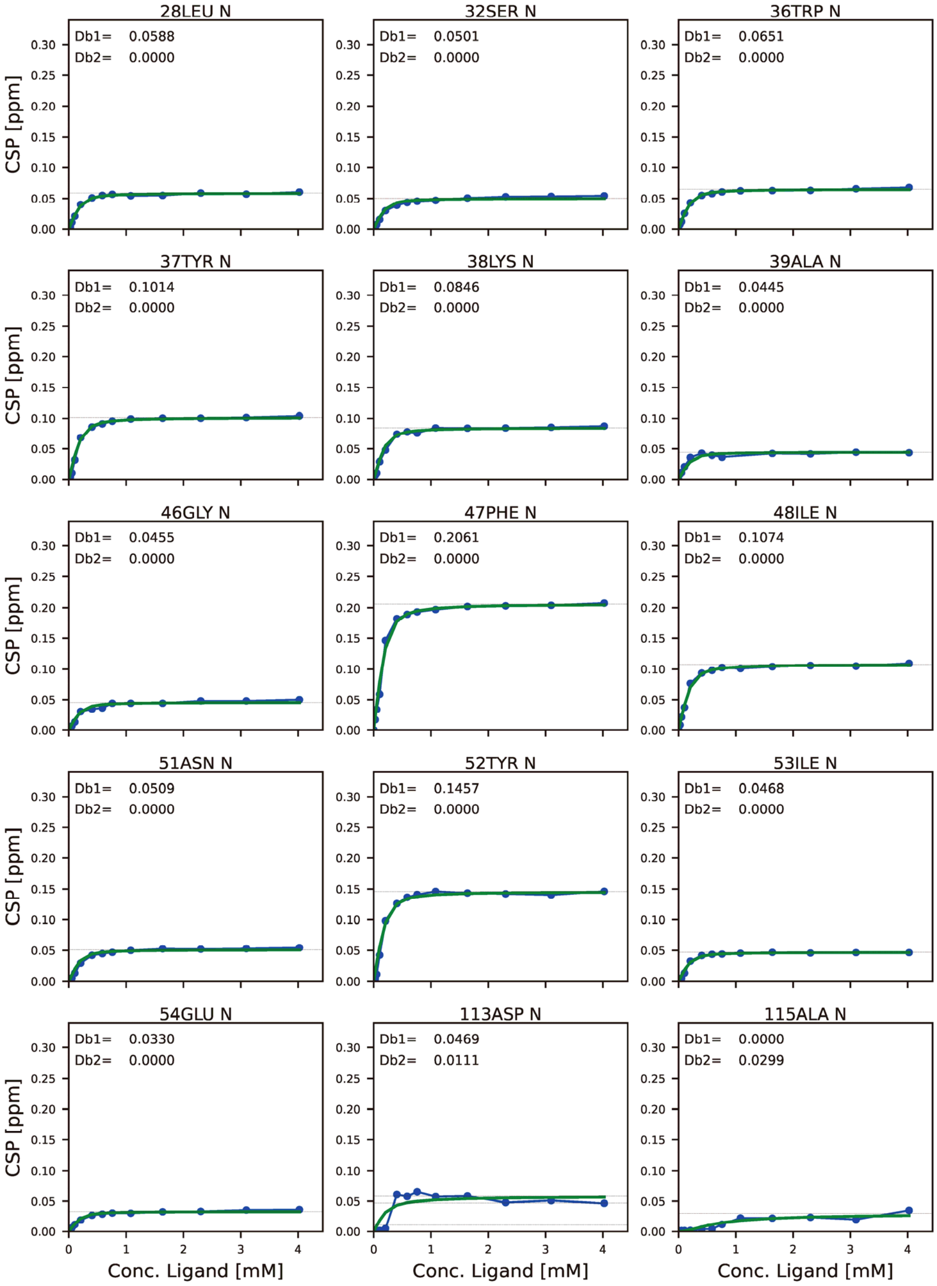

28LEU N
Db1= 0.0588
Db2= 0.0000
32SER N
Db1= 0.0501
Db2= 0.0000
36TRP N
Db1= 0.0651
Db2= 0.0000
37TYR N
Db1= 0.1014
Db2= 0.0000
38LYS N
Db1= 0.0846
Db2= 0.0000
39ALA N
Db1= 0.0445
Db2= 0.0000
46GLY N
Db1= 0.0455
Db2= 0.0000
47PHE N
Db1= 0.2061
Db2= 0.0000
48ILE N
Db1= 0.1074
Db2= 0.0000
51ASN N
Db1= 0.0509
Db2= 0.0000
52TYR N
Db1= 0.1457
Db2= 0.0000
53ILE N
Db1= 0.0468
Db2= 0.0000
54GLU N
Db1= 0.0330
Db2= 0.0000
113ASP N
Db1= 0.0469
Db2= 0.0111
115ALA N
Db1= 0.0000
Db2= 0.0299
CSP [ppm]
Conc. Ligand [mM]

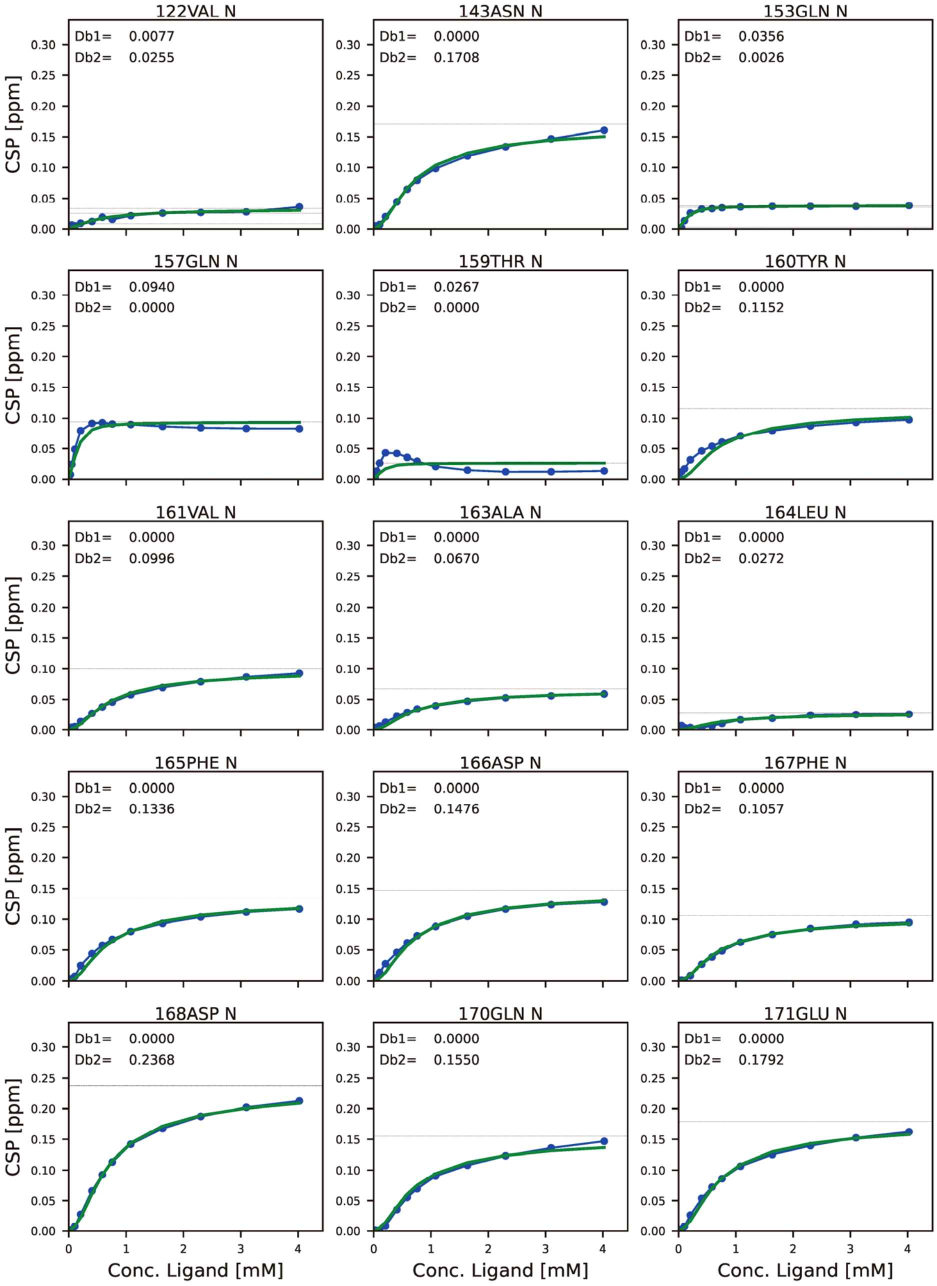
122VAL N
Db1= 0.0077
Db2= 0.0255
143ASN N
Db1= 0.0000
Db2= 0.1708
153GLN N
Db1= 0.0356
Db2= 0.0026
157GLN N
Db1= 0.0940
Db2= 0.0000
159THR N
Db1= 0.0267
Db2= 0.0000
160TYR N
Db1= 0.0000
Db2= 0.1152
161VAL N
Db1= 0.0000
Db2= 0.0996
163ALA N
Db1= 0.0000
Db2= 0.0670
164LEU N
Db1= 0.0000
Db2= 0.0272
165PHE N
Db1= 0.0000
Db2= 0.1336
166ASP N
Db1= 0.0000
Db2= 0.1476
167PHE N
Db1= 0.0000
Db2= 0.1057
168ASP N
Db1= 0.0000
Db2= 0.2368
170GLN N
Db1= 0.0000
Db2= 0.1550
171GLU N
Db1= 0.0000
Db2= 0.1792
CSP [ppm]
Conc. Ligand [mM]

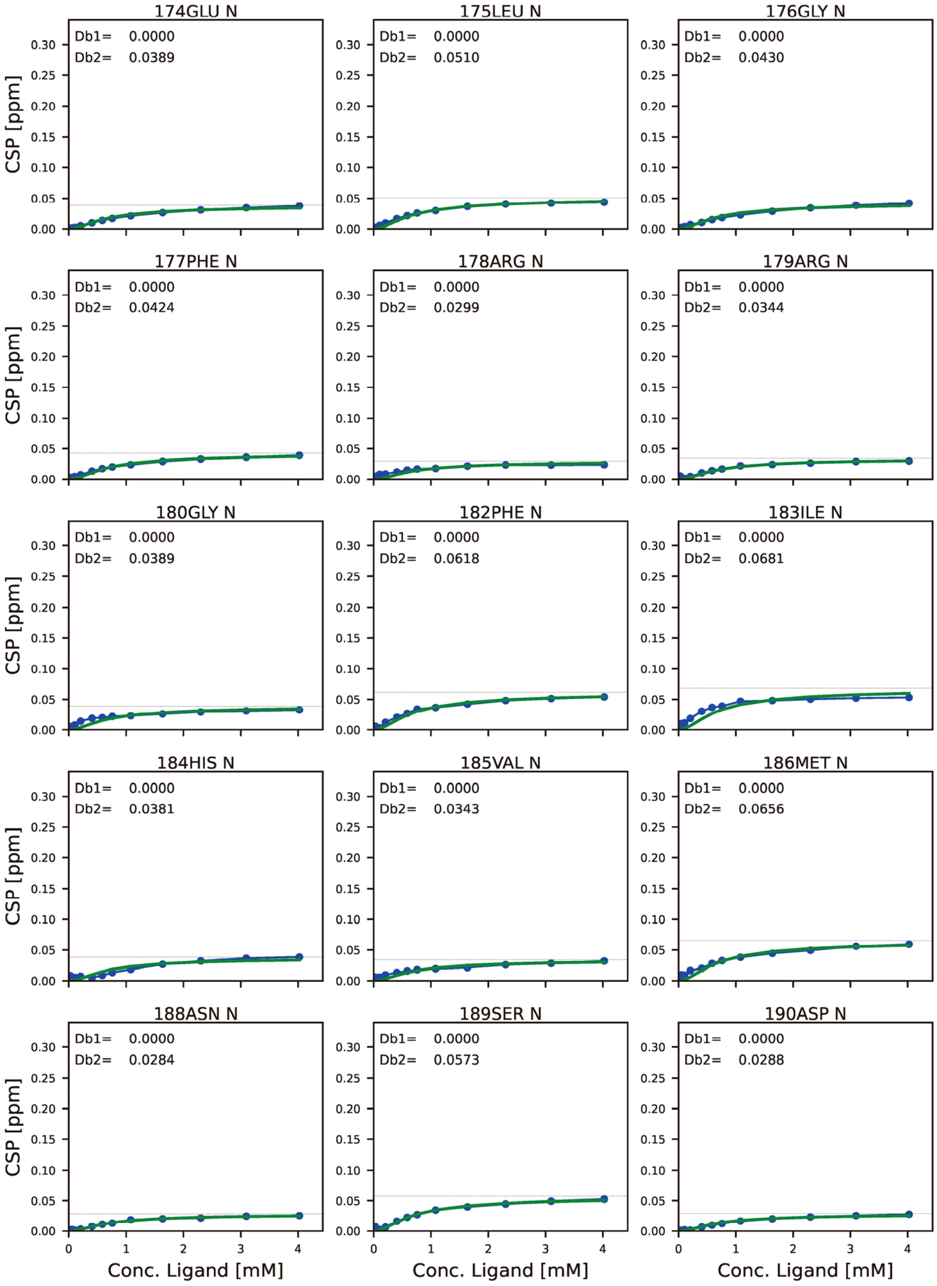
174GLU N
Db1= 0.0000
Db2= 0.0389
175LEU N
Db1= 0.0000
Db2= 0.0510
176GLY N
Db1= 0.0000
Db2= 0.0430
177PHE N
Db1= 0.0000
Db2= 0.0424
178ARG N
Db1= 0.0000
Db2= 0.0299
179ARG N
Db1= 0.0000
Db2= 0.0344
180GLY N
Db1= 0.0000
Db2= 0.0389
182PHE N
Db1= 0.0000
Db2= 0.0618
183ILE N
Db1= 0.0000
Db2= 0.0681
184HIS N
Db1= 0.0000
Db2= 0.0381
185VAL N
Db1= 0.0000
Db2= 0.0343
186MET N
Db1= 0.0000
Db2= 0.0656
188ASN N
Db1= 0.0000
Db2= 0.0284
189SER N
Db1= 0.0000
Db2= 0.0573
190ASP N
Db1= 0.0000
Db2= 0.0288
CSP [ppm]
Conc. Ligand [mM]

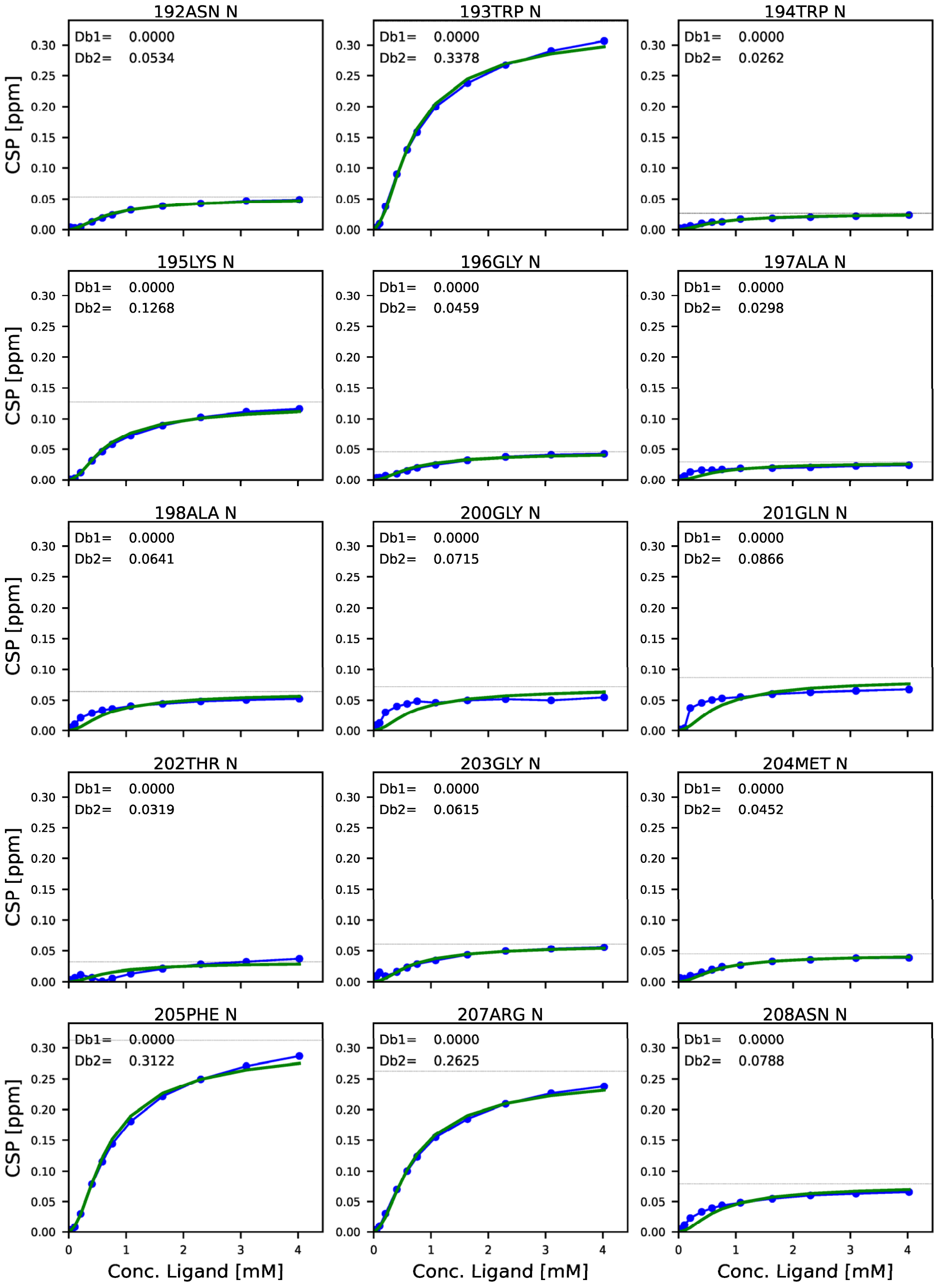
192ASN N
Db1= 0.0000
Db2= 0.0534
193TRP N
Db1= 0.0000
Db2= 0.3378
194TRP N
Db1= 0.0000
Db2= 0.0262
195LYS N
Db1= 0.0000
Db2= 0.1268
196GLY N
Db1= 0.0000
Db2= 0.0459
197ALA N
Db1= 0.0000
Db2= 0.0298
198ALA N
Db1= 0.0000
Db2= 0.0641
200GLY N
Db1= 0.0000
Db2= 0.0715
201GLN N
Db1= 0.0000
Db2= 0.0866
202THR N
Db1= 0.0000
Db2= 0.0319
203GLY N
Db1= 0.0000
Db2= 0.0615
204MET N
Db1= 0.0000
Db2= 0.0452
205PHE N
Db1= 0.0000
Db2= 0.3122
207ARG N
Db1= 0.0000
Db2= 0.2625
208ASN N
Db1= 0.0000
Db2= 0.0788
CSP [ppm]
Conc. Ligand [mM]

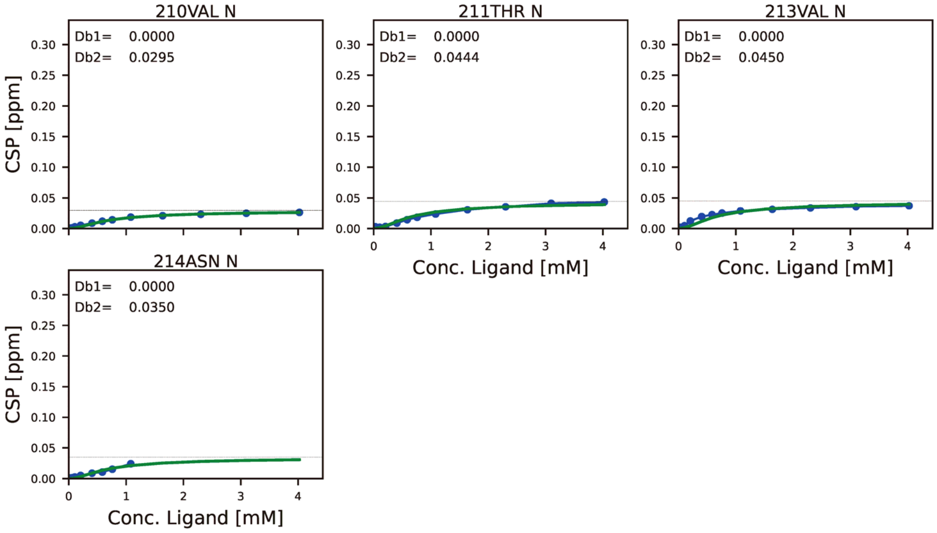


**Figure S13. Chemical shift perturbation profile mapped on each residue, against the concentration of ligand with SOS S9 PRM.**

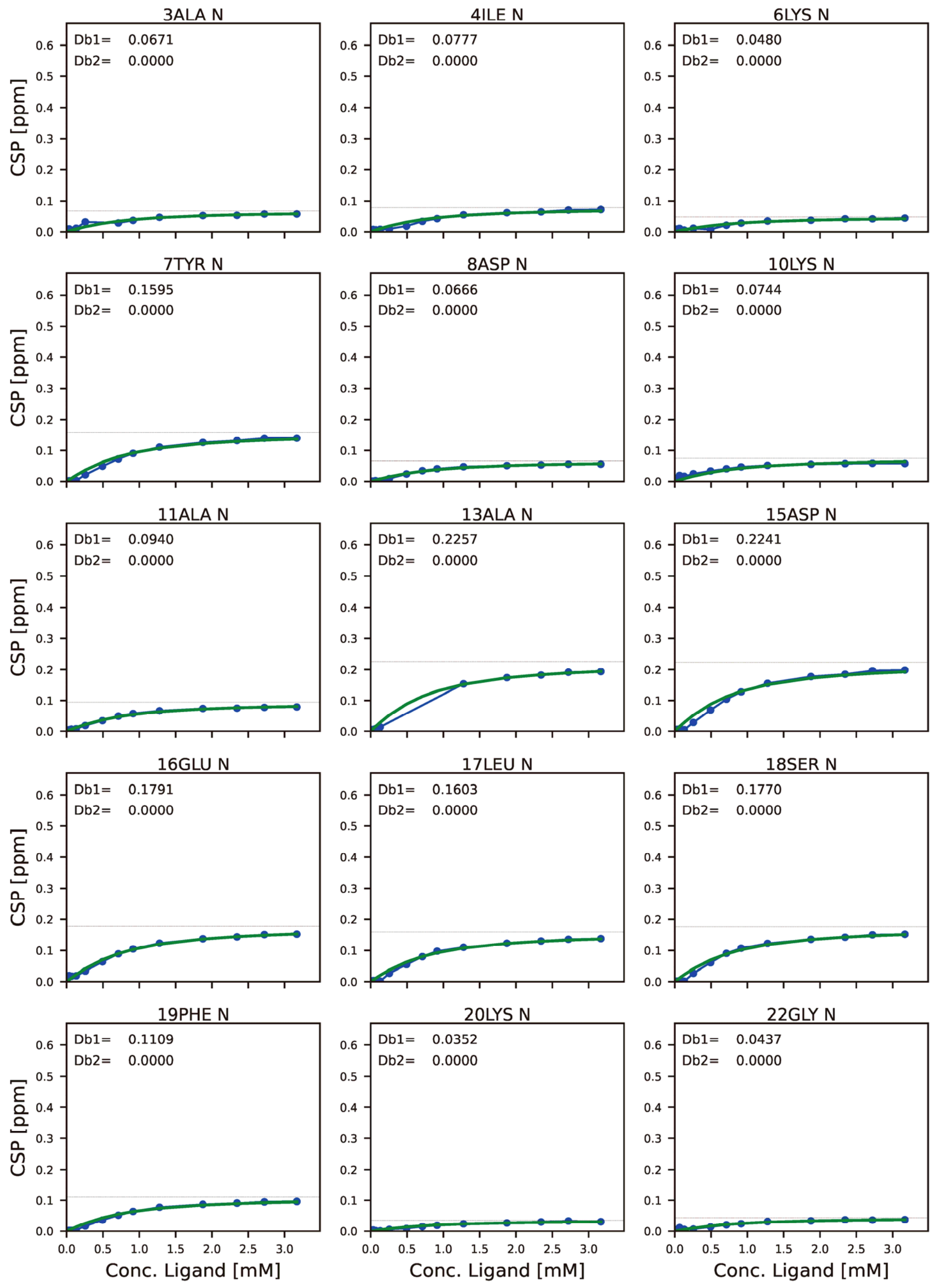
3ALA N
Db1= 0.0671
Db2= 0.0000
4ILE N
Db1= 0.0777
Db2= 0.0000
6LYS N
Db1= 0.0480
Db2= 0.0000
7TYR N
Db1= 0.1595
Db2= 0.0000
8ASP N
Db1= 0.0666
Db2= 0.0000
10LYS N
Db1= 0.0744
Db2= 0.0000
11ALA N
Db1= 0.0940
Db2= 0.0000
13ALA N
Db1= 0.2257
Db2= 0.0000
15ASP N
Db1= 0.2241
Db2= 0.0000
16GLU N
Db1= 0.1791
Db2= 0.0000
17LEU N
Db1= 0.1603
Db2= 0.0000
18SER N
Db1= 0.1770
Db2= 0.0000
19PHE N
Db1= 0.1109
Db2= 0.0000
20LYS N
Db1= 0.0352
Db2= 0.0000
22GLY N
Db1= 0.0437
Db2= 0.0000
CSP [ppm]
Conc. Ligand [mM]

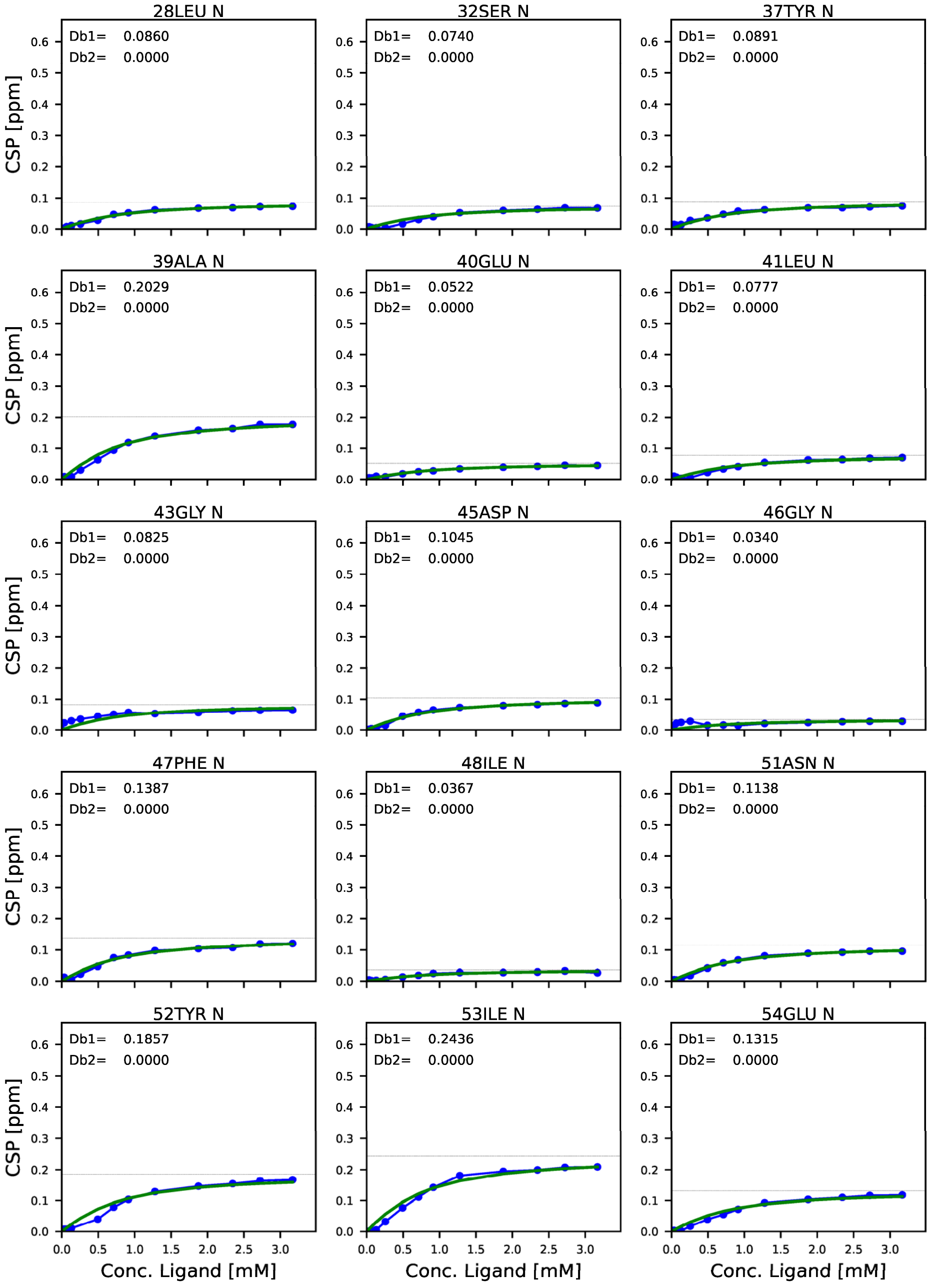
28LEU N
Db1= 0.0860
Db2= 0.0000
32SER N
Db1= 0.0740
Db2= 0.0000
37TYR N
Db1= 0.0891
Db2= 0.0000
39ALA N
Db1= 0.2029
Db2= 0.0000
40GLU N
Db1= 0.0522
Db2= 0.0000
41LEU N
Db1= 0.0777
Db2= 0.0000
43GLY N
Db1= 0.0825
Db2= 0.0000
45ASP N
Db1= 0.1045
Db2= 0.0000
46GLY N
Db1= 0.0340
Db2= 0.0000
47PHE N
Db1= 0.1387
Db2= 0.0000
48ILE N
Db1= 0.0367
Db2= 0.0000
51ASN N
Db1= 0.1138
Db2= 0.0000
52TYR N
Db1= 0.1857
Db2= 0.0000
53ILE N
Db1= 0.2436
Db2= 0.0000
54GLU N
Db1= 0.1315
Db2= 0.0000
CSP [ppm]
Conc. Ligand [mM]

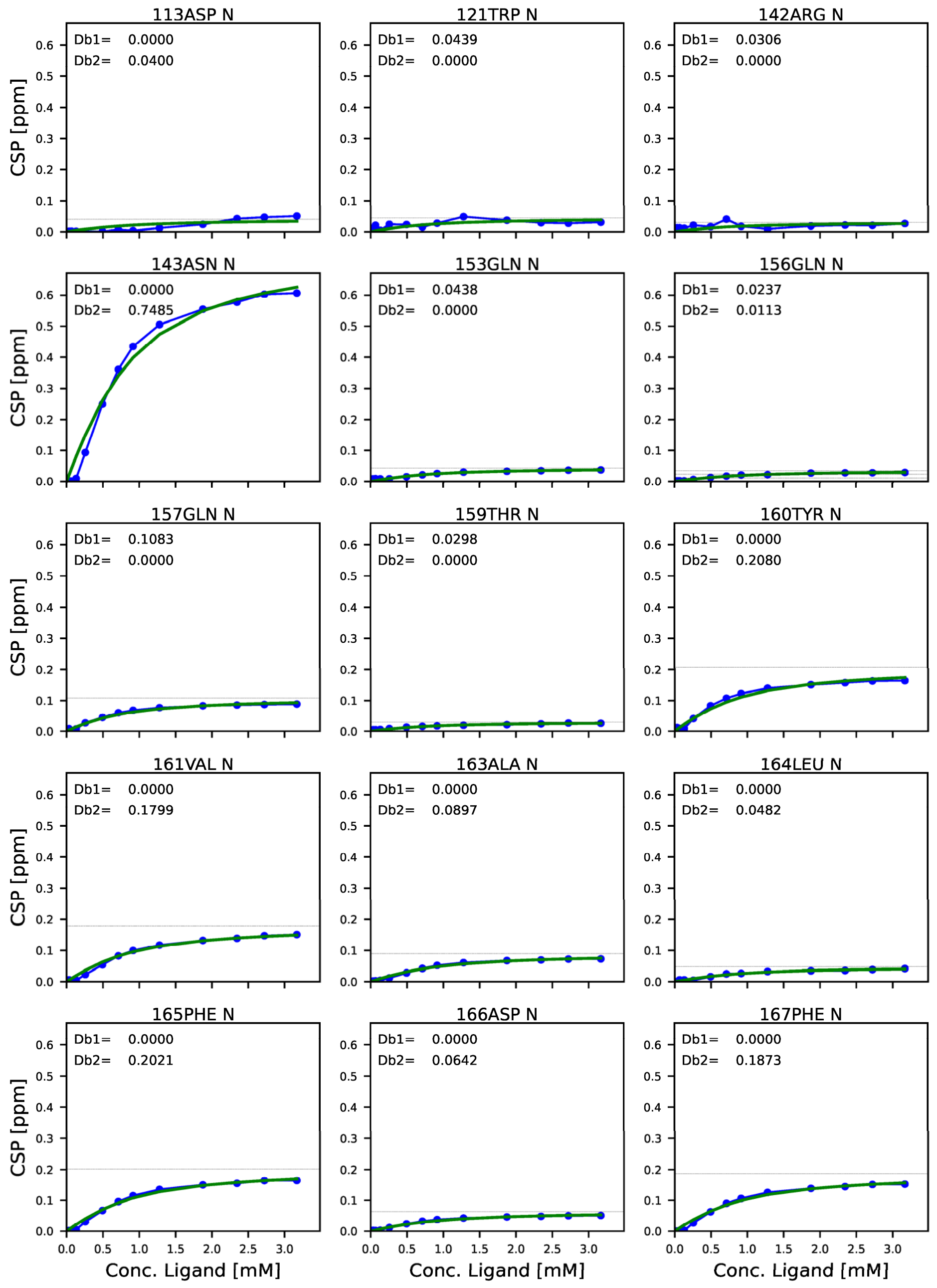
113ASP N
Db1= 0.0000
Db2= 0.0400
121TRP N
Db1= 0.0439
Db2= 0.0000
142ARG N
Db1= 0.0306
Db2= 0.0000
143ASN N
Db1= 0.0000
Db2= 0.7485
153GLN N
Db1= 0.0438
Db2= 0.0000
156GLN N
Db1= 0.0237
Db2= 0.0113
157GLN N
Db1= 0.1083
Db2= 0.0000
159THR N
Db1= 0.0298
Db2= 0.0000
160TYR N
Db1= 0.0000
Db2= 0.2080
161VAL N
Db1= 0.0000
Db2= 0.1799
163ALA N
Db1= 0.0000
Db2= 0.0897
164LEU N
Db1= 0.0000
Db2= 0.0482
165PHE N
Db1= 0.0000
Db2= 0.2021
166ASP N
Db1= 0.0000
Db2= 0.0642
167PHE N
Db1= 0.0000
Db2= 0.1873
CSP [ppm]
Conc. Ligand [mM]

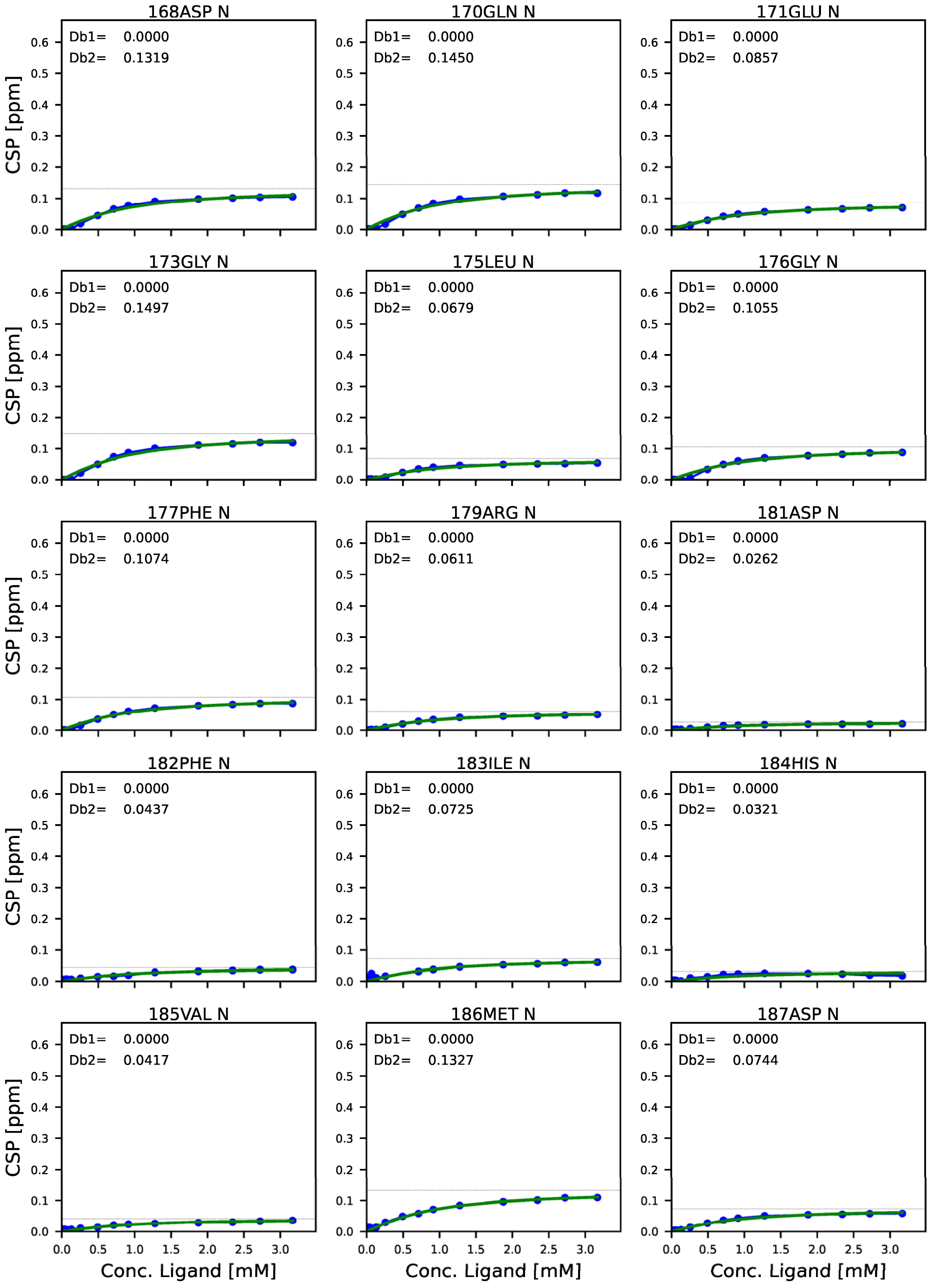
168ASP N
Db1= 0.0000
Db2= 0.1319
170GLN N
Db1= 0.0000
Db2= 0.1450
171GLU N
Db1= 0.0000
Db2= 0.0857
173GLY N
Db1= 0.0000
Db2= 0.1497
175LEU N
Db1= 0.0000
Db2= 0.0679
176GLY N
Db1= 0.0000
Db2= 0.1055
177PHE N
Db1= 0.0000
Db2= 0.1074
179ARG N
Db1= 0.0000
Db2= 0.0611
181ASP N
Db1= 0.0000
Db2= 0.0262
182PHE N
Db1= 0.0000
Db2= 0.0437
183ILE N
Db1= 0.0000
Db2= 0.0725
184HIS N
Db1= 0.0000
Db2= 0.0321
185VAL N
Db1= 0.0000
Db2= 0.0417
186MET N
Db1= 0.0000
Db2= 0.1327
187ASP N
Db1= 0.0000
Db2= 0.0744
CSP [ppm]
Conc. Ligand [mM]

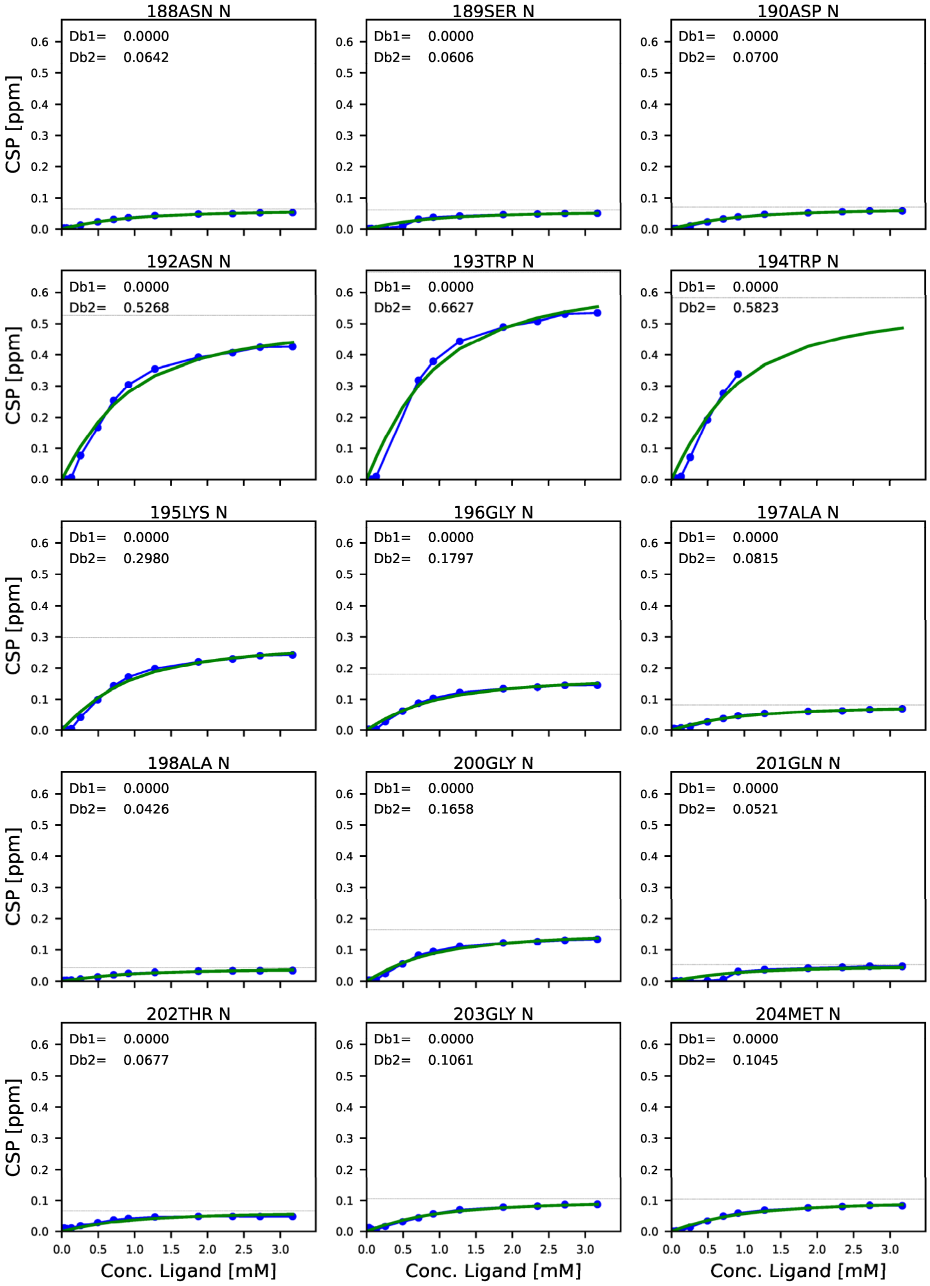
188ASN N
Db1= 0.0000
Db2= 0.0642
189SER N
Db1= 0.0000
Db2= 0.0606
190ASP N
Db1= 0.0000
Db2= 0.0700
192ASN N
Db1= 0.0000
Db2= 0.5268
193TRP N
Db1= 0.0000
Db2= 0.6627
194TRP N
Db1= 0.0000
Db2= 0.5823
195LYS N
Db1= 0.0000
Db2= 0.2980
196GLY N
Db1= 0.0000
Db2= 0.1797
197ALA N
Db1= 0.0000
Db2= 0.0815
198ALA N
Db1= 0.0000
Db2= 0.0426
200GLY N
Db1= 0.0000
Db2= 0.1658
201GLN N
Db1= 0.0000
Db2= 0.0521
202THR N
Db1= 0.0000
Db2= 0.0677
203GLY N
Db1= 0.0000
Db2= 0.1061
204MET N
Db1= 0.0000
Db2= 0.1045
CSP [ppm]
Conc. Ligand [mM]

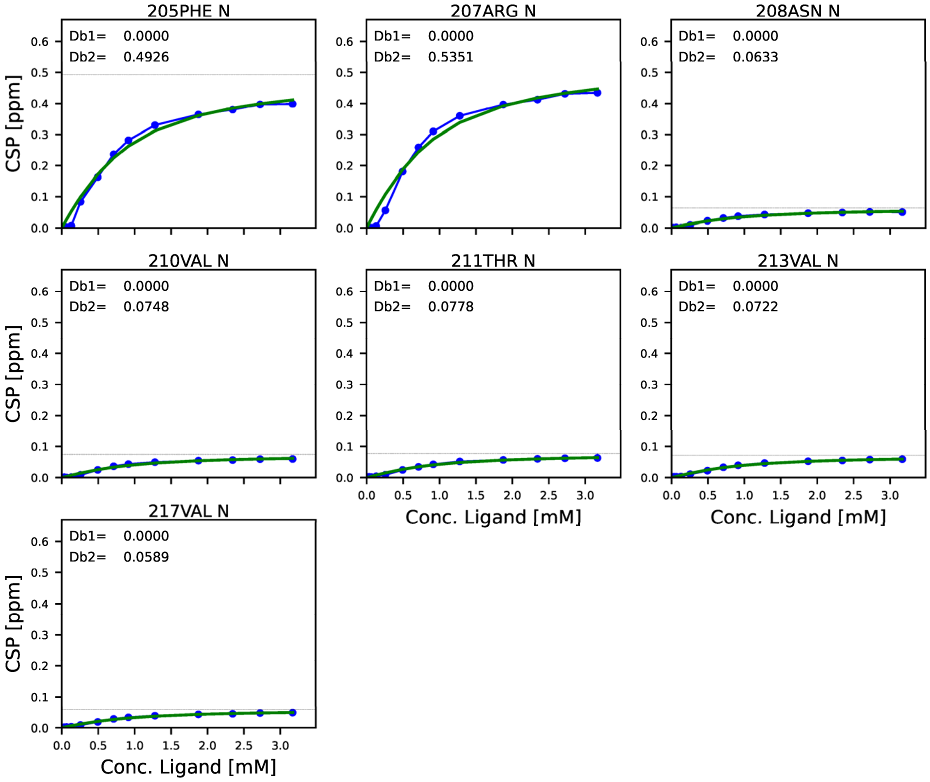


**Figure S14. Chemical shift perturbation profile mapped on each residue, against the concentration of ligand with SOS S10 PRM.**

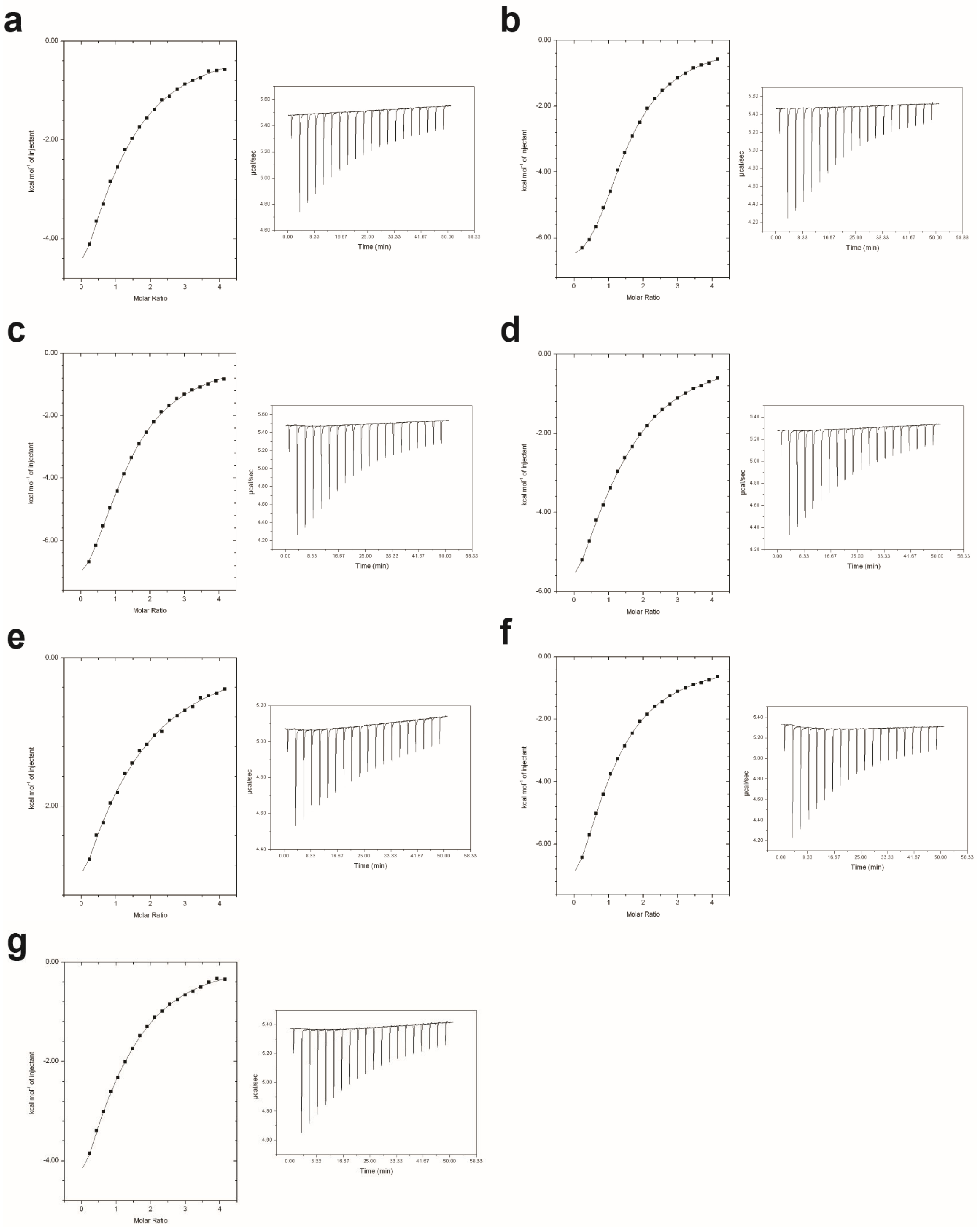


**Figure S15. ITC analysis for the binding of the full-length GRB2 to PRMs.** Figures a-g indicate ITC analysis for S2 (**a**), S4 (**b**), S5 (**c**), S6 (**d**), S8 (**e**) S9 (**f**), and S10 (**g**) PRMs, respectively. Data with S1, S3, and S7 are not shown here because of no clear thermogram curves. The solid lines show the fit of the data to a "Two Set of Sites" model based on the binding of a ligand to a macromolecule, as incorporated in the Origin software.

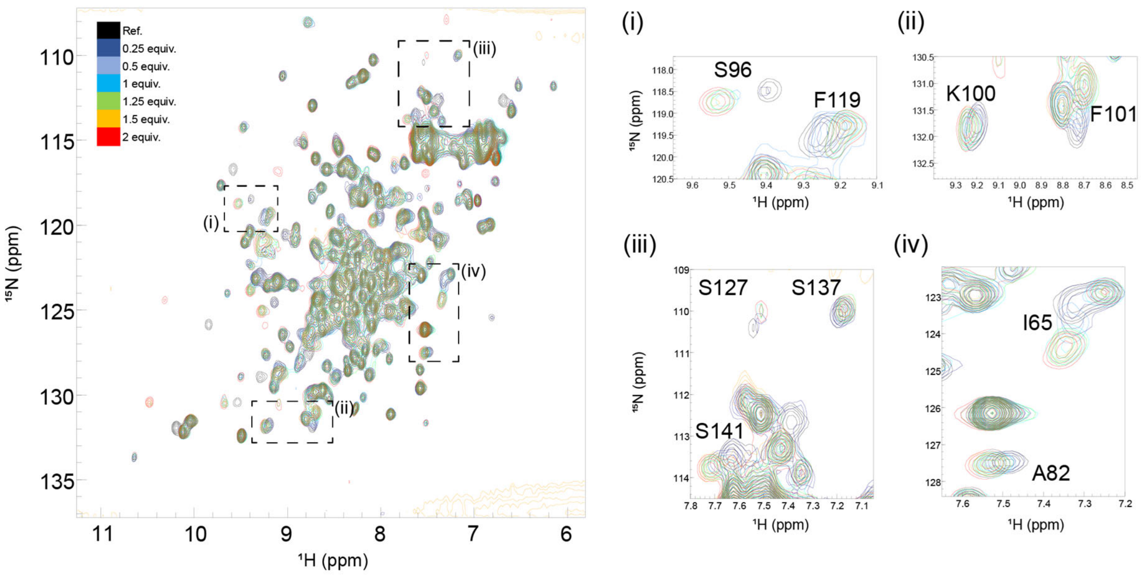


**Figure S16. Overlays of 2D $^{1}H$-$^{15}N$ HSQC spectra from multipoint titrations of $^{15}N$-labelled GRB2 exclusively with the EGFR phosphorylated peptide (EpYINSQV).** The peptide concentration was increased stepwise (the protein: peptide molar ratio of 1:0.25, 1:0.5, 1:1, 1:1.25, 1:1.5, and 1:2). In this figure, the colour codes of $^{1}H$-$^{15}N$ correlation cross-peaks at each titration point, showing the molar ratio of GRB2: EGFR, are as follows: black (1:0); green (1:0.25); blue (1:0.5); yellow (1:0.75); magenta (1:1); cyan (1:1.25); brown (1:1.5); coral (1:2); red (1:4). Cross peaks that showed large chemical shift changes were annotated.

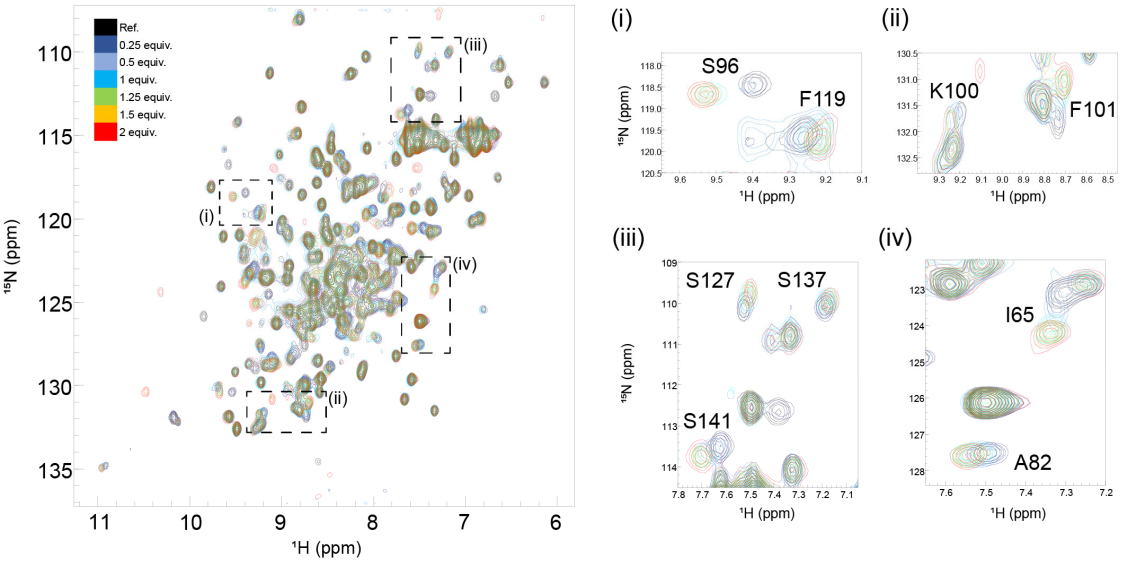


**Figure S17. Overlays of 2D $^{1}H$-$^{15}N$ HSQC spectra from multipoint titrations of $^{15}N$-labelled GRB2 with the EGFR phosphorylated peptide (EpYINSQV), in the presence of S4 PRM.** The peptide concentration was increased stepwise (the protein: peptide molar ratio of 1:0.25, 1:0.5, 1:1, 1:1.25, 1:1.5, and 1:2). In this figure, the colour codes of $^{1}H$-$^{15}N$ correlation cross-peaks at each titration point, showing the molar ratio of GRB2: EGFR, are as follows: black (1:0); green (1:0.25); blue (1:0.5); yellow (1:0.75); magenta (1:1); cyan (1:1.25); brown (1:1.5); coral (1:2); red (1:4). Cross peaks that showed large chemical shift changes were annotated.

**Table S1. Dissociation constants derived from ITC analysis for the full length GRB3 and isolated SH3 domains against SOS1 PRMs**

| PRM | Sequence | ITC analysis | | | |
|---|---|---|---|---|---|
| | | $K_D$ (μM) from the full length GRB2[a] | | $K_D$ (μM) from isolated SH3[b] | |
| | | $K_{D1}$ | $K_{D2}$ | $K_{D1}$ | $K_{D2}$ |
| S1 | VPPPVPPRRR (1148-1159) | NA | NA | — | — |
| S2 | VPPPVPPRRR (1148-1159) | 65.1 ± 23.7 | NA | — | — |
| S3 | VPPPVPPRRR (1148-1159) | NA | NA | — | — |
| S4 | VPPPVPPRRR (1148-1159) | 5.3 ± 0.4 | 51.3 ± 1.7 | 39 ± 1 | 125 ± 13 |
| S5 | DSPPAOPPRQPT (1177-1188) | 15.9 ± 1.5 | 109.7 ± 6.5 | 56 ± 5 | 1396 ± 87 |
| S6 | DSPPAOPPRQPT (1177-1188) | 40.1 ± 6.5 | 97.3 ± 10.5 | 117 ± 2 | 1718 ± 33 |
| S7 | DSPPAOPPRQPT (1177-1188) | NA | NA | — | — |
| S8 | DSPPAOPPRQPT (1177-1188) | 122.9 ± 18.6 | NA | — | — |
| S9 | IAGPPVPPRQST (1287-1298) | 35.6 ± 4.2 | 640.0 ± 291.1 | 82 ± 1 | 1318 ± 44 |
| S10 | PKLPPKTYKREH (1303-1314) | 71.5 ± 24.2 | 130.8 ± 54.4 | — | — |

[a]The dissociation constants were obtained by globally fitting ITC data to the "Two Set of Sites" model using MicroCal ITC-ORIGIN Analysis Software (Malvern Panalytical). [b]The dissociation constants are published data from ITC in the previous work using isolated SH3 domains{McDonald, 2009 #67}.
